# Rare mutations implicate CGE interneurons as a vulnerable axis of cognitive deficits across psychiatric disorders

**DOI:** 10.1101/2025.03.28.645799

**Authors:** Stephanie A. Herrlinger, Jiayao Wang, Bovey Y. Rao, Erica Rodriguez, Jonathan Chang, Joseph A. Gogos, Attila Losonczy, Dennis Vitkup

**Author notes:** Correspondence should be addressed to JG, AL and DV. These authors contributed equally to this work (Stephanie A. Herrlinger, Jiayao Wang).

## Abstract

Neuropsychiatric disorders such as autism spectrum disorder (ASD) and schizophrenia (SCZ) share genetic risk factors, including genes affected by rare high-penetrance single nucleotide variants (SNVs) and copy number variants (CNVs). ASD and SCZ exhibit both overlapping and distinct clinical phenotypes. Cognitive deficits and intellectual disability—critical predictors of long-term outcomes—are common to both conditions. To investigate shared and disorder-specific neurobiological impact of highly penetrant rare mutations in ASD and SCZ, we analyzed human single-nucleus whole-brain sequencing data to identify strongly affected brain cell types. Our analysis revealed caudal ganglionic eminence (CGE)-derived GABAergic interneurons as a key nexus for cognitive deficits across these disorders. Notably, genes within 22q11.2 deletions, known to confer a high risk for SCZ, ASD, and cognitive impairment, showed a strong expression bias toward vasoactive intestinal peptide-expressing cells (VIP+) among CGE subtypes. To explore perturbations of VIP+ GABAergic interneurons in the 22q11.2 deletion syndrome, we examined both their cell intrinsic properties with slice electrophysiology and *in vivo* activity in the *Df(16)A^+/-^*mouse model during a spatial navigation task and observed reduced firing and activity along with altered responses to ambulation and random rewards. At the population level, VIP+ interneurons exhibited impaired spatial encoding and diminished subtype-specific activity suggesting deficient disinhibition in CA1 microcircuits in the hippocampus, a region essential for learning and memory. Overall, these results demonstrate the crucial role of CGE-derived interneurons in mediating cognitive processes that are disrupted across a range of psychiatric and neurodevelopmental disorders.

**Graphical Abstract:** **Left:** Integration of genetic and single-cell transcriptomic data was used to identify cell type-specific disorder biases. Genes implicated in ASD and SCZ^1,2^ and genes in the 22q11.2 deletion were used to calculate cell type mutation biases based on human brain snRNA-seq data from 461 cell types^37^. **Middle:** Disorder cell type bias analysis revealed shared cell type risk patterns between distinct disorders with overlapping cognitive phenotypes. **Right:** Experimental analysis of the 22q11.2 deletion mouse model was used to identify cell type specific deficits *in vivo* using 2-photon calcium imaging to capture neuronal activity during a spatial navigation task.

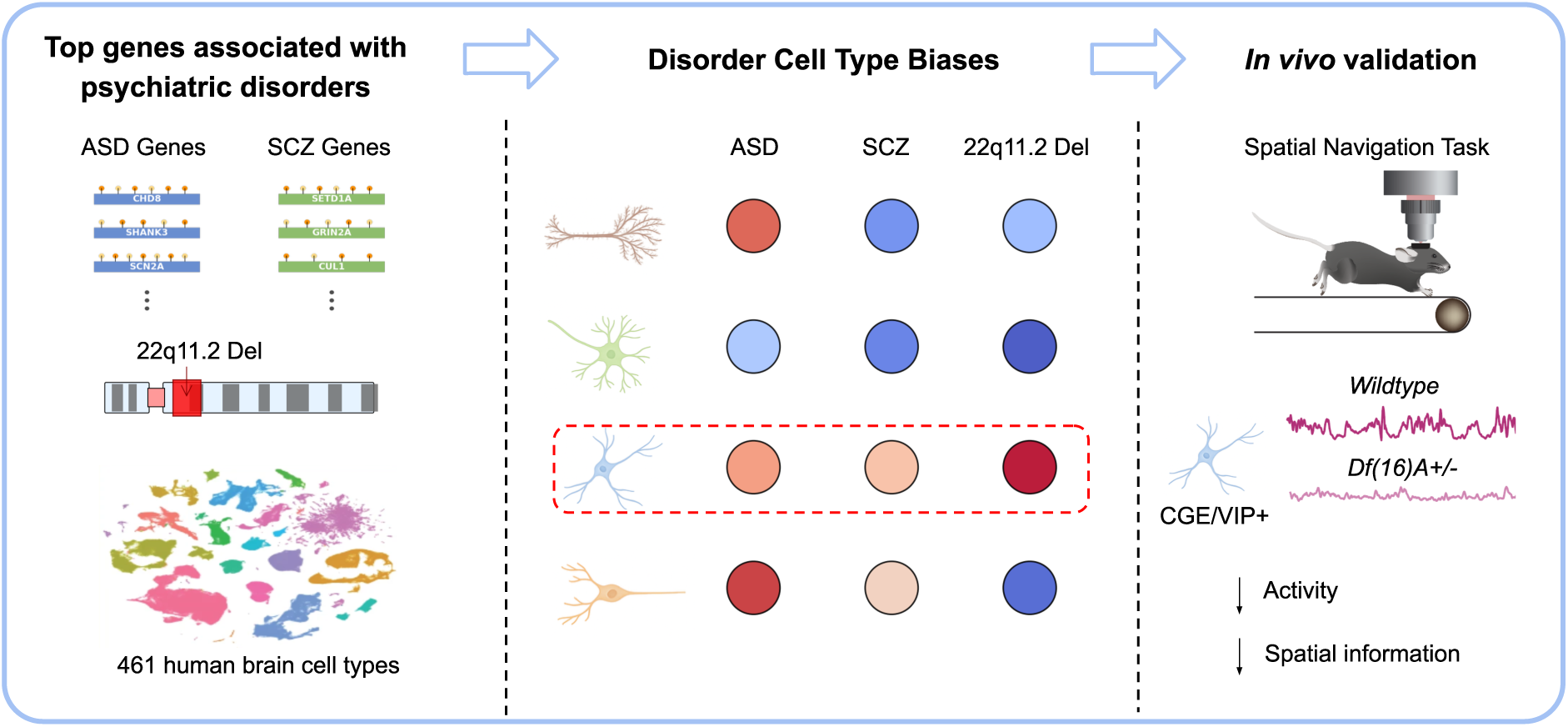

## Introduction

Psychiatric disorders exhibit distinct clinical symptoms, yet frequently share genetic risk factors and underlying cellular and molecular mechanisms^1–6^. For example, schizophrenia (SCZ) is characterized by psychosis, including hallucinations and delusions, as well as cognitive control impairments that contribute to disorganization, impulsivity, and difficulties with working memory^7–11^. In contrast, autism spectrum disorder (ASD) is characterized by restricted interests, difficulties with social communication, and cognitive rigidity along with repetitive behaviors^8,12^. Several copy number variants (CNVs), such as the 22q11.2 deletion, have been implicated in both SCZ and ASD, as well as in other psychiatric disorders^13,14^. Among their diverse symptoms, cognitive impairments and intellectual disability (ID) are frequently comorbid with neuropsychiatric phenotypes^15^ and are especially pronounced in individuals carrying highly penetrant genetic variants^15–18^. Cognitive deficits in psychiatric disorders are particularly debilitating, and span a broad range of functions, including memory, attention, reasoning, decision-making, and social cognition^19^. The severity of these deficits is a strong predictor of patients’ long-term health outcomes and overall well-being^20–22^.

Despite advances in the genetics of disorders such as ASD and SCZ^1,2,23^, therapeutic progress in treating their cognitive symptoms remains limited, largely because of gaps in understanding the neurobiological mechanisms through which genetic risk factors affect cognition. To better understand the relationships between genetics and psychiatric phenotypes, researchers are increasingly integrating genetic analyses with single-cell and single-nucleus sequencing data to identify the cell types affected across psychiatric disorders^24–26^. Animal models, particularly mouse models that recapitulate rare, high-risk genetic alterations, play a critical role in this integrative approach. The evolutionary conservation of brain regions, cell types, and core biological processes between mice and humans^27–30^ facilitates a detailed investigation of genetic risk factors at both the cellular and circuit levels^31–36^.

In this study, we investigated the neurobiological effects of rare, highly penetrant mutations associated with ASD, SCZ, and 22q11.2 deletion syndrome across diverse brain cell types, using recently released human whole-brain single-nucleus RNA sequencing data^37^. Our analyses revealed distinct disorder-specific mutation biases affecting different cell types, with caudal ganglionic eminence (CGE)-derived interneurons consistently implicated across genetic insults associated with cognitive dysfunction, including ASD with intellectual disability, SCZ, and developmental disorders and intellectual disability (DD/ID). Furthermore, our analysis of genes located within the 22q11.2 deletion region revealed that VIP+ CGE-derived interneurons were strongly affected by this genetic alteration. Using *in vivo* two-photon calcium imaging of VIP+ GABAergic interneuron activity in a mouse model of the 22q11.2 deletion, we observed specific alterations in interneuron-selective interneuron (ISI) subtypes, which primarily inhibit other interneurons. These findings implicate CGE/ISI interneuron dysfunction in circuit-level abnormalities contributing to the cognitive and behavioral phenotypes of 22q11.2 deletion syndrome and potentially many other psychiatric conditions.

## Results

### Identifying disorder-specific cell type mutation biases by integrating genetic and single-nucleus expression data

Individually, highly penetrant rare mutations typically have substantially greater functional effects on psychiatric phenotypes than common variants^1,38^. The availability of well-characterized rare variant datasets from ASD and SCZ cohorts enabled us to perform direct comparisons between the two disorders. For ASD, we utilized a recent meta-analysis performed by the SPARK consortium^2^, which included *de novo* mutations from 42,607 ASD probands. This analysis identified 61 genes reaching exome-wide significance based on *de novo* mutation burden. For SCZ, we used a case-control study involving 24,248 SCZ patients and 97,322 control individuals from the SCHEMA consortium^1^. To match the number of considered ASD-associated genes, we selected the top 61 SCZ genes based on the strength of their genetic associations.

A significant fraction of probands in the ASD cohort have IQ scores ≤ 70, a group that we refer to as “ASD with ID”^39,40^ (Figure S1). To facilitate a more appropriate comparison with SCZ, for which the average premorbid IQ is approximately 94^41,42^, we selected ASD cases with IQ scores >70 from the SPARK cohort, a group we refer to as “ASD without ID”, with an average IQ of approximately 95. Notably, individuals with SCZ who carry rare mutations have IQ scores that are only marginally lower than those of the broader SCZ population^43^. Furthermore, genes implicated in ASD without ID exhibit developmental expression profiles that are more similar to those of SCZ genes than to those associated with ASD with ID (Figure S2). It is important to emphasize that although individuals with SCZ and ASD without ID have comparable overall IQ scores, the nature of cognitive dysfunction in these conditions is often distinct. Whereas SCZ is usually characterized by broad deficits across multiple cognitive domains, such as working memory and executive function^44,45^, individuals with ASD without ID tend to show more selective, domain-specific cognitive impairments, particularly in social cognition and communication^46,47^.

To identify the brain cell types most strongly affected by rare coding mutations in ASD and SCZ, we used a recent single-nucleus RNA sequencing dataset collected from the entire human brain^37^. This dataset included samples from approximately 105 anatomical locations dissected from three donors and distributed across the forebrain, midbrain, and hindbrain, with 3,369,219 individual nuclei classified into 31 cell type superclusters, representing broad categories of major brain cell classes, and 461 cell type clusters, representing finer-grained subdivisions based on robust transcriptomic similarity. To evaluate the impact of mutations on brain cell types in ASD and SCZ, we calculated mutation biases across brain cell types by integrating gene expression with disorder-specific mutation burdens (Figure S3 and S4, see Methods). To that end, we first computed a cell type specificity score for each gene, quantifying how selectively the gene is expressed in a given cell type compared with others cell types^26^. Specifically, we first quantified gene expression levels using transcripts per million (TPM) for each cell type and then used these TPM values to compute specificity scores that captured the enrichment of each gene’s expression across cell types. Next, to assess how strongly each cell type was implicated in each disorder, we integrated the expression specificity scores with each gene’s mutation burden, using *de novo* mutation counts for ASD and effective case mutation counts for SCZ (see Methods). The resulting cell type mutation bias captured how selectively disease genes were expressed in each cell type, weighted by their mutational burden in the corresponding patient cohorts. These mutation bias results were robust across a range of gene set sizes (Figure S5) and to cohort downsampling (Figure S6, see Methods). To evaluate the statistical significance of the observed mutation biases, we compared the observed bias for each cell type cluster with a null distribution generated from 10,000 random gene sets matched in size to the disorder-specific gene set, with the original mutation weights randomly reassigned to genes in each null set. To further control for gene-level confounders, we also applied an alternative null model in which random gene sets were matched to the disorder gene set by coding sequence (CDS) length, whole-brain expression level, and evolutionary conservation (Figure S7, see Methods). Empirical P-values were then calculated from these simulations and corrected for multiple testing across the 461 clusters.

### Shared patterns of cell type mutation biases across ASD and SCZ

The cell type superclusters that showed strong mutation biases in ASD without ID included amygdala excitatory neurons, medium spiny neurons, hippocampal dentate gyrus neurons, and deep-layer intratelencephalic (IT) neurons (Figure S8A). The same superclusters also showed strong mutational impacts in ASD with ID, with additional significant cell types including excitatory neurons from the cortex and hippocampus, as well as cortical interneurons (Figure S8B). For SCZ, cell type superclusters with strong mutational impacts included several populations implicated in recent GWAS studies of SCZ^25,26^. Specifically, MGE interneurons were the most strongly impacted cell type, with eccentric medium spiny neurons, Lamp5-LHX6 and Chandelier neurons, CGE interneurons, upper- and deep-layer IT neurons, and midbrain-derived inhibitory neurons also strongly affected (Figure S8C). To confirm that the neuronal cell-type enrichment observed for psychiatric disorders was not an artifact of our computational framework, we applied the same pipeline to non-brain traits from UK Biobank rare-variant burden analyses (HDL cholesterol, inflammatory bowel disease, alanine aminotransferase). Reassuringly, none of these traits showed systematic enrichment in neuronal cell types (Figure S7).

Despite disorder specific differences, several cell type superclusters were strongly impacted in both ASD without ID and SCZ (Figure S8A, Figure S8C) and mutation biases across neuronal cell type clusters were significantly correlated between them (Spearman’s ρ = 0.67, P < 10⁻¹⁰; Figure 1A). This correlation was primarily driven by highly constrained genes contributing to the mutations burdens of both disorders^1,2,39,49,50^ (Figure 1B), rather than by overall gene expression levels in the brain (Figure S9). These shared populations included multiple cortical excitatory classes, as well as excitatory neurons in the amygdala and in hippocampus (Figure 1C). These shared neuronal disruptions may underlie the overlapping dysfunction of brain circuits mediating psychiatric phenotypes^48^. These findings are consistent with several previous studies demonstrating frequent perturbations of shared neuron types across brain disorders. For example, amygdala neurons and their cortical and hippocampal projections have been implicated in a broad range of behavioral disturbances in SCZ and ASD, including deficits in emotion, motivation, and social interactions^51,52^. Moreover, the hippocampus is known to play a key role in social and cognitive deficits in ASD^53^, as well as in learning deficits and positive symptoms associated with SCZ^54,55^. Deep-layer IT neurons, which are central to sensory processing, decision-making, and learning, are also known to be strongly affected in both disorders^56^.

**Figure 1.**
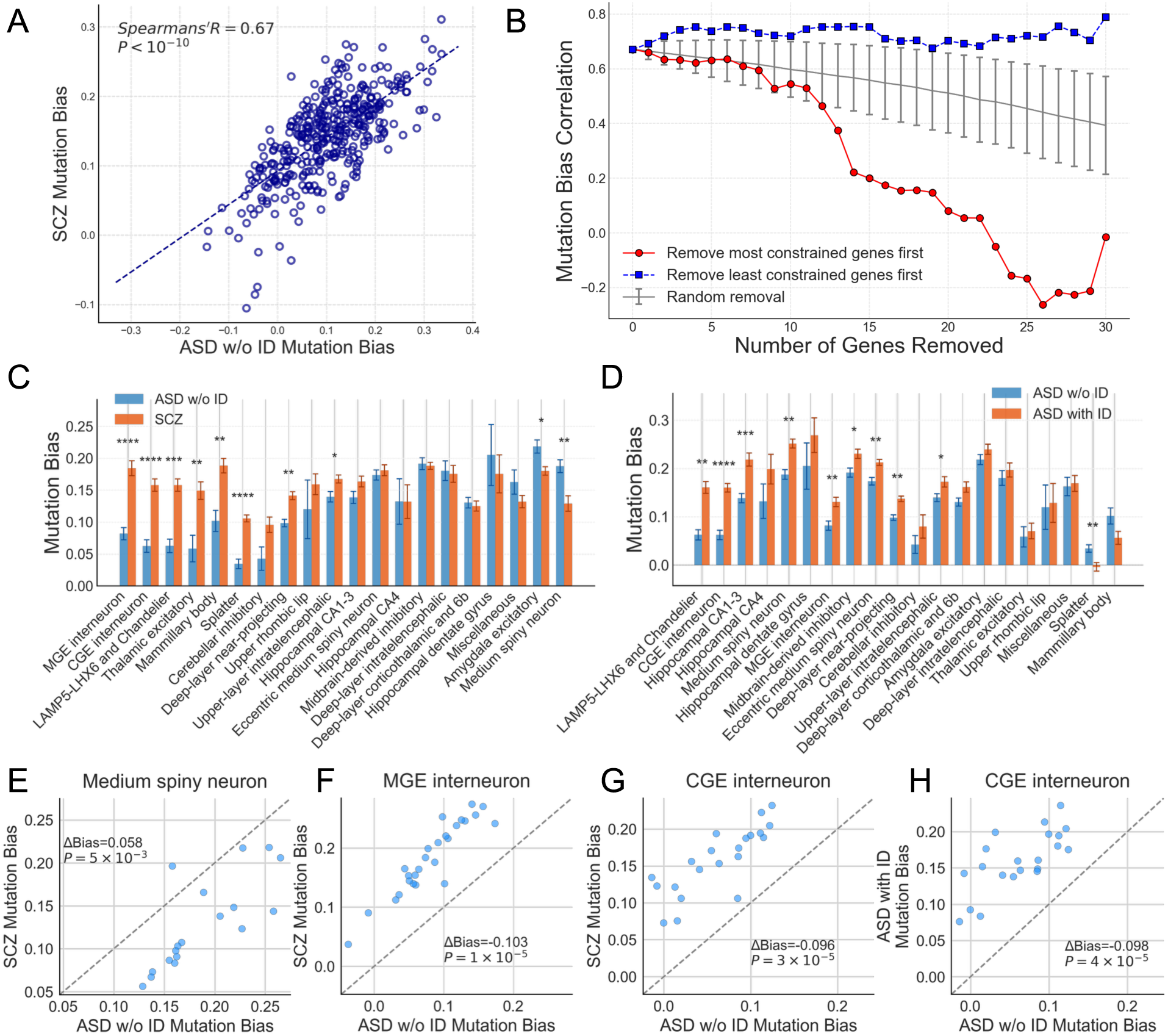
ASD and SCZ Mutation bias towards different brain cell types. **A**. The correlation of mutation biases between ASD w/o ID and SCZ across all neuronal cell types (Spearman’s R = 0.67, P < 10^−10^). Each dot represents a neuron cell type cluster. **B**. Spearman correlation of cell type bias between ASD w/o ID and SCZ genes, assessed by sequential gene removal. Genes were removed one at a time from each disorder’s gene set in parallel, either starting from the most constrained (blue) or least constrained (red) based on LOEUF score^63^. At each step, ASD w/o ID–SCZ bias correlations were recomputed using mutation biases obtained from the remaining genes. The gray line shows the average correlation for random gene removals, with error bars indicating ±1 standard deviation. **C**. Bar plot comparing mutation biases across cell type superclusters between ASD w/o ID (blue) and SCZ (orange). All neuronal cell types at supercluster level are displayed and sorted by average differential bias between ASD w/o ID and SCZ. Error bars represent standard error of the mean of cluster biases from corresponding supercluster. Asterisks indicate significance level between groups *: P < 0.05. **: P < 0.01. ***: P < 0.001, ****: P < 0.0001, Mann-Whitney U test with FDR correction. **D.** Bar plot comparing mutation biases across cell type superclusters between ASD w/o ID (blue) and ASD with ID (orange). **E-H.** Detailed comparison of mutation biases in specific neuron superclusters, each point representing a cell type cluster from the corresponding supercluster, P-values were calculated based on Mann–Whitney U test with FDR correction: **(E.)** Medium spiny neurons show stronger bias in ASD w/o ID, **(F.)** MGE interneurons exhibit stronger bias in SCZ, **(G.)** CGE interneurons display bias toward SCZ and **(H.)** CGE interneurons show stronger bias when comparing ASD with ID to ASD w/o ID.

### Distinct patterns of cell type mutation biases in ASD and SCZ

Given the overall correlation of cell type mutation biases between ASD without ID and SCZ, we next sought to identify cell types preferentially affected in one disorder relative to the other. To that end, we calculated, for each cell type, the differential mutation bias between ASD without ID and SCZ across cell type clusters within each cell type supercluster. This differential mutation bias quantifies how strongly each cell type supercluster is affected in one disorder relative to the other, thereby identifying cellular genetic insults that may contribute to the distinct behavioral and cognitive phenotypes associated with each disorder. We found that mutations in ASD without ID compared with SCZ or ASD with ID are differentially biased toward several specific neuron types (Figure 1C/D, Figure S10). Specifically, medium spiny neurons (MSNs) showed substantially stronger mutation biases in ASD without ID than in SCZ (Figure 1E). This agrees with previous findings implicating MSNs in ASD from computational^57^ and experimental studies^58,59^. In contrast, GABAergic interneurons, including MGE-derived (Figure 1F), CGE-derived (Figure 1G), and Lamp5-LHX6/Chandelier interneurons, showed strong differential biases toward SCZ compared with ASD without ID (Figure 1C), consistent with previous studies implicating interneurons in SCZ^25,26,60,61^.

To investigate which neuronal populations may be preferentially associated with cognitive dysfunction, we next compared cell type mutation biases between ASD without ID and ASD with ID (Figure 1D). We observed that multiple cell types, including MGE and CGE interneurons and MSNs, showed significantly stronger mutation biases in ASD with ID. This result is consistent with the generally stronger impact of mutations associated with ASD with ID across multiple cell types ^39,62^. Notably, the mutation bias toward CGE interneurons was especially strong (P = 4×10^-5^; Figure 1H) in ASD associated with cognitive impairment.

### CGE interneurons are strongly affected across multiple psychiatric phenotypes characterized by cognitive impairments

To further investigate the involvement of CGE interneurons in cognitive function, we analyzed several additional cohorts with related phenotypes (Figure 2A, see Methods). Consistent with the results observed in ASD with ID, we also observed a strong bias toward CGE interneurons (Figure 2A; Figure S10H) in the developmental disorders and intellectual disability (DD/ID) cohort^50^ (Figure S8D), which is characterized by severe cognitive impairments. Furthermore, we analyzed rare protein-coding variants from the UK Biobank associated with verbal-numerical reasoning (VNR)^64^, an assessment of fluid intelligence and the best proxy for general intelligence among the available traits^65^. We found that genes with mutations associated with lower VNR performance (VNR; see Methods), exhibited a pronounced bias toward CGE interneurons compared with genes positively associated with VNR (VNR+), which likely represent near-neutral controls (Figure 2A and Figure S10G). We also observed a strong bias toward CGE interneurons in SCZ. Cognitive impairments are widespread in SCZ, whereas cognitive symptoms in high-functioning ASD are often more localized and domain-specific^44–47^. Notably, the strong CGE mutation bias in SCZ was not driven by shared DD/ID genes; removing the 9 SCZ genes that overlap with the 285 DD exome-wide significant genes^50^ resulted in very similar CGE biases in SCZ (Figure S11A and B).

**Figure 2.**
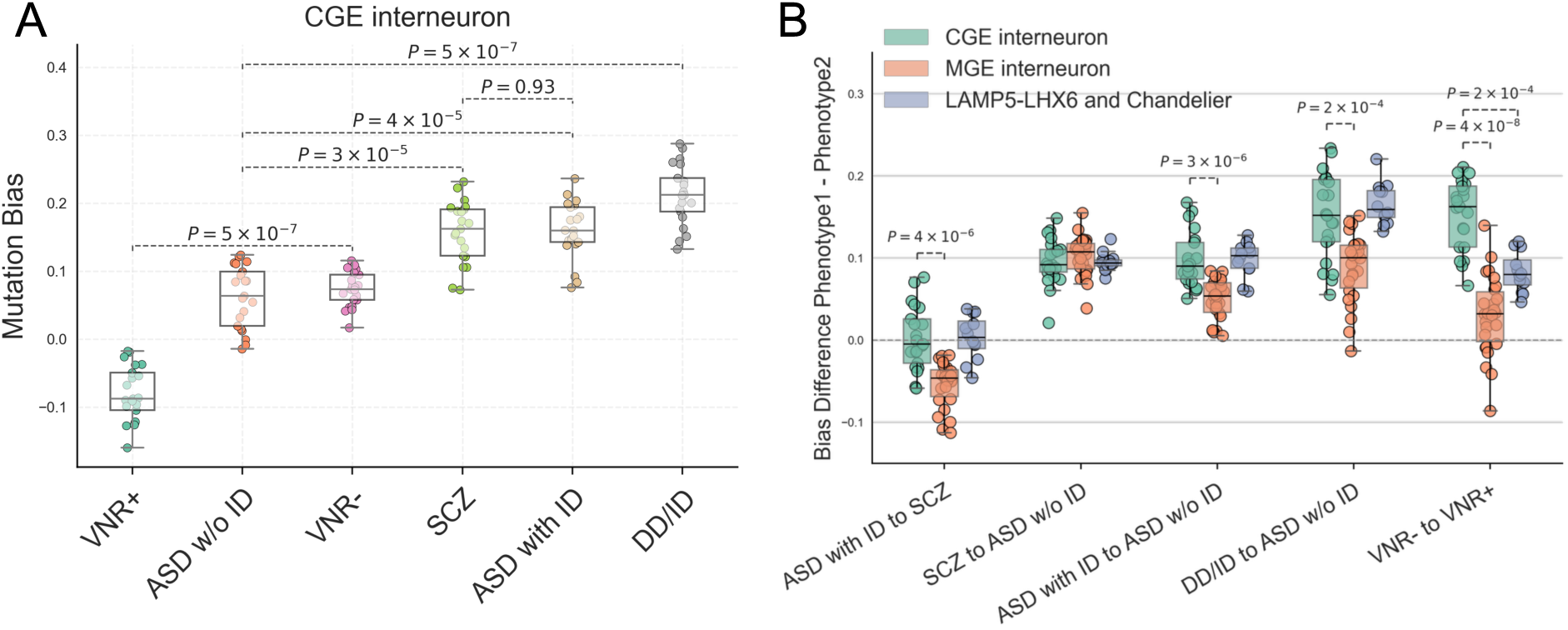
CGE interneurons are impacted in multiple cohorts with cognitive abnormalities. **A.** CGE interneuron mutation biases across different cohorts shows stronger bias in disorders with cognitive deficits. Each dot represents a CGE interneuron subtype. Comparison includes ASD w/o ID, verbal-numerical reasoning negative and positively associated genes (VNR- and VNR+), schizophrenia (SCZ), ASD with ID, and developmental disorders/intellectual disability (DD/ID). P-values indicate the significance of differences in CGE interneuron mutation biases between groups (e.g., SCZ, DD/ID and ASD with ID vs. ASD w/o ID, VNR– vs. VNR+) and were calculated using the Mann–Whitney U test with FDR correction. **B.** Pairwise mutation bias differences between disorders across 3 major interneuron superclusters. For each disorder pair (Phenotype1 – Phenotype2), the y-axis shows the difference in mutation bias per cell type cluster. Each dot represents one cluster from the three interneuron superclusters (CGE, MGE, LAMP5-LHX6/Chandelier). Colored boxplots indicate the supercluster of origin. Statistical comparisons were conducted between CGE clusters and each of the other two superclusters (MGE, LAMP5-LHX6/Chandelier) using the Mann–Whitney U test. Significant (0.05/10 comparisons) differences are annotated with the corresponding p-values.

We next compared mutation biases across the three major interneuron superclusters to assess differences between CGE interneurons and other interneuron populations in disorders associated with cognitive impairment (Figure 2B, Figure S11C and D). Although SCZ and ASD with ID showed correlated mutation biases across cell types, their interneuron profiles differed: MGE interneurons showed stronger biases in SCZ than in ASD with ID, whereas CGE interneuron biases were comparable between the two disorders. The increase in mutation bias in ASD with ID relative to ASD without ID was greater in CGE than in MGE interneurons (P=3×10^-6^). Similarly, between-disorder differences in mutation bias were greater in CGE than in MGE interneurons for DD/ID versus ASD without ID (P=2×10^-4^) and VNR-versus VNR+ (P=4×10^-8^). Together, the consistently strong CGE interneuron mutation bias across multiple cohorts supports the preferential involvement of these neurons in phenotypes characterized by prominent cognitive impairments.

### CGE/VIP+ interneurons are strongly affected by genes within the 22q11.2 deletion

Copy number variants (CNVs) consistently stand out as highly penetrant genetic risk factors for SCZ and ASD^15,36,66,67^. 22q11.2 deletion syndrome is a neurodevelopmental disorder characterized by a significantly elevated risk of SCZ and ASD, along with comorbidities such as intellectual disability^14,68^. Cognitive impairment is a key feature of 22q11.2 deletion syndrome, with nearly half of affected individuals displaying an IQ below 70^69,70^. Similar to most disease-associated CNVs, phenotypic symptoms associated with the syndrome result from the synergistic effects of multiple genes within the 22q11.2 deletion^14^. Given the significant cognitive disability phenotypes associated with the 22q11.2 deletion, we investigated the bias toward CGE and other interneurons among the deletion genes. We note that, in contrast to the large ASD, SCZ, and DD gene sets, only a small subset of genes within the 22q11.2 locus likely to contribute to disease risk^14,35^. Thus, our goal in this analysis was not the brain-wide identification of affected cell types (Figure S12), but rather to test a genetically motivated cross-dataset hypothesis that CGE interneurons, which are implicated in cognitive phenotypes by our rare-variant analyses across multiple disorders, are also strongly affected by genes within the 22q11.2 locus.

The 22q11.2 deletions occur in two main forms: a 3 Mb deletion (containing 46 genes) and a smaller 1.5 Mb deletion (containing 28 genes). Analysis of cell type biases for genes in the 3 Mb deletion (Figure S12) revealed a substantial bias toward CGE-derived interneurons compared with other major interneuron types (Figure 3A). Interestingly, among CGE interneurons, VIP+ cell types exhibited significantly stronger biases compared with VIP-CGE cell types (Figure 3B). This pattern remained consistent and significant when we examined genes within the 1.5 Mb deletion, which corresponds to the 16qA13 deletion in mice (Figure 3B). The selective impact on VIP+ interneurons compared with other CGE interneuron cell types suggests a particularly important role for these cells in mediating the severe cognitive and behavioral phenotypes associated with 22q11.2 deletion syndrome. CGE-derived VIP+ interneurons modulate neural circuits by reducing inhibition and enhancing excitatory activity^71,72^, an essential mechanism for cognitive functions such as perception, attention, learning, and memory^73^. They achieve this by suppressing Sst+, Cck+, and possibly Pvalb+ GABAergic interneurons^74,75^, thereby indirectly amplifying pyramidal neuron activity. Moreover, VIP+ interneurons interact with cholinergic, noradrenergic, serotonergic, and dopaminergic systems^75–80^ to maintain the excitation-inhibition balance critical for network stability and effective cognitive processing^73^.

**Figure 3.**
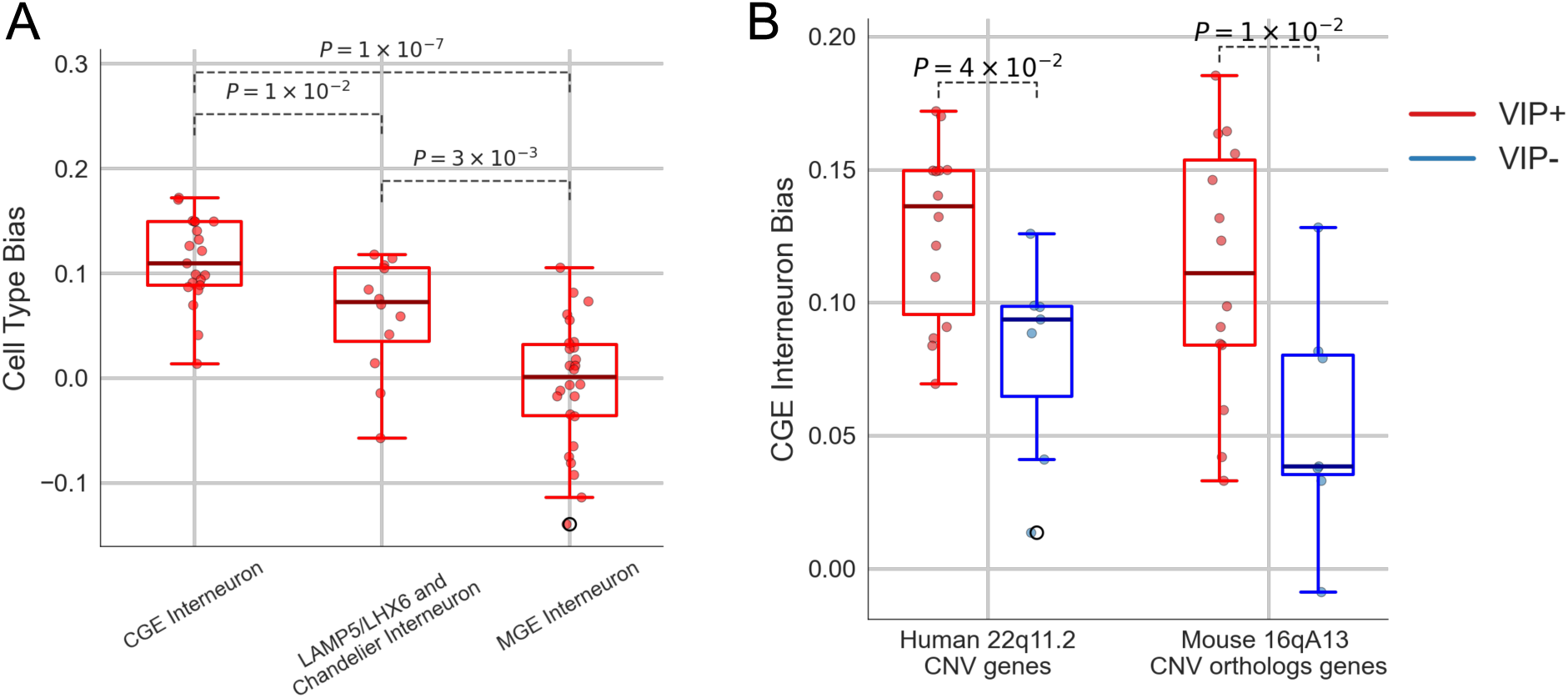
VIP+ CGE interneurons are preferentially affected by 22q11.2 deletion genes. **A.** Boxplot showing expression biases for three major interneuron types (CGE interneurons, Lamp5/LHX6 and Chandelier cells, and MGE interneurons) in the 22q11.2 deletion region. CGE interneurons exhibit the strongest bias compared to other interneuron populations (P-values shown for pairwise comparisons, Mann-Whitney U-test). **B.** The CGE interneuron cell type bias across different 22q11.2 gene sets, including complete deletion genes (3MB deletion, 46 genes) and 28 genes included in 1.5MB deletion which is equivalent to mouse 16qA13 deletion. VIP+ CGE types shown in red and VIP-CGE types shown in blue. Difference between VIP+ and VIP-bias is significant for all 22q11.2 genes and 1.5MB deletion genes (P=0.04 and 0.01, respectively, Mann-Whitney U-test.)

### VIP+ interneurons exhibit altered intrinsic properties in a mouse model of 22q11.2 deletion syndrome

Having identified a robust genetic association between mutation biases toward CGE interneurons and cognitive phenotypes across multiple genetic disorders, with a particular bias toward VIP+ interneurons compared with other interneurons in 22q11.2 deletion syndrome, we next experimentally investigated whether and how this bias is reflected in the altered function of these interneurons within hippocampal circuitry. Although excitatory neuron superclusters (i.e., deep-layer and upper-layer IT neurons) also showed elevated mutation bias for genes within the 22q11.2 locus (Figure S12), these populations were broadly and comparably affected across multiple phenotypes, including ASD without ID (Figure 1C, Figure S8). Therefore, they lacked the phenotypic specificity needed to distinguish biological mechanisms more selectively associated with cognitive dysfunction. In contrast, CGE interneuron bias was consistently associated with cognitive impairment phenotypes, making this interneuron population a strong candidate for experimental investigation. To this end, we used the *Df(16)A*^+/-^mouse model, which faithfully recapitulates the human 1.5 Mb 22q11.2 deletion^13,33,35,81^. This model, known for deficits in working and episodic memory, synaptic plasticity, and cortical and hippocampal development, has proven invaluable for uncovering novel biological pathways relevant to cognitive dysfunction and psychiatric disorders^14,35,36,68,81–84^. Using the model, we explored how mutational impact on CGE/VIP+ interneuron physiology and circuit function may contribute to altered cognitive processes.

To investigate the cell intrinsic properties of VIP+ interneurons in the *Df(16)A*^+/-^mouse model, we performed patch-clamp slice electrophysiology experiments. *Df(16)A*^+/-^mice were crossed with a *VIP-eGFP* reporter line to facilitate the identification of VIP+ cells in fresh tissue (Figure 4A, Methods). Slice preparation was performed as previously described^85^. In brief, 300 µm coronal sections were taken to capture dorsal CA1 in 3-4 slices, and eGFP+ interneurons were then patched (Figure 4B). We measured action potential frequency and resulting membrane voltages with increasing holding current injections (Figure 4C) and found that *Df(16)A*^+/-^VIP+ interneurons were less likely to fire action potentials compared with wildtype control cells (Figure 4C/D). At low current injections, mutant cells exhibited more hyperpolarized membrane potentials (Figure 4E). We observed an increase in action potential half widths and a significant decrease in rise speed in *Df(16)A*^+/-^VIP+ interneurons compared with controls (Figure 4F,4G). Altogether, our slice electrophysiology data confirmed significantly altered intrinsic properties of hippocampal VIP+ interneurons in *Df(16)A*^+/-^mice, culminating in reduced firing.

**Figure 4.**
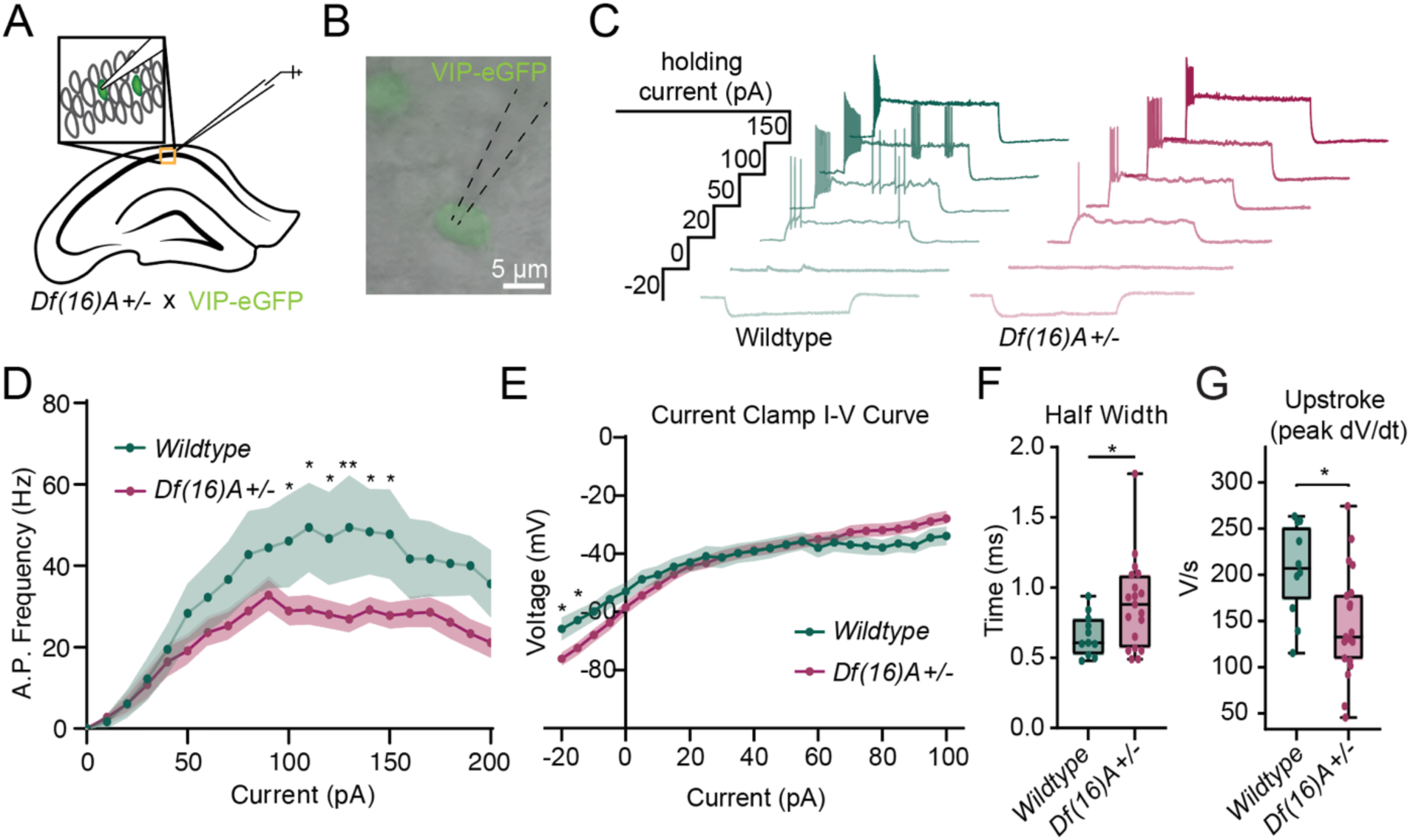
*Df(16)A+/-* VIP+ interneurons exhibit reduced firing in slice experiments. **A.** Schematic of tissue preparation and targeting strategy for patch-clamp experiments. **B.** Example of VIP-eGFP+ patched interneuron in *stratum pyramidale.* **C.** Representative traces from select current steps in the current injection protocol from each genotype. **D.** Action potential firing frequency analysis at each step of the current step injection experiment (2-way ANOVA: P = 0.0488). **E.** Current clamp I-V curve between genotypes (2-way ANOVA: P = 0.8761). **F.** Half width comparison between genotypes. (WT: 0.6618 ms, n = 3 mice, 11 cells ; *Df(16)A^+/-^*: 0.8832 ms, n = 4 mice, 19 cells, Mann-Whitney U: P = 0.0477). **G**. Upstroke peak speed of action potential rise (Wildtype: n = 11 cells; Df(16)A+/-: n = 19 cells, Mann Whitney-U: p = 0.0231). *: P < 0.05.

### Diminished VIP+ *in vivo* interneuron activity and population specificity in a mouse model of 22q11.2 deletion syndrome

To better understand how cell-intrinsic deficits in VIP+ interneurons affect their circuit function, we next investigated their activity in behaving mice. To examine VIP+ interneuron activity *in vivo*, we crossed *Df(16)A^+/-^*mice with VIP-Cre mice and locally injected an adenovirus carrying a floxed GCaMP8m or GCaMP7f calcium indicator into the CA1 region to selectively express GCaMP in VIP+ interneurons (Figure S13A). Using two-photon three-dimensional acousto-optic deflector (3D-AOD)^86–88^ microscopy, we then recorded VIP+ interneuron activity within a three-dimensional volume of the dorsal hippocampal CA1 region (Figure S13B-S13D). Tissue registration and *post-hoc* immunohistochemistry^32,86,89^ (see Methods, Figures S13E and S13F) allowed us to identify four major VIP+ interneuron subtypes: interneuron-selective interneurons type 2 (ISI2), interneuron-selective interneurons type 3 (ISI3), VIP+ cholecystokinin basket cells (CCKBCs), and a group of VIP-expressing cells, referred to as "VIP+ only" (VIPo) cells, that were immunonegative for all other tested markers^71,90,91^ and reside in the *stratum pyramidale* or *stratum oriens*. The distribution of these subtypes across hippocampal layers did not significantly differ between wildtype and *Df(16)A^+/-^*mice (Figures S13G and S13H), and comparable numbers of each interneuron subtype were recorded from both wildtype and the mutant genotype (Figure S13I). Thus, our analysis found no evidence that the 22q11.2 deletion affects the number or relative abundance of VIP+ interneuron subtypes in the hippocampal CA1 region.

As previously established, global interneuron activity is strongly driven by ambulatory state^92–95^, and in treadmill-based experiments we have consistently observed locomotion to be the strongest predictor of global interneuron activity^32,86,96^. To investigate locomotion-related activity of hippocampal VIP+ interneurons, mice were next trained to run on a head-fixed, cue-rich 2-meter treadmill for randomly distributed water rewards while undergoing calcium imaging (Figure 5A)^32,86,89^. Behavioral metrics, including laps run, licking, and belt-region occupancy during the 10-minute trials, were similar across genotypes (Figures S13J-S13M). Expected locomotion-related activity patterns were preserved across genotypes, with the activity of ISI3, ISI2, and VIPo subtypes positively correlated with movement, whereas the activity of VIP+ CCKBCs was anticorrelated (Figure 5B)^86,97^. However, in *Df(16)A^+/-^*mice, the activity of ISI3 and ISI2 cell activity showed significantly weaker correlations with locomotion (Figure 5C). To further investigate locomotion related activity, we examined Run-Start and Run-Stop related activity assessed by Peri-event Time Averages (PETAs, see Methods) and found altered responses in the ISI2 and VIPo subtypes (Figure S14A-D). In addition, ISI3, ISI2, and VIPo interneurons all showed visibly lower calcium transient amplitudes (Figure 5B). Due to the spike-like activity of ISI3, ISI2, and VIPo cells we were able to perform a transient waveform analysis (Figure S14E, see Methods^98^). This analysis confirmed reduced amplitudes in ISI3, ISI2, and VIPo cells (Figures 5D-J), while VIPo cell transients uniquely exhibited a higher transient frequency (Figure 5J, Figures S14F and S14G). Reward responses, again assessed using PETAs, exhibited genotype-dependent differences, including reduced VIPo response amplitudes and enhanced ISI2 and ISI3 response amplitudes (Figures 5K, 5L), suggesting that ISIs in mutant mice exhibit increased sensitivity to randomly occurring stimuli^96^. Overall, these results showed that VIP+ interneurons in the hippocampal CA1 region exhibit reduced *in vivo* activity dynamics in the 22q11.2 deletion model and that ambulatory and reward responses are significantly altered among specific VIP+ subtypes, particularly interneuron-selective mouse subtypes. Following these experimental results in mice, we attempted to perform cross-species transcriptomic mapping of mouse VIP+ interneuron subtypes onto human CGE transcriptional clusters^37^. We found that broadly defined VIP+ clusters, rather than more granular individual subtypes, represent the most robust unit for detecting mutation-bias signals (see Figure S15 and Supplementary Text).

**Figure 5:**
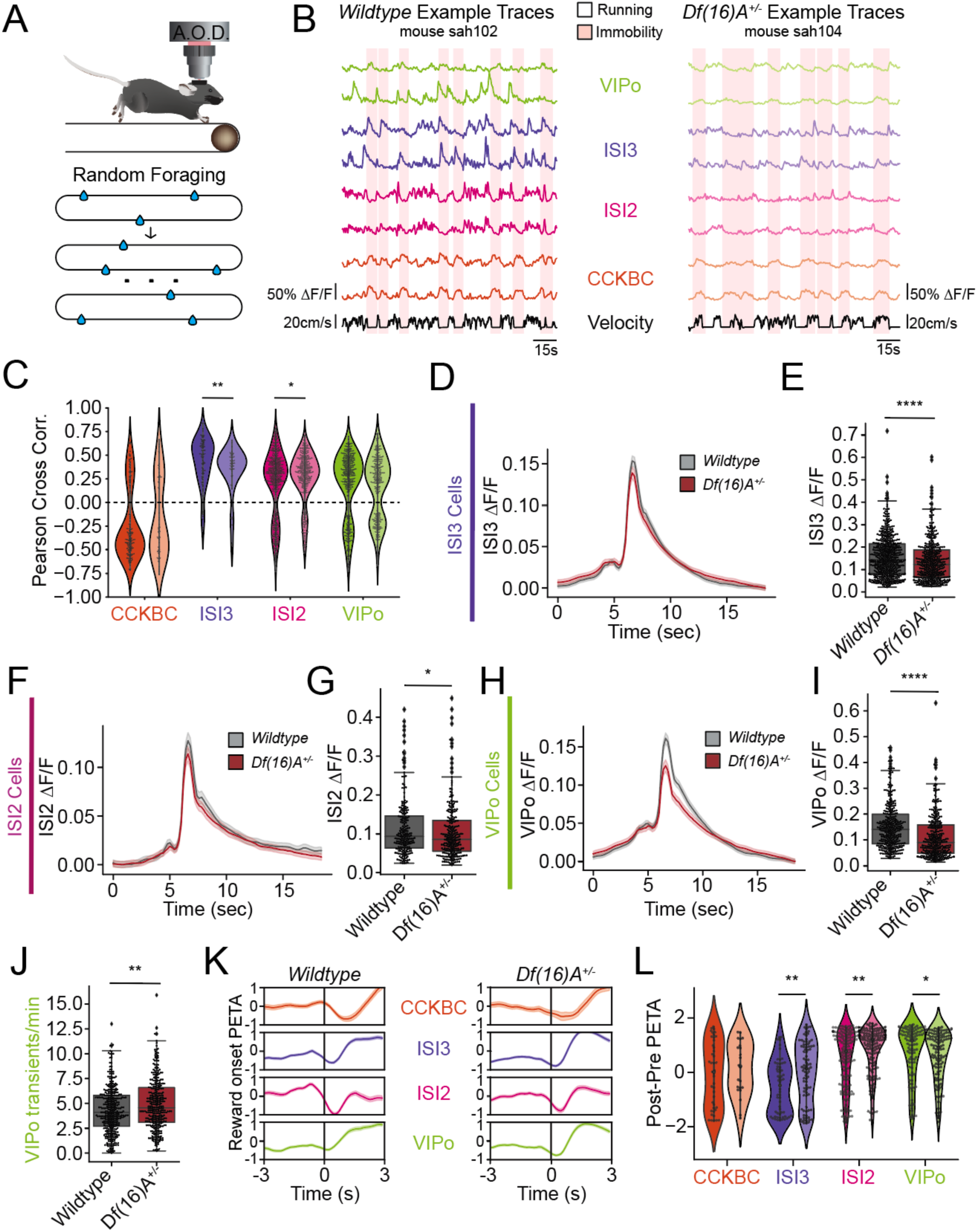
Baseline changes in VIP+ cell type calcium dynamics in mutant mice. **A.** Imaging and behavior schematic: mice are trained to run on a cue-rich belt to expect random water rewards to be deposited across the belt (bottom) while head-fixed and imaged with 2-photon 3D-AOD microscopy (top). **B.** Example traces from all four VIP+ cell types detected in a single session from a Wildtype littermate (left) and a *Df(16)A^+/-^*(right) mouse. **C.** Pearson correlation coefficients of mouse velocity against average subtype calcium activity. **D.** Waveform comparison of average detected calcium transient from wildtype littermates (black) and *Df(16)A^+/-^*(brown) ISI3 cells. **E.** Average amplitude of detected ISI3 waveforms compared between genotypes. **F.** Waveform comparison of average detected calcium transients from wildtype littermates (black) and Df(16)A^+/-^(brown) ISI2 cells. **G.** Average amplitude of detected ISI2 waveforms compared between genotypes. **H.** Waveform comparison of average detected calcium transients from wildtype littermates (black) and *Df(16)A^+/-^*(brown) VIPo cells. **I.** Average amplitude of detected VIPo waveforms compared between genotypes. **J.** Number of transients from VIPo cells per cell between genotypes. **K.** Averaged z-scored reward-onset peri-event time averages (PETAs) between -1.0 and 1.0 z-scores by cell type across all cells within cell type during the Random Foraging task for 3 seconds pre- and 3 seconds post-event. **L.** Post - Pre PETA responses per cell type before and after reward onset comparing the 3 seconds pre- to 3 seconds post-event. *CCKBC:* VIP+ and cholecystokinin + basket cell; *ISI3:* Interneuron-selective interneuron type 3; *ISI2:* Interneuron-selective interneuron type 2; *VIPo*: VIP+/Cal-/M2R-/CCK-cells in layer oriens or layer pyramidale. *: P < 0.05. **: P < 0.01. ****: P < 0.0001. All experiments and analyses performed with n = 4 mice per genotype. See Table S1 for detailed statistics.

Having identified cell-intrinsic deficits in *Df(16)A^+/-^* VIP+ interneurons both in slices and *in vivo,* we next sought to determine how these altered properties may impact the hippocampal circuitry. Hippocampal place cells, a subset of pyramidal neurons, are active in specific environmental locations and provide cellular substrates for episodic memory and goal-directed spatial learning^99,100^. In *Df(16)A^+/−^* mice, our previous study revealed significant impairments in learning performance when both contextual cues and reward locations were altered during a spatial navigation task^83^. These learning deficits were associated with reduced long-term stability of place fields, impaired context-related remapping and plasticity, and a lack of reward-related reorganization. Further investigation showed that parvalbumin-and somatostatin-expressing GABAergic interneurons, which are directly inhibited by VIP+ interneuron activity, displayed diminished place fields and heightened reward-related activity, likely contributing to the observed place cell deficits^32^. Changes in the activity dynamics of VIP+ cells in dorsal CA1 may also reflect altered spatial information encoding by these cells. To explore this, we next evaluated how well mouse position could be decoded based on VIP+ subtype activity during random foraging (Figure 5A) by training a support vector machine using the activity of all recorded VIP+ interneurons (Figure 6A). Decoding error did not differ between genotypes when the total number of imaged cells per session was not normalized. However, when controlling for the number of cells included in the analysis (Figure 6B) or the number of laps run per session (Figure 6C), decoding error was significantly higher in *Df(16)A^+/-^*mice, indicating that individual mutant cells convey less spatial information than cells in wildtype mice. Furthermore, analysis of place preference in VIP+ activity revealed that fewer VIP+ cells in *Df(16)A^+/-^*mice exhibited significant place fields, with a reduced proportion of cells exhibiting at least one place field (Figure 6D). These findings demonstrate that VIP+ interneurons in the 22q11.2 deletion model exhibit markedly impaired spatial encoding within the CA1 microcircuit.

**Figure 6.**
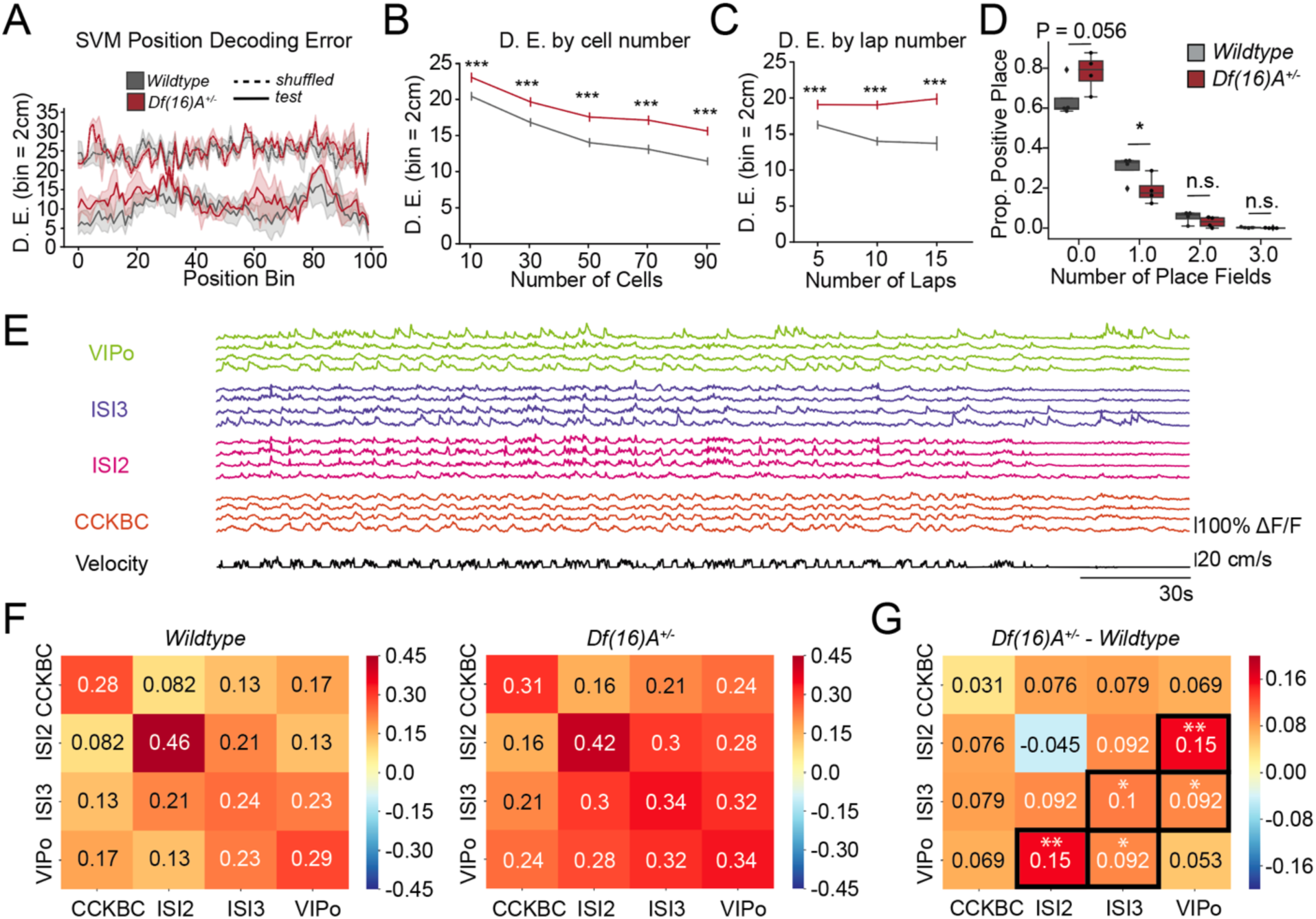
Reduced specificity in *Df(16)A^+/-^* VIP+ interneuron population activity. **A.** All recorded VIP+ interneuron activity was used to train a support vector machine (*SVM*) to decode discrete position bins (*D.E.*: decoding error) across the belt. Each bin is equivalent to 2 cm. **B.** SVM results bootstrapped 100 times by number of cells used per genotype per bin. **C.** SVM results bootstrapped 100 times by the number of laps used per genotype per bin. **D.** Proportion of positive place fields detected using the Peak Place method. **E.** Example traces of different VIP+ subtypes exemplifying differences in within group and between group correlated activity. **F.** Pairwise Pearson’s correlation coefficient value matrix between cell types of (Δ*F*/*F*) traces across the trial during the Random Foraging paradigm from WT (left) and *Df(16)A^+/-^*(right). **G.** Comparative differences between cell type correlated activity between genotypes. *CCKBC:* VIP+ and cholecystokinin + basket cell; *ISI3:* Interneuron-selective interneuron type 3; *ISI2:* Interneuron-selective interneuron type 2; *VIPo*: VIP+/Cal-/M2R-/CCK-cells in layer oriens or layer pyramidale. *: P < 0.05. **: P < 0.01. ***: P < 0.001. All experiments and analyses performed with n = 4 mice per genotype. See Table S1 for detailed statistics.

Local brain circuits display both coordinated and unique activity patterns depending on input properties, connectivity, and brain state. We observed that VIP+ interneuron subtypes exhibited distinct patterns of correlated activity both within and across subtypes (Figure 6E). Taking advantage of our simultaneous recordings of multiple VIP+ subtypes, we computed pairwise Pearson’s correlation coefficients between calcium activity traces within and between different VIP+ subtypes (see Methods). We found that VIP+ cells in *Df(16)A^+/-^*mice exhibited unusually high correlations between subtypes during random foraging. This suggests a loss of subtype-specific heterogeneity of *in vivo* activity dynamics, particularly among disinhibitory VIP+ subtypes, indicating a potential microcircuit-level disruption and impaired fine-scale coordination between VIP+ subtypes (Figures 6F and 6G). These findings reveal that VIP+ interneurons in the CA1 region of *Df(16)A^+/-^*mice exhibit impaired spatial encoding, reflected in increased decoding errors and reduced place preference, and that this disruption is also accompanied by a loss of subtype-specific heterogeneity, evident from unusually high inter-subtype correlations during the foraging task. These physiological consequences illuminate how perturbations of VIP+ interneurons disrupt network dynamics and information processing relevant to cognition.

## Discussion

Cognitive dysfunction is central to psychopathology, as many psychiatric conditions share overlapping yet challenging-to-treat cognitive impairments^19–22,40,69^. For instance, SCZ often presents with deficits in executive function and working memory^8,12^. Similarly, ASD cases driven by rare damaging mutations^57,101^ are frequently associated with pronounced cognitive deficits. Cognitive disabilities are also common in genetic conditions such as 22q11.2 deletion syndrome^14,68,69^. In this study, we focused on the cell type-specific impact of rare, highly penetrant genetic mutations across disorders^1,2,50^. We observed a correlation between cell type biases in ASD and SCZ, which was primarily driven by mutations in highly constrained genes. We also identified disorder-specific cell type mutation biases. For example, medium spiny neurons of the striatum showed significantly stronger mutation biases in ASD compared to SCZ, consistent with the core ASD symptoms of repetitive behaviors and social impairments, which have frequently been associated with striatal dysfunction^58,59^. In contrast, multiple interneuron subtypes exhibited stronger mutation biases in SCZ, consistent with recent GWAS-based studies^25,26^, particularly parvalbumin-expressing interneurons crucial for gamma oscillations and cognitive processing^102,103^, as well as somatostatin-expressing interneurons important for information processing^60^. Supporting these findings, transcriptomic studies have observed alterations in these interneuron populations in SCZ patients^61^.

Our genetic analysis revealed CGE-derived GABAergic interneurons as an important locus of vulnerability contributing to cognitive deficits across multiple psychiatric and neurodevelopmental conditions (Figure 2). Specifically, we found that CGE interneurons are more affected in SCZ compared to high-functioning ASD, consistent with cognitive impairments being a core and defining symptom of SCZ^21,22^. We also observed a strong genetic impact on CGE interneurons in low-functioning compared to high-functioning ASD, as well as in intellectual disability and 22q11.2 deletion syndrome. Extending this pattern, CGE interneurons also showed strong biases in UK Biobank gene sets associated with lower cognitive performance^64^.

Within the broader class of CGE interneurons, VIP+ subtypes were strongly affected by genes within the 22q11.2 deletion region, a highly penetrant CNV conferring risk for SCZ, ASD, and intellectual disability. To investigate the functional consequences, we evaluated the intrinsic properties and functional activity of VIP+ CGE-derived interneurons using the *Df(16)A^+/−^*mouse model. We observed significant defects at both the cellular and circuit levels, which likely contribute to broader hippocampal dysfunction previously reported in this mouse model^32,83^. Our cell-intrinsic electrophysiology results identified reduced firing and slower action potential recovery. *In vivo*, VIP+ interneurons exhibited dampened calcium activity, suggesting a reduction in the top-down disinhibition^73,104^ normally exerted by these cells in cortical circuits. In addition, we found a reduction in the spatial information conveyed by their activity. Because synaptic plasticity is critical for fine-tuning synaptic connections and forming dynamic spatial representations and episodic memories, this reduction may reflect impaired plasticity or other alterations in circuit dynamics. Such alterations may hinder the ability of MGE-derived interneurons and pyramidal neurons to effectively encode and update spatial information within the hippocampal network^32,83^. Furthermore, we observed increased correlations in the *in vivo* activity of VIP+ interneuron subtypes, suggesting less distinct activity patterns and potentially reduced signal diversity in the circuit. Given that VIP+ disinhibitory motifs are highly conserved across cortical circuits^71,72,105^, the deficits we uncovered in the CA1 circuit may also occur in neocortical brain regions. However, these experiments do not establish that VIP+ dysfunction alone causes the broader hippocampal phenotype, which may also reflect abnormalities in other neuronal populations or altered excitatory and neuromodulatory inputs. Cell-type-specific perturbation and rescue experiments, together with recordings from additional interneuron populations, will be needed to distinguish these contributions.

Because VIP+ interneurons inhibit other major interneuron subtypes, their activation increases pyramidal neuron excitability, thereby improving the detection of low-contrast signals^91,106^. Thus, these interneurons are essential for learning and attention, as they respond to salient stimuli by adjusting overall neural excitability. For instance, during fear conditioning, blocking VIP+ interneurons disrupts projection neuron activity and impairs fear learning^72^. In the hippocampal CA1 region, proper disinhibitory VIP+ interneuron function is vital for spatial learning as excessive inhibition of pyramidal neurons hinders place cell remapping and performance, while their activation enhances learning outcomes^96^. Similarly, in the prefrontal cortex, activation of VIP+ interneurons improves working memory and attention, whereas activation of Pvalb+, Sst+, or pyramidal neurons may lead to detrimental effects^80,96,106,107^. Notably, impaired disinhibition and CGE-derived VIP+ interneuron dysfunction have also been previously described in other neurodevelopmental disorders^108–110^. In Rett syndrome selective deletion of Mecp2 in CGE/VIP+ interneurons, but not in Pvalb+ or Sst+ interneurons, leads to deficits in cortical state transitions and social memory^109^. In Dravet syndrome, SCN1A mutations lead to persistent VIP+ interneuron dysfunction in juvenile and adult mutant mice, contributing to social and cognitive deficits^108,110^.

While further research is needed to link specific disease-related behaviors to CGE/VIP+ interneuron circuit dysfunction, our combined computational and experimental results suggest that disruption of VIP+ interneuron activity may be a key contributor to cognitive deficits in 22q11.2 deletion syndrome and potentially in other neurodevelopmental and psychiatric disorders associated with cognitive impairment^111^. Given the persistent cognitive comorbidities across multiple psychiatric disorders, pharmacologically targeting CGE/VIP+ interneuron dysfunction is a potential avenue for therapeutic intervention.

## METHODS

### Rare mutation genetic data for neuropsychiatric disorders and phenotypes

For the analysis on cell type mutation biases of ASD and SCZ, we primarily used the aggregated mutation data obtained from the SFARI Simons Foundation Powering Autism Research for Knowledge (SPARK) cohort and the Schizophrenia Exome Sequencing Meta-Analysis (SCHEMA) cohort. In the SPARK study^2^, de novo mutations were identified through familial sequencing of 42,607 ASD probands and 8,267 unaffected siblings. We focused our analysis on de novo likely gene-disrupting (LGD) mutations and damaging missense (Dmis) mutations. Dmis mutations were identified based on the REVEL score^112^, which estimates the probability of a missense mutation is pathogenic; mutations with REVEL scores higher than 0.5 were defined as deleterious. For our main analysis, we used 61 genes that reached exome-wide significance (P-value < 1.3x10^-6^) identified in the SPARK study, using the DeNovoWEST statistical model^50^. In the SCHEMA study^1^, which analyzed 24,248 SCZ cases and 97,322 controls, to match the number of analyzed ASD genes, we selected the top 61 genes with the strongest association for our analysis (P-value < 1.0x10^-3^ or FDR < 0.3). Dmis mutations in SCZ were identified based on the MPC score^113^, and we included in our analysis LGD and Dmis mutations with MPC scores greater than 3. We also investigated cell type biases across different cohorts with cognitive abnormalities. Specifically, we investigated genes within the 22q11.2 deletion region as defined by Karayiorgou *et al.*^14^. We also analyzed genes associated with verbal-numerical reasoning (VNR) from the UK Biobank study by Chen *et al.*^64^. We separately analyzed genes with positive and negative impacts (indicated by the beta coefficients of gene-phenotype associations), focusing on European population data for optimal statistical power. Additionally, we analyzed genes associated with developmental disorders and intellectual disability (DD/ID) from Kaplanis *et al.*^50^, ranking the top genes according to DeNovoWEST P-values.

### ASD subgroup definition

ASD probands were stratified by cognitive function into two subgroups based on full-scale IQ (FSIQ) scores available in the SPARK, SSC and ASC cohort^2,62^. FSIQ phenotyping is available for a subset of the full ASD cohort, and we restricted the subgroup analysis to the 5,402 probands with both exome sequencing and FSIQ data. "ASD with ID" was defined as probands with FSIQ ≤ 70 (N = 1,792), and "ASD without ID" as probands with FSIQ > 70 (N = 3,610). This categorization is defined by our analysis according to reported FSIQ (when available) and does not correspond to inherent clinical labels in the source datasets. The underlying de novo mutation data were drawn from the same cohort and sequencing pipeline^2^, ensuring that the only systematic difference between subgroups is cognitive ability. De novo mutation counts and gene weights were recomputed separately for each subgroup using the same statistical framework applied to the full ASD cohort.

### Human whole brain single-cell data

For our single-cell analysis, we used the recent human brain single-nucleus sequencing dataset^37^, comprising 3,369,219 nuclei sampled from 105 regions across the entire brain of three donors. This comprehensive single-cell brain atlas was classified, based on cellular transcriptional similarities and developmental origins, into 461 cell type clusters organized into 31 cell type superclusters.

### Gene expression specificity score for individual cell types

To identify neuronal populations preferentially impacted by rare mutations, we calculated a gene expression specificity score for each cell type cluster (Figure S3). To that end, for each gene *g* in each cell type *ct*, we first quantified gene expression using Transcripts Per Million (TPM), calculated from raw UMI counts as:

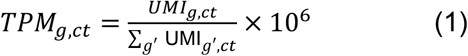

where *UMI*_g,*ct*_ is the expression count of gene *g* in a cell type *ct*, and the sum in the denominator is across all genes *g*′ in that cell type. In this way, TPM values normalized expression across cell types by accounting for the differences in the total transcript abundance. To filter out genes with very low expression in individual cell types and reduce noise in the specificity calculation, we applied a minimum TPM threshold of 0.1 for each gene in each cell type. Additionally, genes with total UMI counts below 10,000 across all 3 million cells of the human brain cell atlas were excluded from the analysis, removing genes with very low overall expression.

Next, we computed a cell type specificity score to quantify how selectively a gene is expressed in each cell type. For each gene *g*, specificity toward cell type *ct* was calculated as:

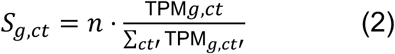

where *n* is the total number of cell types. In case of the uniform expression of a gene across all cell types, the expected value of *S*_g,*ct*_ is 1, and higher values indicate stronger cell type specific expression. To prevent extreme values from a small number of genes from dominating downstream bias computations, we capped *S*_g,*ct*_ at twice the global mean specificity across all genes and cell types (approximately 2.0, as the expected value under uniform expression is 1). This cap prevented technical inflation of specificity in cell types with low sequencing depth, as high uncapped specificity was disproportionally observed in cell types with small total UMI counts (Spearman ρ = −0.50 between the fraction of genes exceeding the cap and total UMI per cell type; Figure S4A). The 2× cap was chosen empirically: Spearman’s correlations between mutation-bias rankings computed with the 2× cap and using alternative caps in the 1.5–5× range exceeded 0.88 for all three disorder cohorts (Figure S4B).

To further ensure comparability across cell types with different expression distributions, we applied a per-cell type centering step. For each cell type, *ct*, we subtracted the average specificity across all human genes:

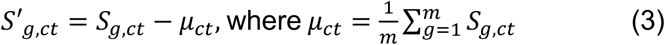

with *m* denoting the total number of genes. The resulting centered specificity score *S*′_g,*ct*_ reflects how strongly a particular gene is enriched in a given cell type relative to all genes in that cell type.

### Disorder mutation bias for individual cell types

To calculate cell type-specific mutation biases for each disorder cohort, we weighted gene expression specificities (*S*′_g,*ct*_, Equation 3) with their corresponding genetic burdens. This weighted approach accounted for the relative genetic contributions, proportionally to the genetic burden, of individual genes to the disorder. Specifically, for the ASD cohort, we quantified each gene’s contribution with the following mutation weight:

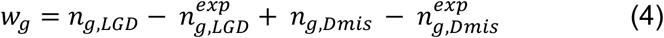

where *n*_g,*LGD*_ and *n*_g,*Dmis*_ are the numbers of observed *de novo* LGD and Dmis mutations in gene 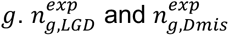 representing number of expected LGD and Dmis mutations for each gene, given the cohort size.

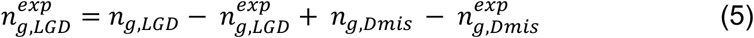

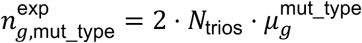

Where 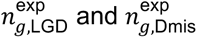 are the expected numbers of de novo LGD and damaging missense mutations in gene *g*, computed as the product of the number of trios (*N*_trios_), ploidy (2), and the sum of trinucleotide-context-specific mutation rates 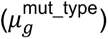 across all coding positions in the gene^114^. This normalization accounts for gene length and sequence-context-dependent mutability, ensuring that the mutation weight reflects excess burden above the baseline expectation.

For the SCZ cohort, we calculated mutation biases by integrating both *de novo* and inherited mutations identified in case-control studies. Since the genetic evidence comes from these two distinct sources, we computed a mutation counts that combine their contributions. For case-control mutations, the effective count for each gene represented the number of excess mutations in cases after subtracting the expected counts, which were calculated by scaling control mutations by the ratio of the number of cases to controls. We applied this approach separately for each mutation class (LGD and Dmis) and then combined these values with observed *de novo* mutations:

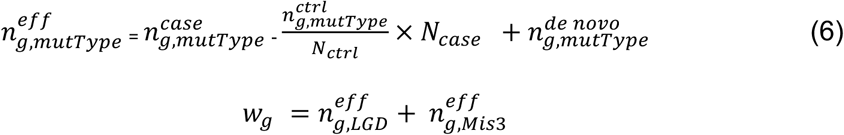

where 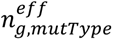 is the effective count of mutation in gene *g*, 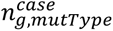 is number of mutations observed in case cohort in gene, 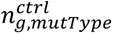 is number of mutations observed in controls in gene *g*, 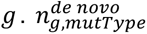 is number of de novo mutations in gene *g*. *mutType* includes LGD and Mis3 mutations. *N_case_* and *N_ctrl_* are total number of cases and controls in SCZ cohort, respectively.

For genes associated with the UK Biobank cognitive phenotypes, we utilized each gene’s effect size, i.e., beta coefficient from gene-phenotype association tests, as the gene weight. This approach assumes that genes with larger phenotypic effects contribute more strongly to cell type biases relevant to cognition. For genes in the 22q11.2 deletion, we applied uniform weights for all genes (*w*_g_ = 1), as each individual gene effectively contributes one LGD mutation to the CNV.

Finally, based on the calculated mutation weights for each gene (Equations 4-6), the mutation bias for all disorder-associated genes toward a cell type *ct* was calculated as follows:

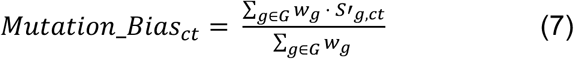

where the summation is over genes *g* in the set *G* containing all implicated genes for a disorder or a phenotype, and *w*_g_ is the mutation weight of each gene *g*.

#### Non-brain negative control traits

To confirm that neuronal cell-type enrichment is specific to psychiatric and cognitive phenotypes rather than an artifact of the analytical framework, we applied the full mutation-bias pipeline to 3 non-brain traits from UK Biobank exome-wide rare-variant burden analyses^115^: HDL cholesterol, inflammatory bowel disease (IBD), and alanine aminotransferase. For each trait, we selected the top 61 genes ranked by loss-of-function burden P-value and assigned weights according to beta coefficient of LoF burden. None of the non-brain traits showed significant enrichment in neuronal superclusters (Figure S7), confirming that the framework identifies trait-relevant cell types only where they exist.

### Statistical significance of the cell type mutation biases

To determine the statistical significance of the calculated cell type mutation biases, we used a permutation-based framework that compared the observed bias in disorder data against null distributions derived from random gene sets. For each considered disorder or phenotype, we generated 10,000 random gene sets matched for the gene set size, drawn from genes with available expression specificity scores in the atlas. For each random gene set, we computed the cell type mutation bias profile using the same weighted-average procedure (Equation 7). The empirical P-value for each cell type was then calculated as the fraction of null gene sets producing an equal or greater bias; P-values were corrected for multiple testing across cell types using the Benjamini-Hochberg false discovery rate (FDR) procedure.

To evaluate whether cell-type mutation bias results could be driven by gene-level confounders, we also implemented an alternative permutation framework in which null gene sets were matched to the disorder gene set for three properties: gene length (number of CDS bases), whole-brain expression level (mean expression across BrainSpan developmental stages and samples), and evolutionary conservation (average phastCons score). For each permutation, we drew 1,000 random candidate gene sets from genes across the genome, each containing the same number of genes as the disorder gene set, and retained the candidate set with the minimal Euclidean distance between the mean percentile values of these three variables and the corresponding values for the disorder gene set:

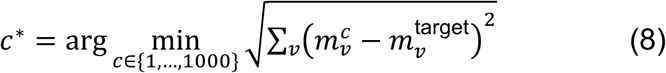

where *v* indexes the three matching confounder variables, 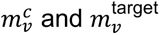 are the corresponding mean percentile values for the variable *v* across the c-th candidate gene set and the disorder gene sets, respectively. This procedure was repeated 10,000 times per trait to construct the matched null distribution. As expected, confounder matching reduced the nominal significance of several traits (Figure S7) because many genes with matched conservation and brain expression are themselves enriched for CNS-related functions^63^. FDR-corrected significance values from both the random and confounder-matched null models are reported for all traits in Table S4 and S5.

### Robustness to gene set size

To assess the sensitivity of mutation-bias estimates to the number of disorder genes used to calculate cell type mutation biases, we recalculated cell type mutation bias profiles using gene sets ranging from the top 10 most significant genes to the top 200 most significant genes for each disorder, ranked by the statistical significance of their disorder association. This analysis was performed separately for each of the four disorder gene sets: SCZ, ASD without ID, ASD with ID, and DD (Figure S5A–D). For each disorder, genes were ordered by their association significance in the corresponding study, and progressively larger sets were constructed by adding genes in decreasing order of significance. At each set size, the full cell type mutation bias profile was recomputed using the same burden-weighted, centered-specificity procedure used in the main analysis (Equation 7), so that only gene-set membership varied between runs. To measure how the bias profile changed as additional genes were included, we computed the Spearman‘s correlation across cell types between each expanded profile and the original 61-gene profile used in the main analysis (Zhou et al., 2022). We also calculated the association significance (−log₁₀ P) of the least significant gene included at each set size (Figure S5), providing a reference for the strength of the association signal among the additional genes.

### Impact of genetic architecture on inference stability

To empirically characterize how genetic architecture affects cell type mutation bias estimates, we performed a cohort-downsampling analysis. For each disorder, mutation counts were binomially subsampled at fractions of 10–90% of the full cohort (100 bootstrap iterations at each fraction). Gene discovery was then repeated on each subsample using Poisson rate tests for de novo cohorts (ASD, DDD) or one-sided Fisher’s exact tests for case–control data (SCZ). Each variant class (loss-of-function and damaging missense) was tested separately and the resulting P-value were combined using Fisher’s method. For SCZ, case mutation counts were subsampled while the control cohort was held fixed at N = 97,322, reflecting the realistic scenario in which future studies would expand case ascertainment against existing large control resources (e.g., gnomAD). Gene weights were recomputed for each iteration using BGMR correction for de novo cohorts and case-control excess correction for SCZ, consistent with the weighting schemes in Equations 4-6. Cell type mutation bias profiles were then recalculated from the resulting top 61 genes, matching the gene set size used in the original analyses.

### Sensitivity of the correlation between ASD w/o and SCZ cell type mutation biases to gene properties

To assess how gene-level properties (mutation constraint and brain expression level) influence the correlation of cell-type mutation biases between ASD without ID and SCZ, we evaluated how this correlation changes as genes are sequentially removed from each gene set according to either property. Gene constraint was quantified using LOEUF scores from the gnomAD v4.0 release ^63^; if a gene lacked a LOEUF score in v4.0, we used the corresponding value from gnomAD v2.1. Brain expression level was obtained from the BrainSpan^116^, and calculated as the mean log₁₀ RPKM across all samples. For each disorder, the gene list was sorted by one property at a time, and genes were removed in parallel from the two disorders: at each step, the gene with either the highest or the lowest value was removed from each set, ensuring an equal number of genes were removed from both gene sets throughout. After each removal step we recomputed the cell type mutation biases for ASD without ID and SCZ and calculated their Spearman’s correlation across all neuronal cell type clusters. As a matched control, we additionally performed 1,000 iterations of gene removal in random order and computed the corresponding distribution of correlations at each removal step.

### Difference in cell type mutation biases between disorders

To identify disorder-specific mutational biases at the supercluster level, we assessed differences in mutation bias between disorders (ASD w/o ID and SCZ), quantified by the average difference in mutation biases across the cell type clusters comprising each supercluster. This differential bias measure, which describes how strongly a given supercluster is preferentially affected in one disorder relative to the other, was calculated as follows:

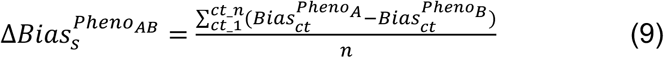

where 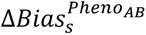 is the differential bias of supercluster *s* between phenotype A and B. 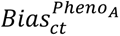 is the mutation bias for a cell type cluster in supercluster *s* for phenotype A, and 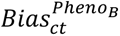 is the corresponding mutation bias for the cell type cluster in supercluster *s* in phenotype B. The summation is over all *n* cell type clusters within the supercluster.

To evaluate whether these differential biases were statistically significant, we applied the Mann-Whitney U test to compare the distributions of cell type mutation biases between disorders for each supercluster (e.g., CGE interneurons, MGE interneurons, and medium spiny neurons). Multiple hypothesis correction was performed using the Benjamini-Hochberg procedure to control the false discovery rate across the 31 superclusters. Between-disorder comparisons included mutation-bias estimates from all clusters within each supercluster, regardless of whether their biases were statistically significant within either disorder.

### Animals used in this study

All experiments were carried out in accordance with the animal welfare guidelines of the Institutional Animal Care and Use Committee of Columbia University. The *Df(16)A^+/-^*line was developed in house^81^. For slice electrophysiological targeting of VIP+ interneurons, we crossed the *Df(16)A^+/-^*line with a VIP-eGFP line to clearly visualize VIP+ interneurons (MMRRC strain #31009, STOCK Tg(Vip-EGFP)37Gsat, University of California, Davis, CA, USA). For all *in vivo* calcium imaging experiments, we crossed the *Df(16)A^+/-^*line with VIP-Cre mice (JAX strain #010908, STOCK *Vip^tm^*^1^*^(cre)Zjh^*/J, The Jackson Laboratory, Bar Harbor, Maine, USA). Mice were held in a 12-hour light/dark cycle vivarium and housed 2-5 mice in each cage. All experiments were conducted during the light portion of the cycle. All mice used in the *in vivo* imaging experiments were male (n = 8), with (n = 4 WT, n = 4 *Df(16)A^+/-^*). Slice experiments included n = 3 WT and n = 4 *Df(16)A^+/-^*mice. All wild-type mice were littermate controls of the *Df(16)A^+/-^*mice.

### Hippocampal slice electrophysiological recordings

8-12 -week old *Df(16)A^+/-^*;*VIP-eGFP^+^*mice were anesthetized with isofluorane and transcardially perfused with ice-cold N-methyl-d-glucamine (NMDG) artificial cerebrospinal fluid (NMDG-ACSF, containing 90 mM NMDG, 2.5 mM KCl, 1.2 mM NaH2PO4, 30 mM NaHCO3, 20 mM HEPES, 25 mM glucose, 5 mM sodium ascorbate, 2 mM thiourea, 3 mM sodium pyruvate, 10 mM MgSO4, 0.5 mM CaCl2, 93 mM HCl) that was bubbled with 5% CO2–95% O2. The brain was extracted and sectioned into 300-µm-thick coronal slices using a vibratome (VT1200, Leica Microsystems) in ice-cold oxygenated NMDG-ACSF. Coronal sections including CA1 were bubbled in the same solution at 37 °C for 8 min and transferred to oxygenated modified-HEPES ACSF at room temperature (20–25 °C; 92 mM NaCl, 2.5 mM KCl, 1.2 mM NaH2PO4, 30 mM NaHCO3, 20 mM HEPES, 25 mM glucose, 5 mM sodium ascorbate, 2 mM thiourea, 3 mM sodium pyruvate, 2 mM MgSO4, 2 mM CaCl2) for at least 1 h before recording. Recordings were performed in a submerged chamber, superfused with continuously bubbled ACSF (125 mM NaCl, 2 mM KCl, 1.25 mM NaH2PO4, 25 mM NaHCO3, 11.102 mM glucose, 2 mM CaCl2, 1 mM MgCl2) at near-physiological temperature (34 ± 1 °C). For whole-cell current-clamp recordings, the electrodes (4–6 MΩ) were filled with an intracellular solution containing 135 mM K-gluconate, 4 mM KCL, 10 mM HEPES, 10 mM phosphocreatine, 4 mM Mg-ATP, 0.3 mM GTP-Na, and 2 mM Na2-ATP. Current-clamp responses were recorded with a Multiclamp 700B amplifier (filtered at 2–4 kHz) and digitized at 10 kHz (Digidata 1440). Series resistance was always <20 MΩ and was compensated at 80– 95%. Transgenically expressed eGFP signal was visualized using infrared differential interference contrast and fluorescence video microscopy (Olympus; CoolLED) with a ×40 objective. eGFP+ cells were targeted in *stratum pyramidale*.

### Electrophysiology recording analysis

Data were collected and analyzed off-line using the AxographX v1.7.6. The action potential frequency was measured during holding currents of 0–250 pA. The membrane voltage at the end of the current clamp was measured and plotted against each holding current. This allowed us to see the resting membrane potential, the input resistance, and the membrane time constant. All measurements were averaged across 19 neurons in *Df(16)A^+/-^*mice and 11 neurons in wildtype controls. The resting membrane potential was measured while cells were held at 0 pA. Rheobase was measured as the smallest current needed to depolarize the cell membrane. Initial spike latency was defined as the time from the onset of holding current change to the first measurable deflection of the potential from baseline. The firing threshold was defined as the voltage where first deflection of the potential from baseline was measured. Half-width was defined as the width between 10% and 90% of the action potential. Rise time and decay time were defined as the time between 10% and 90% of the rise or decay of the action potential, respectively. The membrane time constant was defined as the time it took the voltage level of the cell to decay to a resting state after delivering a 200 ms step of −20 pA. Ih (hyperpolarization-activated current) sag was quantified as the difference between the minimum peak voltage and the steady-state plateau voltage (sag ratio). Afterhyperpolarization (AHP) was measured by finding the difference between the resting potential and the most negative point (peak) of the AHP following an action potential. Input resistance was measured using ΔV/pA = mΩ for each current step and averaging across all current steps for individual neurons. Upstroke and downstroke velocities were measured as the maximum and minimum dV/dt, respectively, of the action potential using the Allen Institute ipfx package (https://github.com/AllenInstitute/ipfx), from the 1st AP evoked at Rheobase + 20 pA. All other relevant measurements of action potential characteristics were done analyzing the 1^st^ AP of Rheobase. All measurements were averaged across neurons.

### Viruses

Cre-dependent recombinant adeno-associated virus (rAAV) for GCaMP7f (rAAV1-syn-FLEX-jGCaMP7f-WPRE, Addgene #1944820-AAV1, titer: ≥1:10^13 vg/mL) and GCaMP8m ((rAAV1-syn-FLEX-jGCaMP8m-WPRE, Addgene # 162378-AAV1, titer: ≥1:10^13 vg/mL) were used to express GCaMP7f or GCaMP8m in the right, dorsal hippocampus of VIP-Cre mice crossed with Df(16)A^+/-^mice. The Cre-dependent recombinase enabled expression of calcium sensor GCaMP7f or GCaMP8m in VIP+ interneurons.

### Viral injections and hippocampal window/headpost implant

During isoflurane anesthesia, Df(16)A^+/-^x VIP-cre mice were placed onto a stereotaxic surgery instrument. Meloxicam and bupivacaine were administered subcutaneously to reduce discomfort and provide analgesia. The skin was cut along the midline to expose the skull, and a craniotomy was performed over the right hemisphere with a dental drill. A Nanoject injector was loaded with a sterile glass capillary filled with mineral oil and rAAV1-syn-FLEX-jGCaMP7f-WPRE or rAAV1-syn-FLEX-jGCaMP8m-WPRE and lowered into dorsal CA1 of the right hemisphere. Injections were made at AP -2.2, ML -1.75, DV - 1.8, -1.6, -1.4, -1.2, -1.1, -1.0, -0.8 relative to bregma with 64 nl of virus injected at each DV level. The pipette was held within the brain for 5 minutes after the last injection and slowly raised from the brain. The skin was sutured, and a topical antibiotic was applied to the suture site. 1.0 mL of saline was subcutaneously given to each mouse, while the mice recovered in their home cage with a heat pad. Four days after viral injection, a circular cannula with a glass window was inserted into the cavity produced by craniotomy as previously described^117^. Similarly as described above, mice were anesthetized with isoflurane and placed on a stereotaxic surgery instrument. Meloxicam and bupivacaine were subcutaneously given to the mice. The skull was exposed, and a 3 mm craniotomy was performed over the right hippocampus at AP -2.2, ML -1.75 relative to bregma, centered around the injection site. After removal of the dura, the cortex was slowly aspirated with negative pressure and the consistent application of ice-cold cortex buffer. When the horizontal fibers over CA1 were exposed, aspiration was stopped and a sterile collagen membrane was applied to stop bleeding. A 3 mm circular cannula with a glass window was placed into the craniotomy and glued into the site with Vetbond. A headpost was then attached to the posterior of the skull with layers of dental cement. 1.0 mL of saline was subcutaneously given to each mouse, while the mice recovered in their home cage with a heat pad. All mice were monitored for 3 days of post-operative care until behavior training began.

### Behavioral training and paradigms

Following the post-op recovery period, mice were acclimated to handling and head-fixation for a few days, while being water restricted to 85-90% of their original weight. Mice were trained to run on a fabric, cue-free burlap belt with random rewards that were automatically dispensed at a decreasing number of random locations. Mice were trained for ∼1 week on the cue-free belt and then were transitioned to a cue rich belt during the imaging experiments.

### Random Foraging Task

Water restricted mice were imaged on a cue rich familiar belt for 2 days with 3 random rewards distributed across the length of the belt, randomized every lap.

### AOD-based two-photon calcium imaging

After the mice were acclimated and trained in the respective tasks, a high-resolution structural Z-stack was captured 24 hours prior to imaging, while the mouse was anesthetized with isoflurane. The reference Z-stack was used to identify the 3D position of the GCaMP-positive neurons for access by the custom-modified AOD (acousto-optic deflector) microscope (Femto3D-ATLAS, Femtomics Ltd). The AOD microscope was used as previously described in Geiller et al. 2020, and Vancura and Geiller et al. 2023.

To provide stable transmission parameters during chronic imaging in the entire 3D scanning volume, the AOD microscope was extended with a high-speed and precision beam stabilization unit, which was directly attached to the AOD scan head, sensitive to input beam misalignment. The beam stabilization unit consisted of two quadrant detectors (PDQ80A and TPA101, Thorlabs) and two broadband dielectric mirrors (Thorlabs) mounted on motorized mirror mounts (Femtonics). Beam alignment was performed by the LaserControl software (Femtonics).

A water immersion objective (x16 Nikon CF175) was used to focus at the dorsal CA1 pyramidal layer and then fixed in position. A tunable two-photon laser (Coherent Ultra II) was adjusted to wavelength = 920 nm. The high-resolution structural Z-stack was captured in CA1 from ∼100 um below to glass to ∼500 um to span the entirety of CA1 from the *stratum oriens* to the *stratum lacunosum-moleculare* with a 800x800 pixel capture with a resolution of 1.25 um per pixel with a 4 um step size. Laser power and photomultiplier tube (PMT) detectors (GaAsP, H10770PA-40, Hamamatsu) were compensated in power and gain throughout the Z-stack capture. After the Z-stack was completed, the mouse was returned to its home cage and given 24 hours to recover before functional imaging.

Using the integrated software from Femtonics (MES), the Z-stack was examined, and 100-150 interneurons were manually selected to generate a spreadsheet of the x-y-z positions at the center of each respective cell. These x-y-z positions served as the regions of interest (ROIs) that were then used on subsequent days by the AOD microscope to find the position of the corresponding interneurons. For each day of functional imaging in each mouse, the identical field of view was identified within the CA1 pyramidal layer and then the ROIs were loaded in using the software for AOD 3D imaging. Each imaging session was taken for 10 minutes, with approximately 5-10 Hz for each recording experiment (frame rate depended on the number of ROIs, ROI size, imaging resolution, and scanning speed).

### Immunohistochemistry

The immunohistochemical (IHC) staining strategy is consistent with our previously published methods^86,89^. In brief: mice were collected the last day after imaging and perfused with 20 mL Phosphate Buffered Saline (PBS) followed by 20 mL of 4% paraformaldehyde (PFA, Electron Microscopy Sciences) and the brain extracted for overnight fixation in 4% PFA. The next day, the brains were washed 3 x 5 minutes with 1x PBS and subsequently vibratome sliced for horizontal sections from the dorsal side of the brain into 75 µM sections. The relevant sections were then taken through our IHC protocol. Sections were permeabilized for 2 x 20 minutes (0.03% Triton-X 100 (Sigma-Aldrich), 1x PBS), blocked for 45 minutes (0.03% Triton-X 100, 10% Normal Donkey Serum (NDS), 1x PBS), and incubated with primary antibodies (Table 1) for 1 hour at room temperature and overnight at 4 degrees with agitation. After 48 hours, primary antibodies were washed out (3 x 5 minutes 1x PBS) and secondary antibodies were applied for 2 hours (Table 1). Secondary antibodies were removed and the sections were washed with 1x PBS 5 x 15 minutes, and the sections were mounted for confocal imaging.

**Table 1.** Antibodies used in this study.

| Cell Marker | Round | Primary Antibody (dilution) | Secondary Antibody (dilution) | Catalog # |
| --- | --- | --- | --- | --- |
| Cholecystokinin (CCK) | 1 | rabbit anti-proCCK (1:500) | donkey anti-rabbit DyLight 405 (1:300) | Frontier Institute CCK-pro-Rb-Af350 |
| Calretinin (CalR) | 1 | Guinea pig anti-Calretinin (1:1000) | donkey anti-guinea pig Alexa 568 (1:300) | Swant CRgp7 |
| Muscarinic Cholinergic M2 Receptor (M2R) | 1 | rat anti-M2R (1:2000) | donkey anti-rat Alexa 647 (1:300) | Millipore MAB367 |

### Confocal imaging

The NikonA1 confocal microscope used to take multi-channel fluorescence images (405 nm, 488 nm, 561 nm, and 640 nm excitation for the four blue, green, red and far-red colored channels) of labeled tissue sections. Image capturing yielded 2048 x 2048 pixel z-stacks of the entire depth of the section (with 3 microns between each image of a given stack). Z-stacks were viewable via Fiji (Image J).

### Registration of confocal images to *in vivo* z-stacks and identification of immunopositivity/negativity

Unique groupings of cells from *in vivo* z-stacks were used as visual fiducial markers to manually register *ex vivo* 75 μm z-stack images in ImageJ, section by section. Care was taken to note the orientation of the *ex vivo* sections regarding the *in vivo* z-stack images. The coordinates taken from the AOD software when selecting ROIs for imaging against the *in vivo* z-stack were used to identify ROI ID numbers in the registered immunohistochemically stained images.

### Assignment of subtype identity based on immunostaining

Interneuron subtypes were assigned based on our previously published classification strategy^86^.

#### Cholecystokinin-expressing basket cells (CCKBCs)

GCaMP-expressing, immunopositive for proCCK, immunonegative for CalR and M2R^118,119^. This population represents perisomatic-targeting basket cells.

#### Interneuron-selective interneuron type 2 (ISI2s)

GCaMP-expressing, immunonegative for CalR, CCK and M2R. Present in *stratum radiatum* or *stratum lacunosum moleculare* CA1 layers^118,119^.

#### Interneuron-selective interneuron type 3 (ISI3s)

GCaMP-expressing, immunopositive for CalR and immunonegative for CCK and M2R. Present in all CA1 layers^106,118–121^.

*VIP only (VIPos):*

GCaMP-expressing, immunonegative for CalR, CCK and M2R. Present in *stratum oriens* or *stratum pyamidale* CA1 layers. This category excludes CCKBCs, ISI2s, and ISI3s, and could possibly include long-range projecting interneurons^122^, but it may not be restricted to them; therefore we called them VIP only for our purposes in this study.

### Cross-species CGE interneuron subtype mapping

The mouse VIP+ subtypes examined in our *Df(16)A^+/−^* experiments (CCKBC, ISI2, ISI3, VIPo) were defined by immunohistochemical markers and hippocampal layer position. Several of these distinctions have no direct counterparts in human cortical transcriptomic atlases: the ISI2/VIPo distinction relies entirely on hippocampal layer position, and CCK mRNA is broadly expressed across human cortical neurons^123^, as confirmed in the Siletti et al. atlas^37^. We therefore restricted the cross-species comparison to transcriptionally defined putative CCKBCs and other VIP+ CGE clusters. Mouse CCKBCs were operationally defined as the Sncg subclass together with the Vip Sncg and Vip Serpinf1 supertypes, following the integrated morpho-electric-transcriptomic (MET) classification of Scala et al.^124^. Per-cell transcriptomic type assignments were obtained from published Patch-seq metadata: Gouwens et al.^125^ for primary visual cortex (V1; 282 putative CCKBCs among 1,556 CGE cells) and Scala et al.^124^ for primary motor cortex (M1; 51 putative CCKBCs among 257 CGE cells). To identify candidate human CCKBC-like clusters, we mapped the 1,813 mouse CGE Patch-seq cells onto a human reference comprising the interneuron and control superclusters of the Siletti et al. atlas^37^ using Harmony^126^ and scVI^127^. Harmony mapping used inverse-distance voting among 30 human reference neighbors in the corrected PCA embedding. scVI query mapping used 30 candidate human cluster centroids and 100 global human reference neighbors. Voting was restricted to neighbors within candidate clusters when at least 10 such neighbors were available; otherwise, all global neighbors were used. Votes were weighted by inverse distance. For each of the 21 human CGE clusters (276–296), we calculated the fraction of assigned mouse CGE cells carrying the putative CCKBC label, separately for M1 and V1 (Figure S15A; Table S8). This fraction was left undefined for clusters with no assigned cells.

To assess within-species transcriptomic correspondence, we also mapped M1 Patch-seq cells onto mouse cortical GABAergic reference cells from the Allen Brain Cell (ABC) Atlas^128^ using Harmony and inverse-distance voting among 30 reference neighbors. Each query cell was assigned to an ABC GABAergic subclass, providing a within-mouse control for the transcriptomic mapping. We independently evaluated cross-species correspondence using MetaNeighbor^129^. Mouse CGE/MGE GABAergic cells from the WMB-10Xv3 Isocortex expression files were compared with human CGE clusters 276–296. Input cells were sampled at a maximum of 50 cells per cluster before restricting mouse cells to cortical CGE/MGE types. We used one-to-one orthologs and selected the 3,000 genes with the highest variance across the combined input cells. Expression values were ranked within each cell, then centered and normalized before computing cross-species voting scores. We used the human-as-test direction: for each human cluster, we selected the mouse cluster whose scores best distinguished the human cluster from other human CGE cells, as measured by AUROC, and reported its mouse subclass (Table S8). We additionally examined pseudobulk expression of CCKBC-associated and discriminating markers (CCK, CNR1, CXCL14, CRH, SNCG, VIP, CHRM2 and RELN; Figure S15B). Candidate assignments were evaluated using the combined evidence from transcriptomic mapping, MetaNeighbor correspondence, and marker expression, as detailed in the Supplemental Text.

To assess electrophysiological correspondence, we compared electrophysiological properties across four groups defined by transcriptomic labels: putative mouse CCKBC cells (n = 50; M1), mouse VIP-other cells (n = 112, M1), human VIP-SERPINF1 CCKBC candidate cells (n = 2 with usable measurements; human interneuron Patch-seq dataset^130^), and human VIP-other cells (n = 100). Human candidate cells carried the transcriptomic label Inh L2-5 VIP SERPINF1; other human VIP cells were required to have both a VIP subclass label and a VIP transcriptomic-type label. We used published electrophysiological measurements from the Scala et al. mouse and the Lee/Dalley human feature tables. Missing values were excluded separately for each feature, and groups were compared within species using two-sided Mann–Whitney U tests (Figure S15C; Supplemental Text).

### Pre-processing of Ca2+ imaging data

Raw movies of each individual cell were independently motion corrected using the whole-frame cross registration implemented in the SIMA package^131^. ROIs were then hand-drawn over each imaged cell. Fluorescence was extracted from each drawn ROI using SIMA. DeltaF/F was then computed based on a 1st percentile baseline value over a 30-second rolling window. The DeltaF/F traces were then smoothed through an exponential filter. For analyses involving velocity regressed traces, each animal’s velocity was smoothed with a Hanning filter and then covariate linear regression was applied to each fluorescence trace where the smoothed velocity is regressed out.

### Velocity Correlation

For each cell, velocity correlation was computed by taking Pearson’s correlation coefficient between the DeltaF/F trace and animal’s smoothed velocity with a -5 second to +5 second rolling lag. The highest absolute value of the correlation coefficients in this time window was taken as the correlation coefficient for the respective cell.

### Peri-Event Time Averages (PETA)

For salient stimuli such as reward distribution events, a PETA was constructed around each event respectively. These events were z-score and then averaged for each respective cell, and the z-scored averages were visualized using a heatmap or an averaged line-plot. A Post-Pre metric was calculated based on the z-scored averages as a measure for event modulation.

### Spatial Tuning Curve

The 2 meter belt was divided into 100 bins (∼2cm per bin). Spatial tuning curves for each cell during an experiment were generated by averaging the velocity regressed DeltaF/F in each bin, when the animal was running over 5 cm/s.

### Place field calculation

The spatial tuning curves were shuffled 1000 times by circularly rotating the velocity-regressed ΔF/F tuning curves around the 100 spatial bins to generate a shuffle distribution. Cells were classified as being spatially selective (i.e., having a place field) if the cell’s tuning curve exceeded the 95th percentile (or was under the 5th percentile for negatively selective cells) with at least 10 sequential bins over/under the cutoffs from the shuffled distribution. The number of bins over or under the 95th and 5th percentile curves were visualized and a cutoff of 30 consecutive bins was used for the figures as it represented the approximate 75th quartile for place field width.

### Position Decoding

A linear classifier support vector machine (SVM) was trained on the tuning curves of each respective cell and the decoding error was visualized. The SVM was cross-validated by training the classifier on (n-1) laps and evaluating on the held-out lap and this process was repeated such that every lap was utilized as a test lap at least once. Furthermore, the SVM was bootstrapped with different numbers of input cells and input laps to compare the position decoding efficacy between genotypes.

### Cross-correlation Analysis

Pairwise Pearson’s correlation coefficients were computed for every pair of cells within a given experiment. The average coefficient value for each subtype was then calculated per trial pair and averaged across mice.

### Transient Analysis

Online Active Set method to Infer Spikes (OASIS) was used to generate deconvolved spike events based on transient events within the dF/F traces^98^. We understand that these are not true spiking events, and we are using OASIS to algorithmically identify transient events rather than interpret the deconvolved spiking events.

#### Frequency

The deconvolved events were filtered to remove events that were within 10 frames. The total number of filtered deconvolved events by OASIS was normalized by the total duration of the recording.

For each cell, the average transient was segmented within a 15-second time window and computed by averaging along aligned deconvolved spike times. If multiple detected events were within 10 frames, the events were treated as a single transient. A cell was only used if there were at least 3 detected transients within the total trace to exclude cells without obvious GCaMP-Ca^2+^ dynamics. For each average transient, we computed the median value based on 5 seconds pre peak transient and the range 5-10 seconds after the peak transient (given that the average transient took significantly less than 5 seconds to resolve) to act as the baseline.

#### Amplitude

The difference between the max value of the average transient and the baseline value was computed

## QUANTIFICATION AND STATISTICAL ANALYSIS

### Statistical analyses

Subtype comparisons were conducted using one-way ANOVA tests with Tukey’s range tests with correction for multiple testing if appropriate. For comparisons across genotypes, unpaired t tests were utilized if the data followed a normal distribution (Shapiro-Wilk and Kolmogorov-Smirnov, where applicable). Otherwise, for non-normally distributed comparisons, the Mann-Whitney U test was used. *, P < 0.05, **, P < 0.01, ***, P < 0.001. All data analysis and visualization were done with custom software built on Python version 2.7.15.

### Code and Data availability

All data used in this report are publicly available. ASD genes and mutations counts can be found in Supplementary table from Zhou et al^2^. SCZ genes and mutation counts can be found in Supplementary table from Singh et al^1^. DD/ID genes and mutations counts can be found in Supplementary table from Kaplanis et al^50^. Genes implicated in UKBB adult cognitive functions can be found in Supplementary table from Chen et al^64^. Genes in 22q11.2 3Mb deletion and 1.5 Mb deletion can be found in Karayiorgou et al^14^. Human brain cell atlas can be found (https://cellxgene.cziscience.com/collections/283d65eb-dd53-496d-adb7-7570c7caa443).

Individual genetics and phenotypic data used in our study from the SPARK project are available through the Simons Foundation (https://base.sfari.org) following the access procedures described in Zhou et al. Gene constraint score from gnomAD can be found at (https://gnomad.broadinstitute.org/news/2024-03-gnomad-v4-0-gene-constraint/). Brainspan expression data can be found at (https://www.brainspan.org/).

The code used for our implementation of the Cell type bias with analysis pipeline, and code to generating all figures and tables, is available on GitHub: (https://github.com/explorerwjy/CellTypeBias_VIP.git).

## Acknowledgements

S.A.H. is supported by a Burroughs Welcome Fund grant and a Brain & Behavior Research Foundation Young Investigator award. E.R. is supported by NIGMSK99GM153720. This work has been supported by the grants NIMHR01MH124923 to D.V. and J.A.G, NIMHR01MH124047 to J.A.G, Stavros Niarchos Foundation to J.A.G., and NIMHR01MH124047 NIMHR01MH124867 NINDSR01NS133381 to A.L. We thank members of the Gogos, Losonczy, and Vitkup labs for fruitful scientific discussions.

## Author Contributions

S.A.H., J.W., J.A.G., A.L., and D.V. conceived the study. J.W. and J.C. performed genetic bias and computational analyses. S.A.H., B.Y.R. and E.R. collected and analyzed experimental data. S.A.H., J.W., J.A.G., A.L., and D.V. wrote the manuscript.

## Supplemental Tables and Figures

**Table S1:**
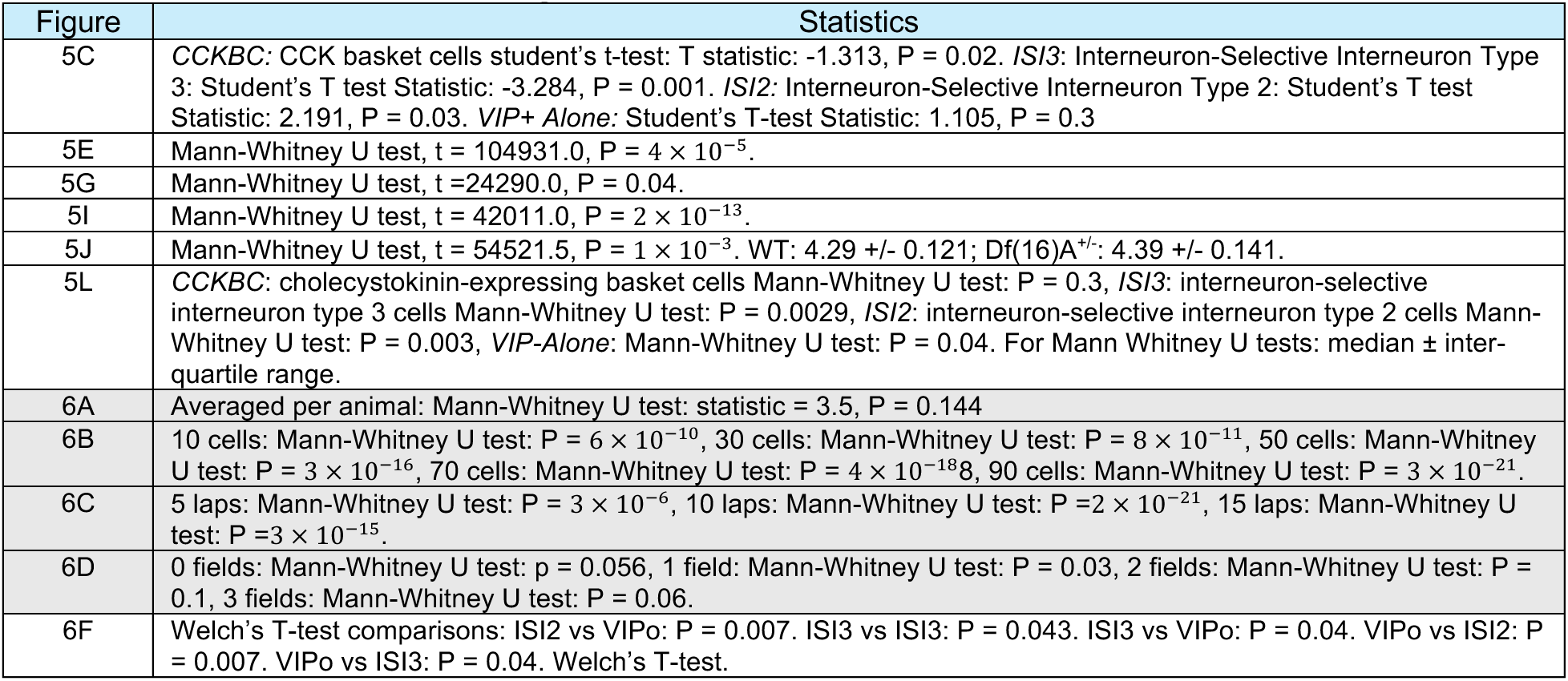
Statistical detail summary. Boxplots show the 25th, 50th (median), and 75th quartile ranges, with the whiskers extending to 1.5 interquartile ranges below or above the 25th or 75th quartiles, respectively. Outliers are defined as values extending beyond. the whisker ranges. For Welch’s t tests: mean ± sem. For Mann Whitney U tests: median ± inter-quartile range. n.s.: not significant. *: P < 0.05. **: P < 0.01. ***: P < 0.001.

**Table S2. Gene set index.** Definitions, sources, weighting methods, and gene counts for the considered disorder and traits gene sets.

**Table S3. Gene-level data and weights.** Gene membership, observed and expected variant counts, and gene weights used to calculate cell type mutation biases.

**Table S4. Cell type mutation biases for disorder and cognitive gene sets.** Mutation Biases, ranks, and empirical *P*-values and FDR-adjusted *q*-values for the 13 considered gene sets across 461 human brain cell types, assessed against random and confounder-matched null models.

**Table S5. Cell type mutation biases for negative controls.** Corresponding results for the six control gene sets representing non-brain traits and cognitive performance.

**Table S6. Supercluster mutation bias comparisons.** Pairwise mutation bias differences, and Mann– Whitney U test results with FDR correction for the five disorder comparisons across neuronal superclusters.

**Table S7. Stability of mutation bias estimates under cohort downsampling.** Correlations with full-cohort mutation bias profiles and gene set overlap for ASD, DD/ID, and SCZ across cohort fractions.

**Table S8. Cross-species evidence for candidate human CCKBC-like clusters.** Harmony/scVI integration, MetaNeighbor similarity, and marker gene expression across the 21 human CGE interneuron clusters, supporting clusters 278–281 as candidate counterparts of mouse cholecystokinin-expressing basket cells (CCKBCs).

**Figure S1:**
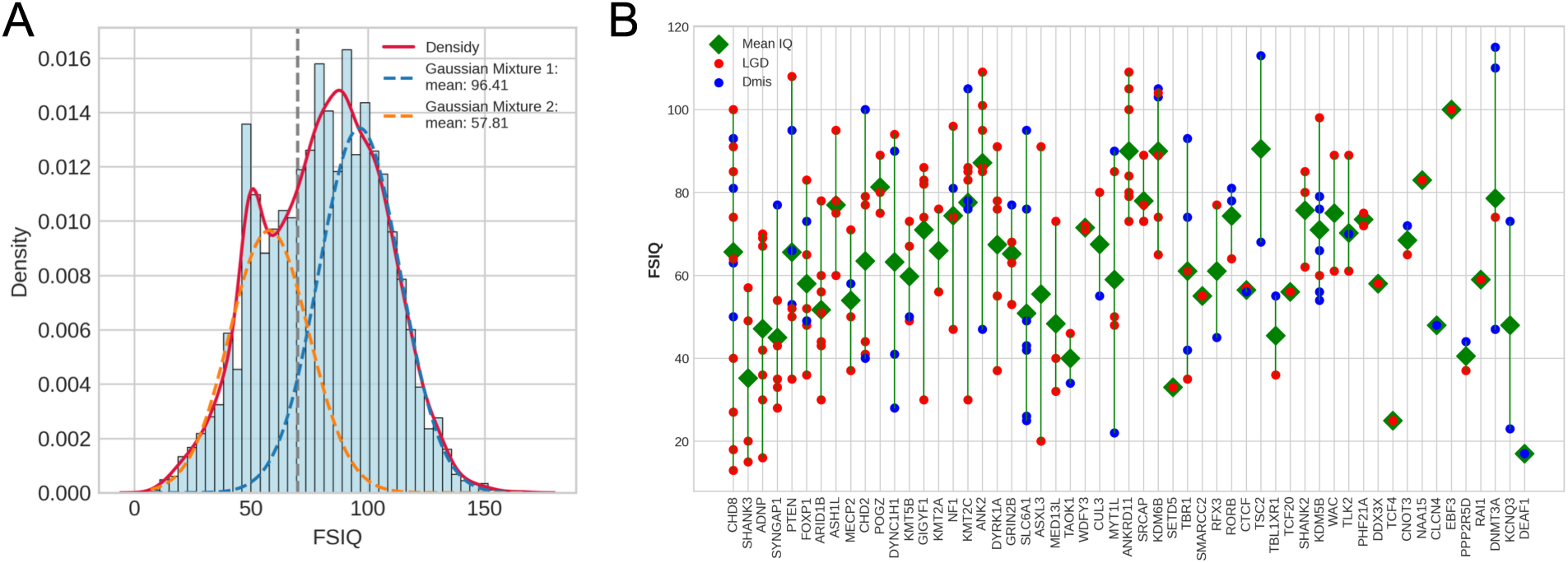
Full-scale IQ distribution among ASD probands in the SPARK and ASC cohorts. **A.** The distribution of FSIQ scores among ASD probands from the SPARK and ASC cohorts. The histogram (blue bars) shows the FSIQ distribution with an overlaid density plot (red line). A two-component Gaussian mixture model was fitted to the data (blue and orange dashed curves), revealing distinct subpopulations within the FSIQ distribution. The vertical dashed line at FSIQ = 70 indicates a commonly used threshold for intellectual disability (IQ < 70). FSIQ phenotype data of SPARK and ASC probands were obtained from SFARI base and Satterstrom *et al*.^62^ **B.** Gene-specific FSIQ patterns across the top 61 high-confidence ASD genes, represented by vertical lines. Individual probands are represented as dots, colored by mutation type, with likely gene-disrupting (LGD) mutations in red and damaging missense (Dmis) mutations in blue. The green diamonds indicate the average FSIQ in the cohorts for each gene. Genes are arranged along the x-axis based on *DenovoWEST* P-values from Zhou *et al*.^2^.

**Figure S2.**
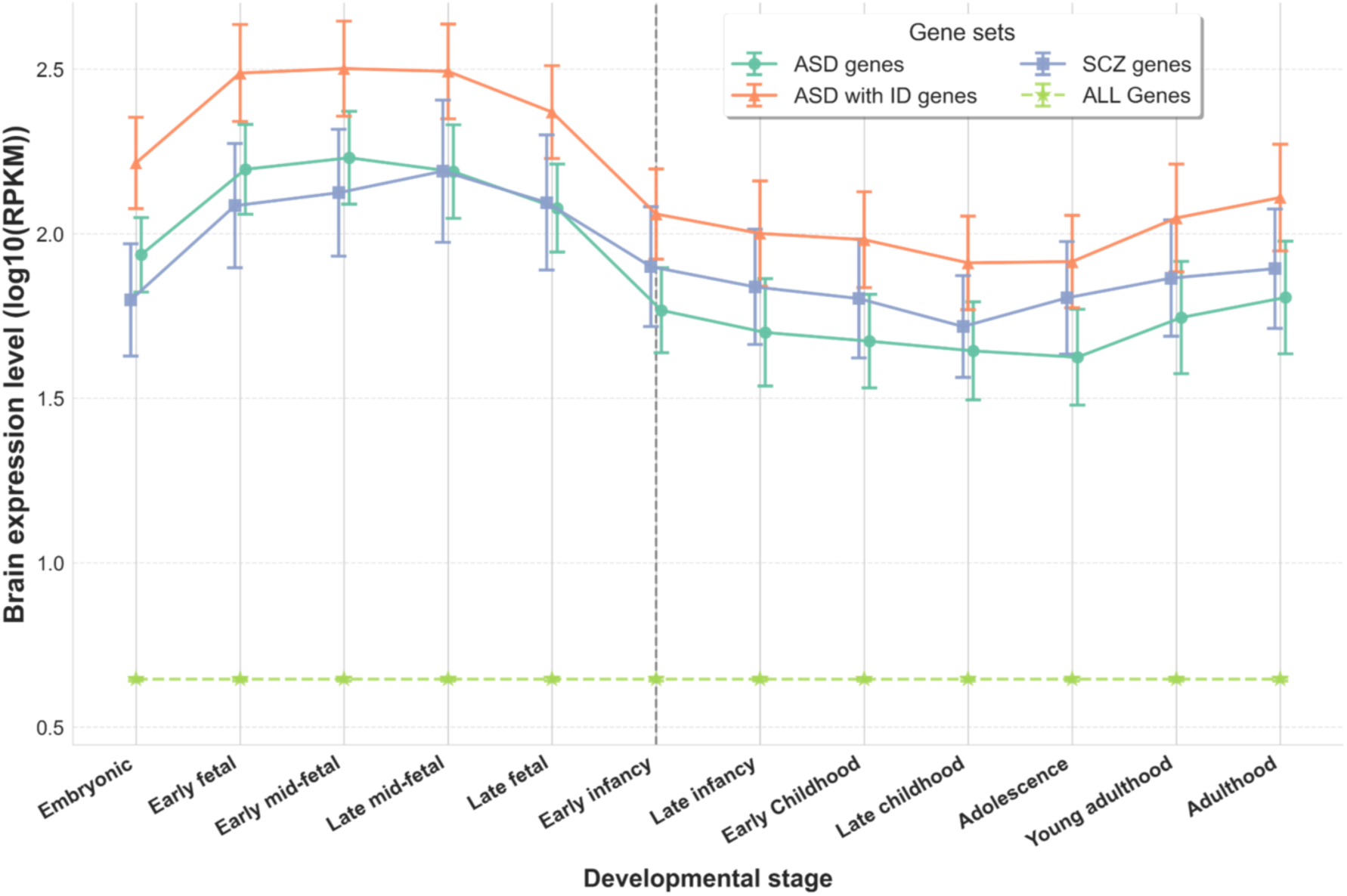
Temporal expression patterns of disorder-associated genes across human brain development stages. Average expression levels (log10 RPKM) of genes associated with different disorders across human brain developmental stages, from embryonic to adult periods. Gene sets include ASD w/o ID genes (green), ASD with ID genes (orange), SCZ genes (blue), and all human genes (dashed yellow). Error bars represent standard error of the mean. The vertical dashed line marks the transition from prenatal to postnatal periods. Expression data were obtained from the BrainSpan Atlas of Human Brain Development^116^.

**Figure S3.**
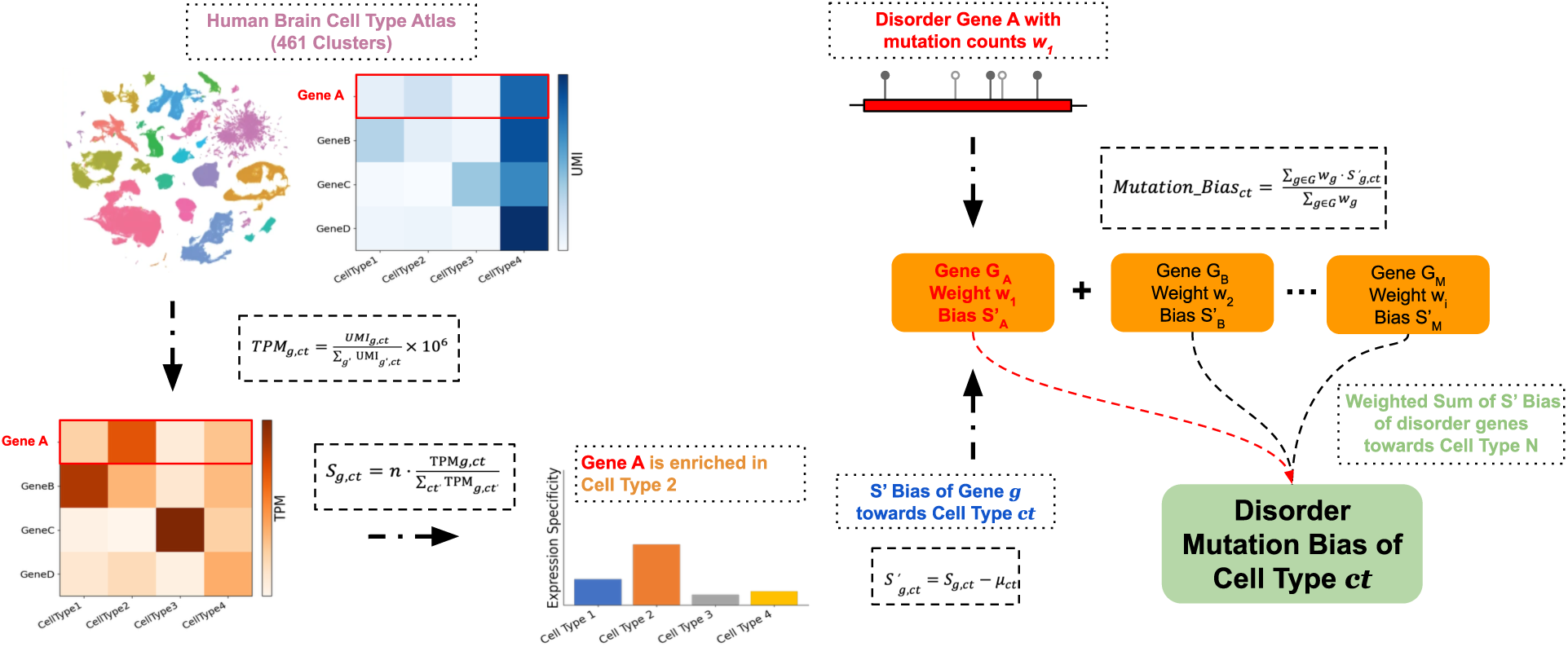
Workflow for computing cell type mutation biases from disorder-associated genes. This schematic illustrates the updated two-step procedure used to compute disorder mutation bias for each brain cell type. **(Left)**: Starting from a comprehensive single-cell gene expression atlas of 461 human brain cell types, we first compute gene expression levels as TPM (transcripts per million) for each gene across all cell types. Expression specificity scores are then derived by normalizing each gene’s TPM in a given cell type relative to its total TPM across all cell types. **(Right)**: For each disorder, cell type–specific mutation bias is calculated by aggregating expression specificity scores across all disorder-associated genes, weighted by their mutational burden. This yields a composite mutation bias score for each cell type, reflecting its relative vulnerability to rare pathogenic mutations in the disorder.

**Figure S4.**
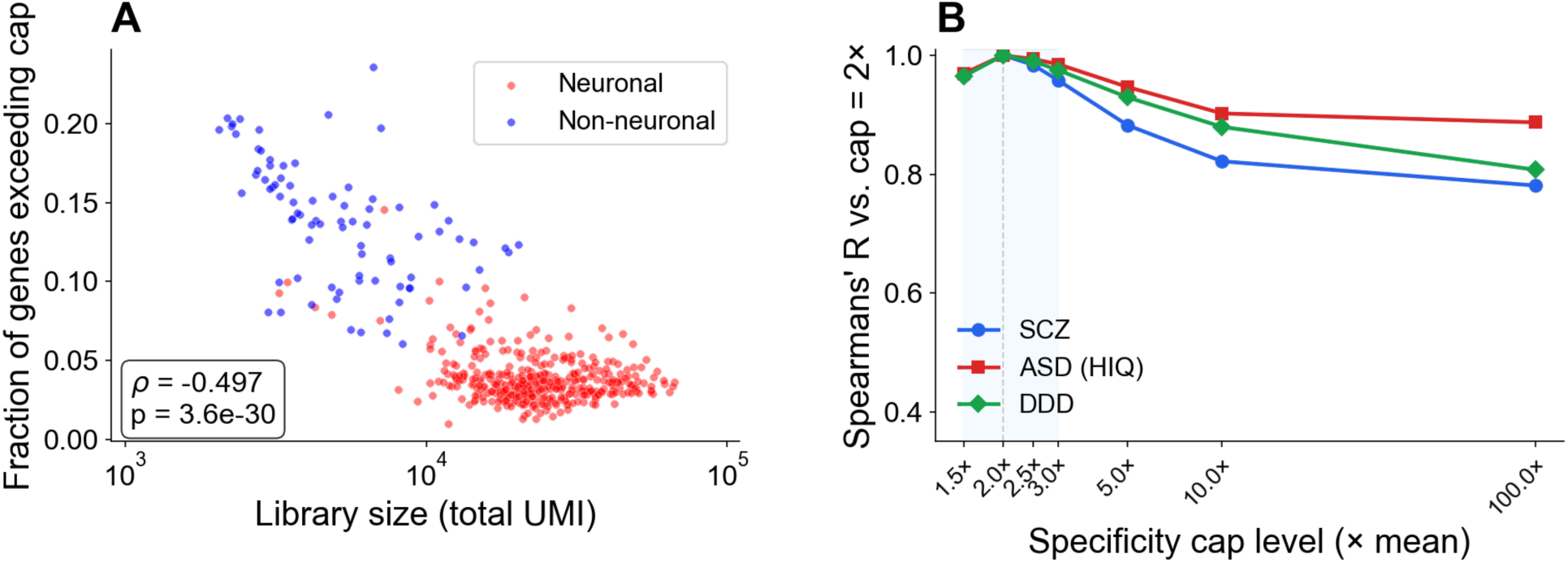
Specificity cap at 2x mean and its effects on cell-type mutation bias rankings. **A.** Fraction of genes with specificity exceeding the 2.0 cap plotted as a function of library size (total UMI) for 461 cell types, categorized into neuronal (red) and non-neuronal (blue) groups. Spearman rho and P-values are indicated. The fraction of genes exceeding the cap was inversely associated with cluster library size, with high fractions occurring predominantly in non-neuronal clusters. **B.** Spearman rank correlation between mutation bias calculated at a 2x cap and a range of alternative cap levels (1.5x to 100x) for SCZ, ASD w/o ID, and DDD. Blue shading indicates the 1.5-3x range.

**Figure S5.**
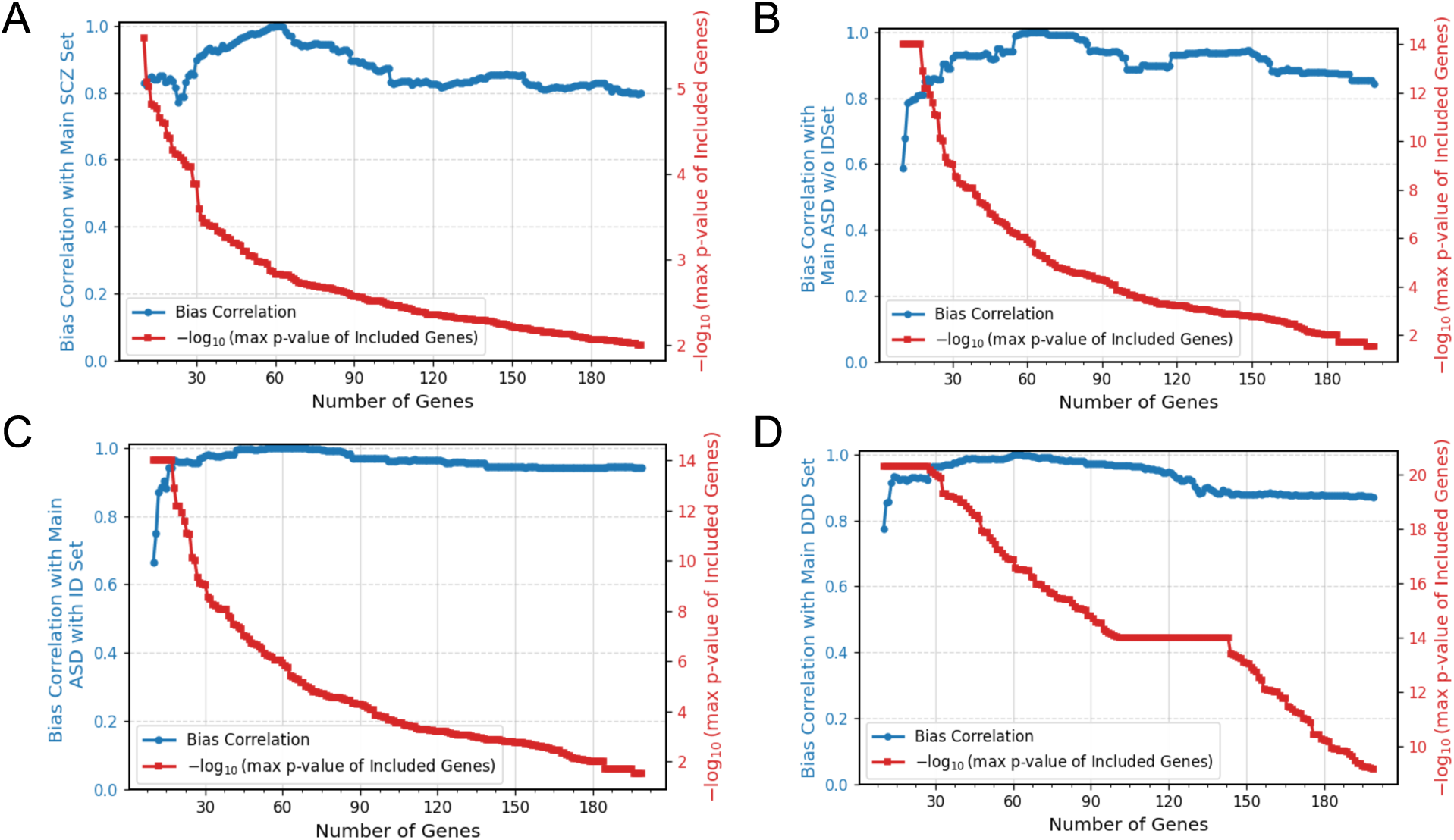
Impact of additional genes included in the mutation bias analysis. Correlations of cell type mutation bias profiles for the primary and expanded gene sets. Mutation bias correlations were calculated between the primary gene set (61 most significant genes) and gene sets of varying sizes (10–200 genes). For each disorder, cell type–specific biases were recalculated while progressively increasing the number of genes included, starting with the most significantly associated and adding progressively less significant genes. Panels show results for: **A.** SCZ; **B.** ASD w/o ID; **C.** ASD with ID, and **D.** DDD. Blue lines indicate the Spearman’s correlation between the mutation bias profile of the expanded set and the primary 61-gene profile. Red lines indicate the –log10 (P-value) of the least significant gene included at each gene set size.

**Figure S6.**
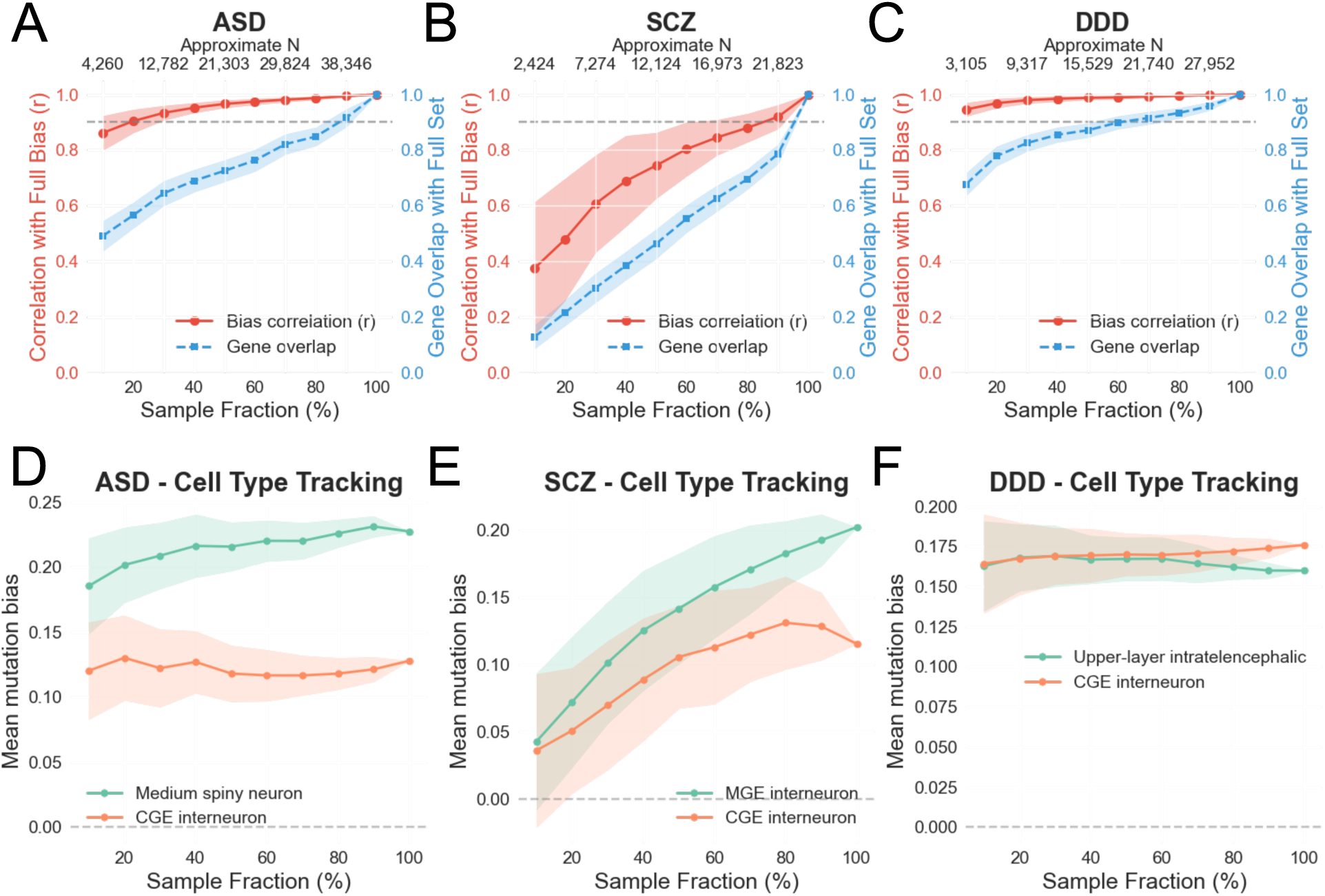
Stability of cell-type mutation bias under cohort down-sampling. **A.-C.** Stability curves for ASD (N = 42,607), SCZ (N = 24,248 cases), and DDD (N = 31,058). Cohorts of each disorder were down-sampled at fractions from 10% to 90% (100 bootstrap iterations per fraction) and recomputed gene discovery and cell-type bias profiles across all 461 cell types (see Methods). Red lines (left axis) show the Pearson’s correlation between the subsampled and full-cohort bias profiles (mean +/-SD across iterations); blue dashed lines (right axis) show the fraction of genes overlapping with the full gene set. **D.-F.** Cell-type tracking panels showing the mean mutation bias (+/-sd) for selected superclusters across sample fractions: **D.** ASD: medium spiny neurons and CGE interneurons; **E.** SCZ: MGE and CGE interneurons; **F.** DDD: Upper-layer intratelencephalic, and CGE interneurons.

**Figure S7.**
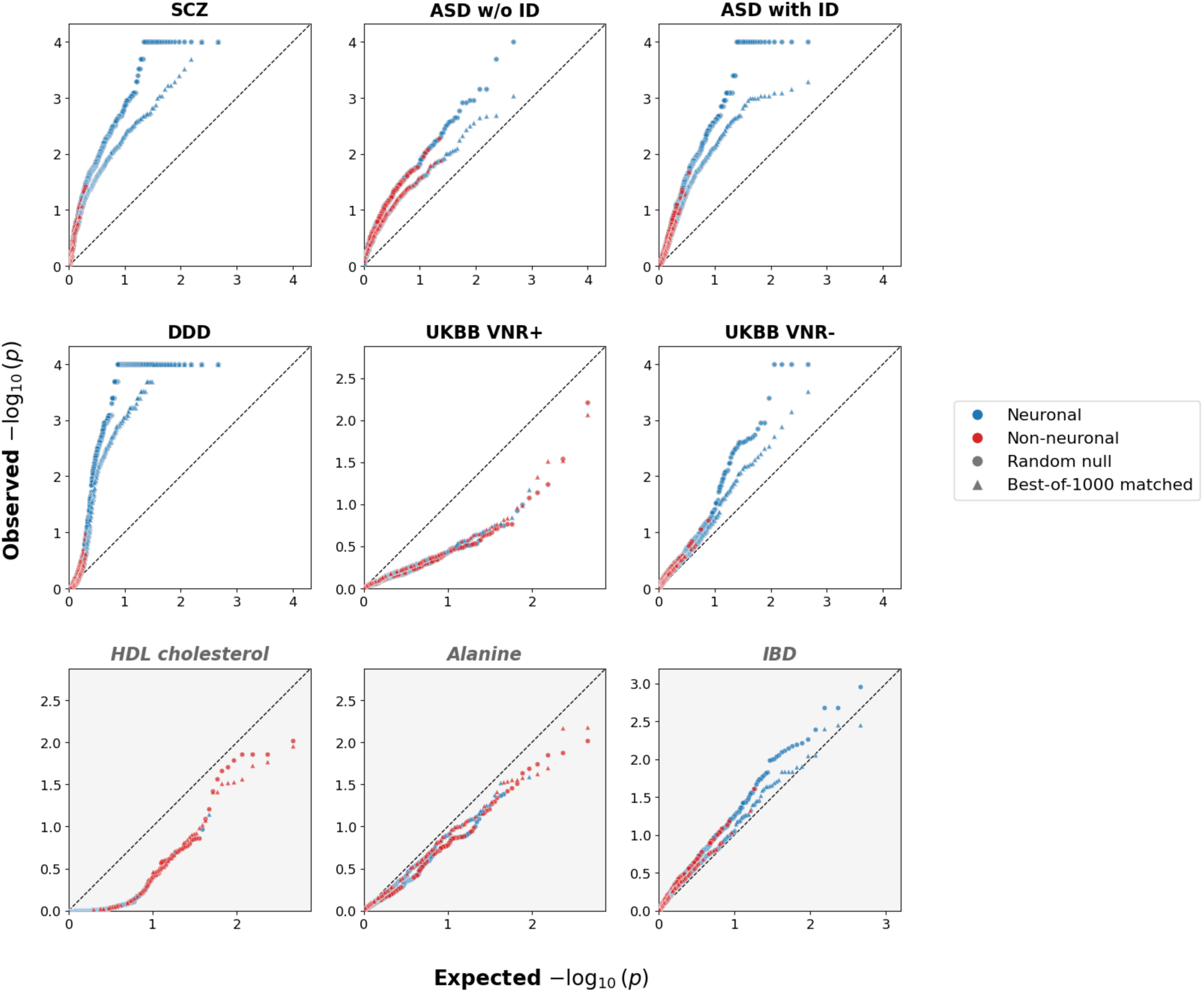
Cell-type mutation bias P-values based on random and confounder-matched gene set null models. Each panel shows a quantile–quantile (QQ) plot of the observed versus expected −log₁₀(P-value) for cell-type mutation bias across 461 cell types. P-values were calculated using 10,000 null gene sets generated under two null models: random gene sampling (circles) and best-of-N confounder matching (triangles, N = 1,000 candidate sets matched for coding-sequence length, whole-brain expression level, and evolutionary conservation). Point colors denote cell class: neuronal (blue) versus non-neuronal (red).

**Figure S8.**
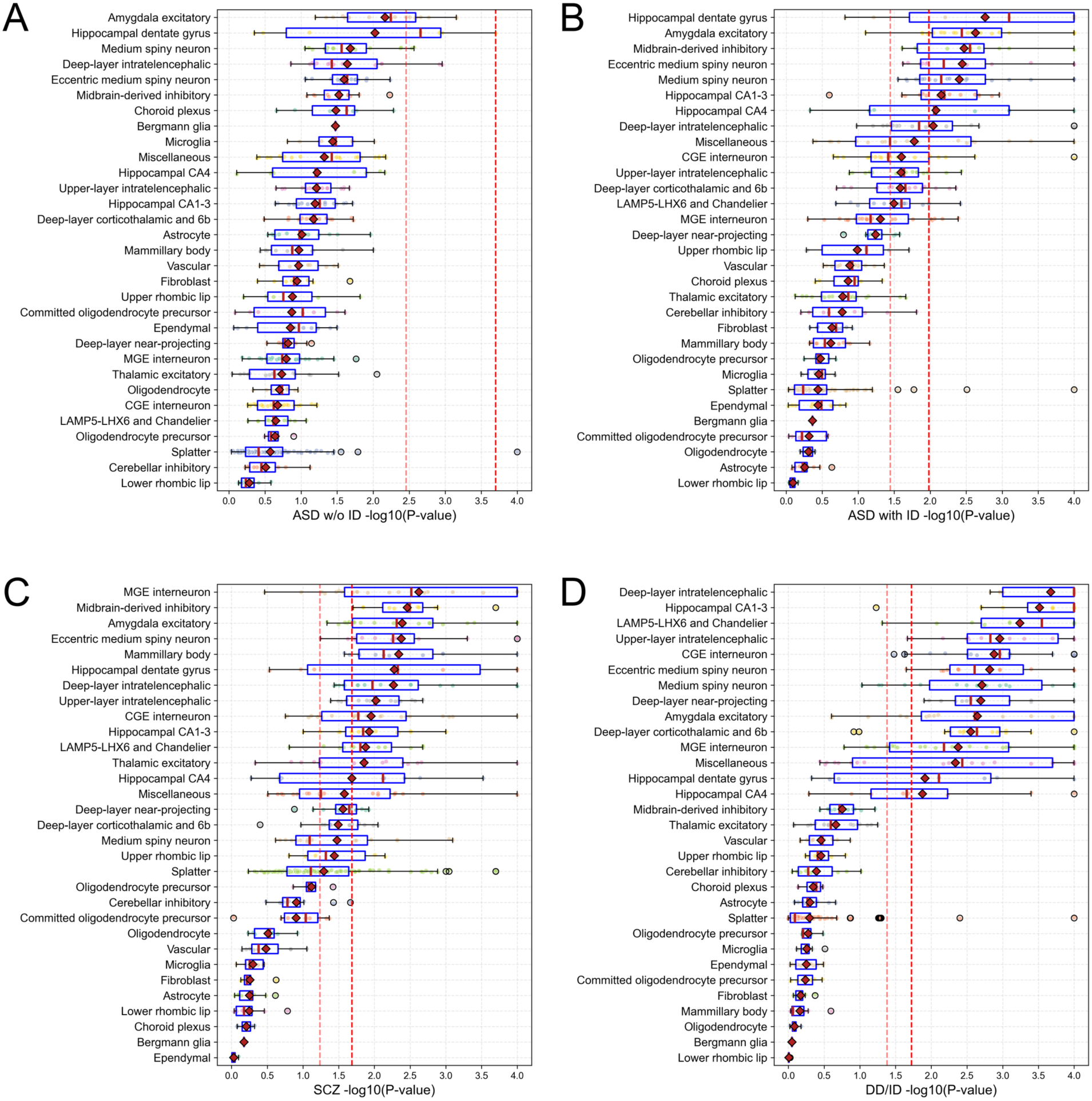
Supercluster-level mutation biases across brain cell types in psychiatric disorders. Box plots show the distribution of mutation bias significance (–log₁₀(P)) for major brain cell type superclusters in: **A.** ASD w/o ID, **B.** ASD with intellectual disability (ID), **C.** schizophrenia (SCZ), and **D.** developmental disorders and intellectual disability (DD/ID). Each point represents a cell type (cluster) within a given supercluster. P-values were calculated by comparing the observed mutation bias of each cluster to a null distribution generated from 10,000 random gene sets (see Methods). Red diamonds indicate the average –log₁₀(P) for each supercluster, and red vertical lines indicate the median. The red dashed line corresponds to an FDR threshold of 0.05; the lighter red dashed line corresponds to FDR = 0.1.

**Figure S9.**
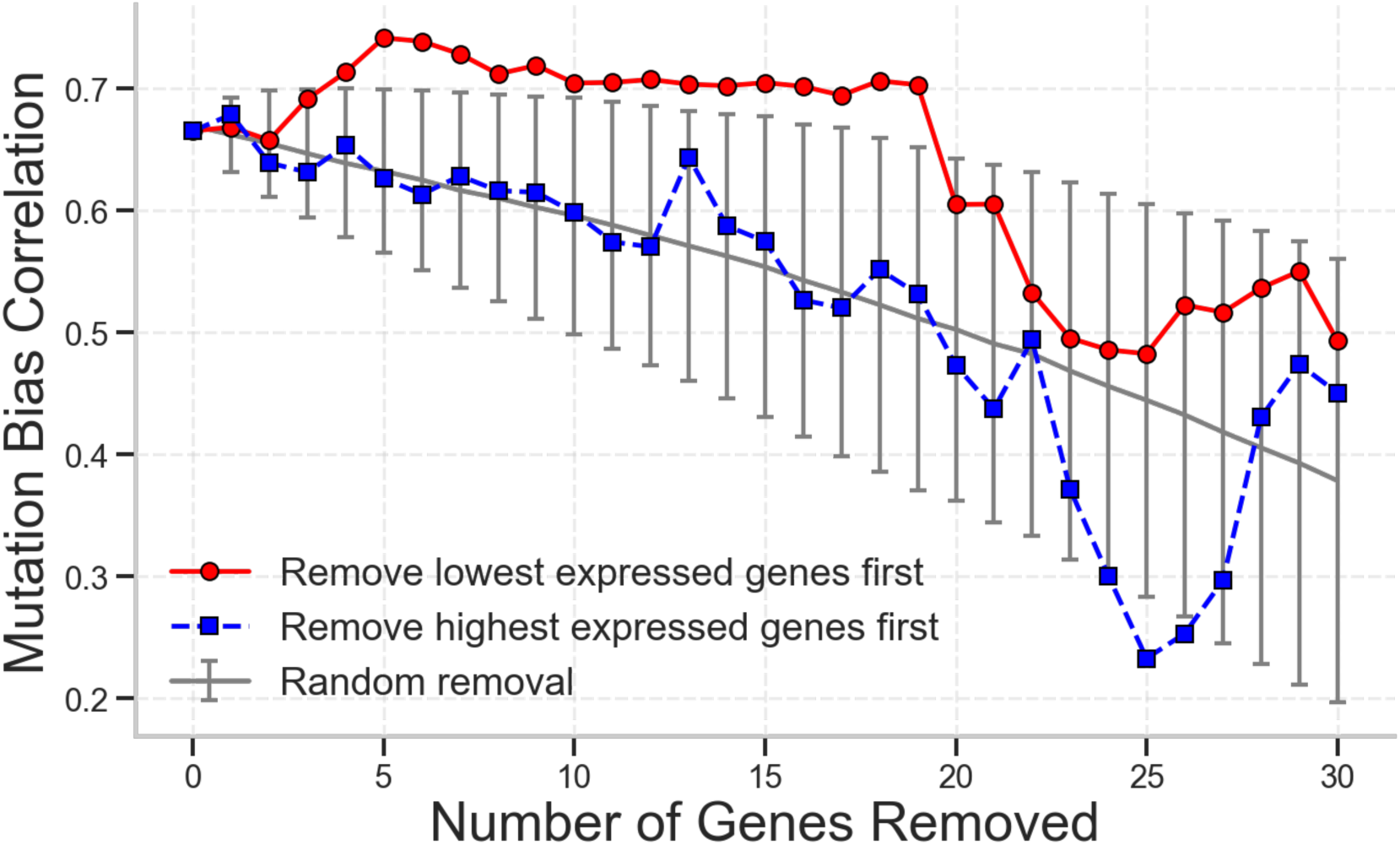
ASD w/o ID–SCZ cell-type bias correlation is robust to removal of genes stratified by expression level. Effect of sequentially removing genes based on their expression levels (based on brain span expression^116^) on the cell type mutation bias correlation between ASD w/o ID and SCZ. Red line shows correlation changes when removing genes with lowest expression levels first, while blue dashed line shows removal of highest expressing genes first. Grey line and error bars show mean ± standard deviation for random gene removal.

**Figure S10.**
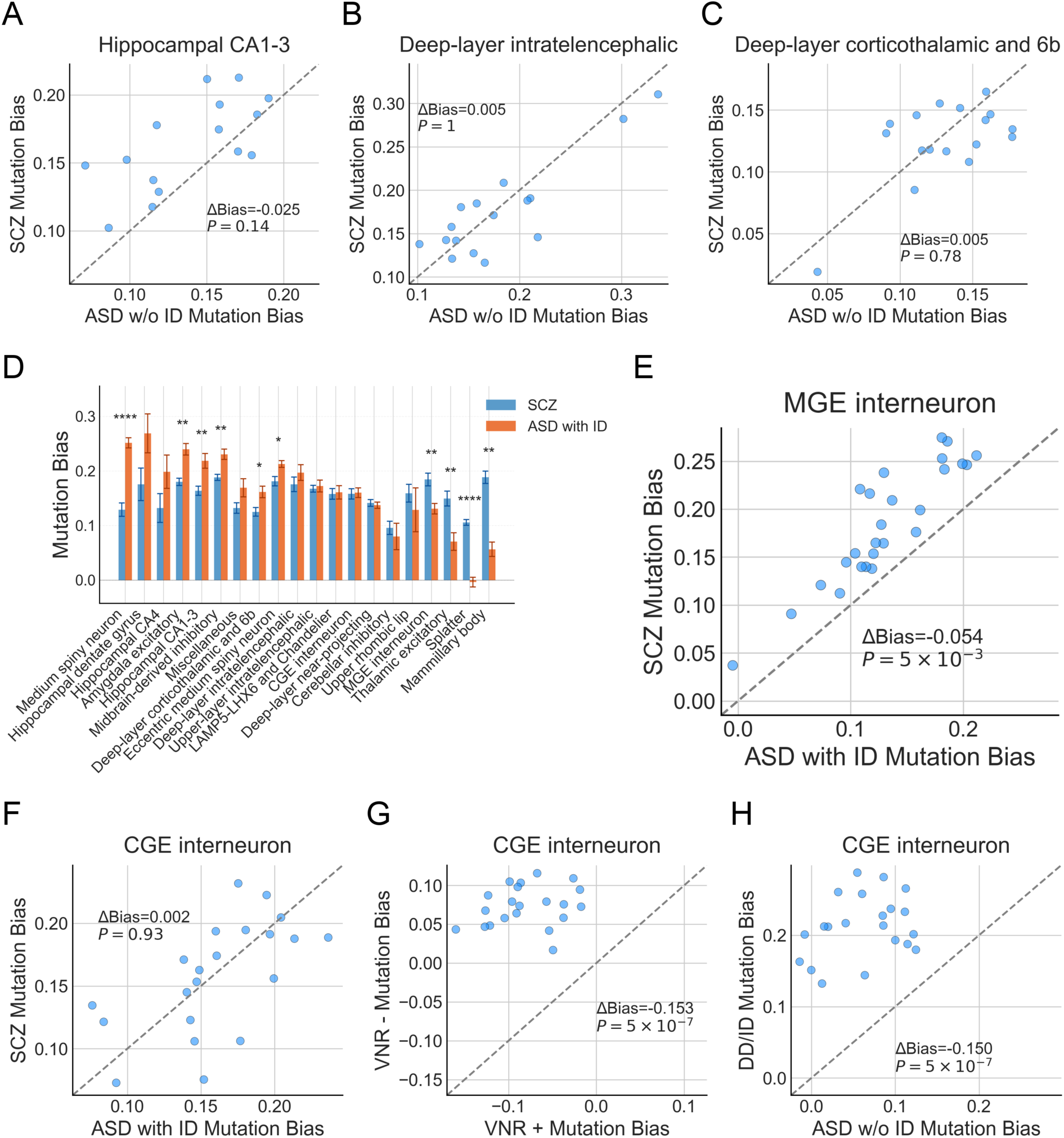
Supercluster-level mutation-bias contrasts across ASD, SCZ, DD/ID, and VNR gene sets. **A-C.** Detailed comparison of mutation biases in specific neuron superclusters between ASD w/o ID and SCZ, P-value was calculated based on Mann–Whitney U test with FDR correction: **A.** Hippocampal CA1-3 neurons. **B.** Deep-layer IT. **C.** Deep-layer corticothalamic and 6b. **D.** Bar plot comparing mutation biases across major neuron types between SCZ (blue) and ASD with ID (orange). Asterisks indicate significance level between groups (*: P < 0.05. **: P < 0.01. ***: P < 0.001, ****: P < 0.0001, Mann–Whitney U test with FDR correction). All neuronal cell types at supercluster level are displayed. Error bars represent standard error across clusters from corresponding supercluster. **E.** MGE interneurons show stronger bias in SCZ genes compared to ASD with ID genes **F.** CGE interneurons show similar bias in SCZ genes compared to ASD with ID genes. **G.** CGE interneurons show stronger bias in VNR-genes compared to VNR+ genes (verbal-numerical reasoning negative and positively associated genes). **H.** CGE interneurons show stronger bias in DD/ID genes compared to ASD w/o ID genes.

**Figure S11.**
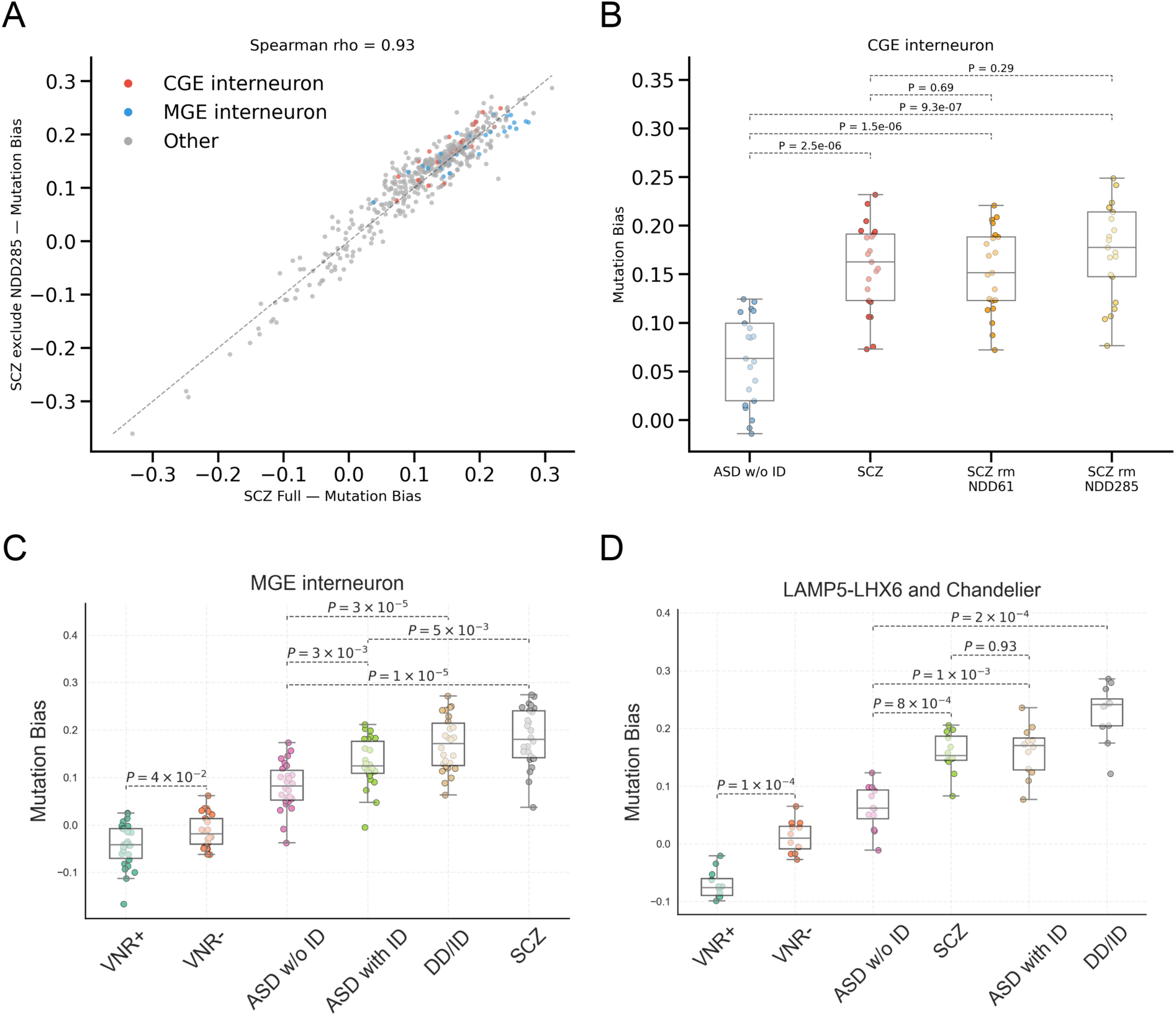
CGE bias in SCZ after removing DD-overlapping genes and MGE/LAMP5-LHX6 / Chandelier bias across disorders. **A.** Scatter plot of cell-type mutation bias (461 cell types) for the full SCZ gene set vs. SCZ after removing 9 DD285-overlapping genes. CGE interneurons highlighted in red, MGE interneurons in blue. **B.** CGE interneuron bias comparison across ASD without ID, SCZ (61 genes), SCZ rm DD61 (60 genes), and SCZ rm DD285 (52 genes). Mann– Whitney U test P-values are shown for each pairwise comparison; SCZ profiles before and after DD gene removal are not significantly different. **C**. Mutation bias distribution for MGE interneurons across different cohorts. **D**. Mutation bias distribution for LAMP5-LHX6 and Chandelier cells across different cohorts. P-values for pairwise comparison were shown, Mann–Whitney U test with FDR correction.

**Figure S12.**
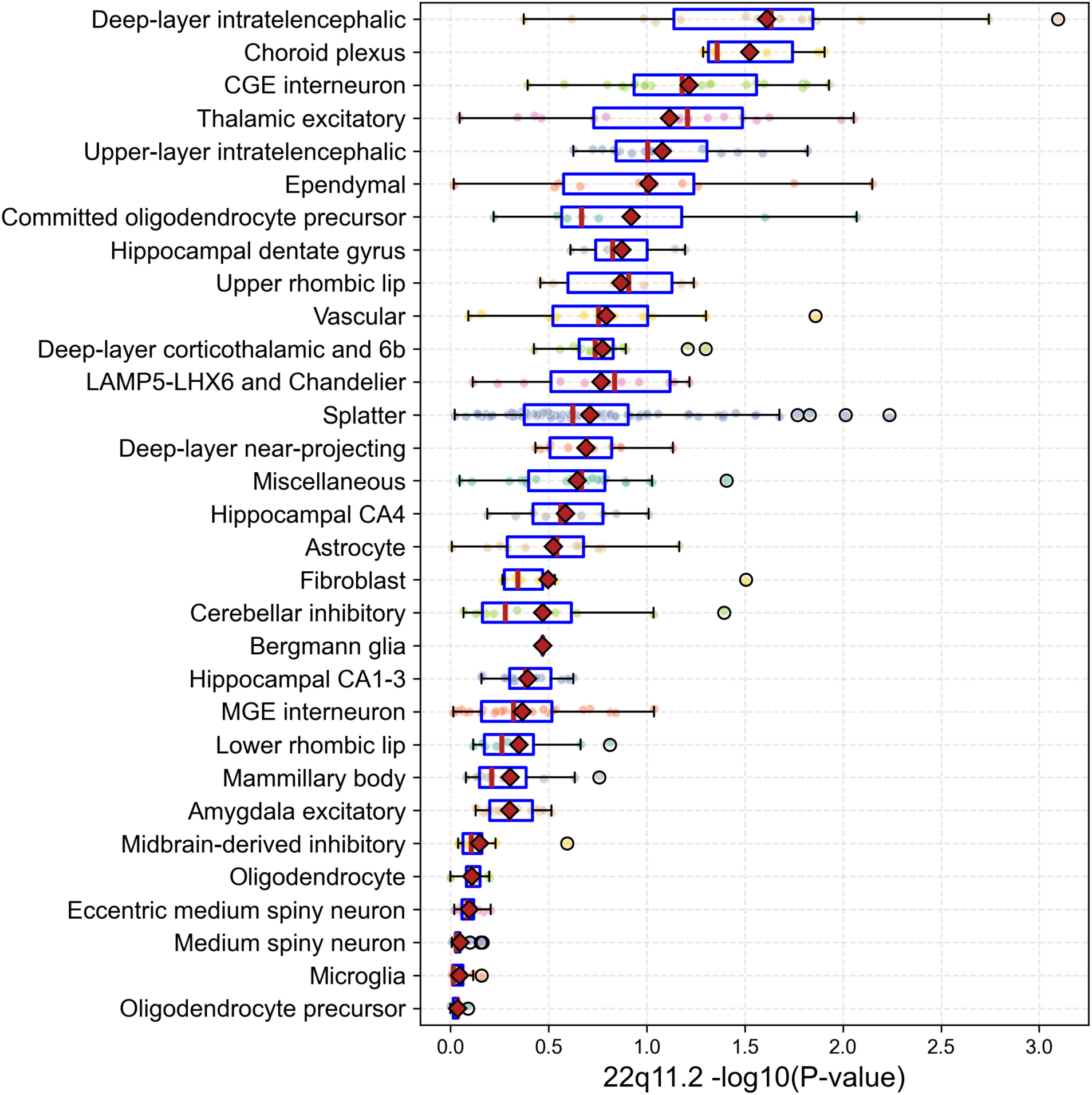
Supercluster mutation biases across brain cell types in 22q11.2 deletion. Box plots showing significance of mutation biases for all major brain cell types (superclusters). Each bar represents the bias distribution of the human adult brain neuron type belonging to a supercluster, X-axis shows the significance of biases as -log10(P-value). Red diamond shows average of each supercluster, red vertical line shows medium of each supercluster. No individual cell type cluster reached FDR-corrected significance (q < 0.1) for the 22q11.2 gene set.

**Figure S13.**
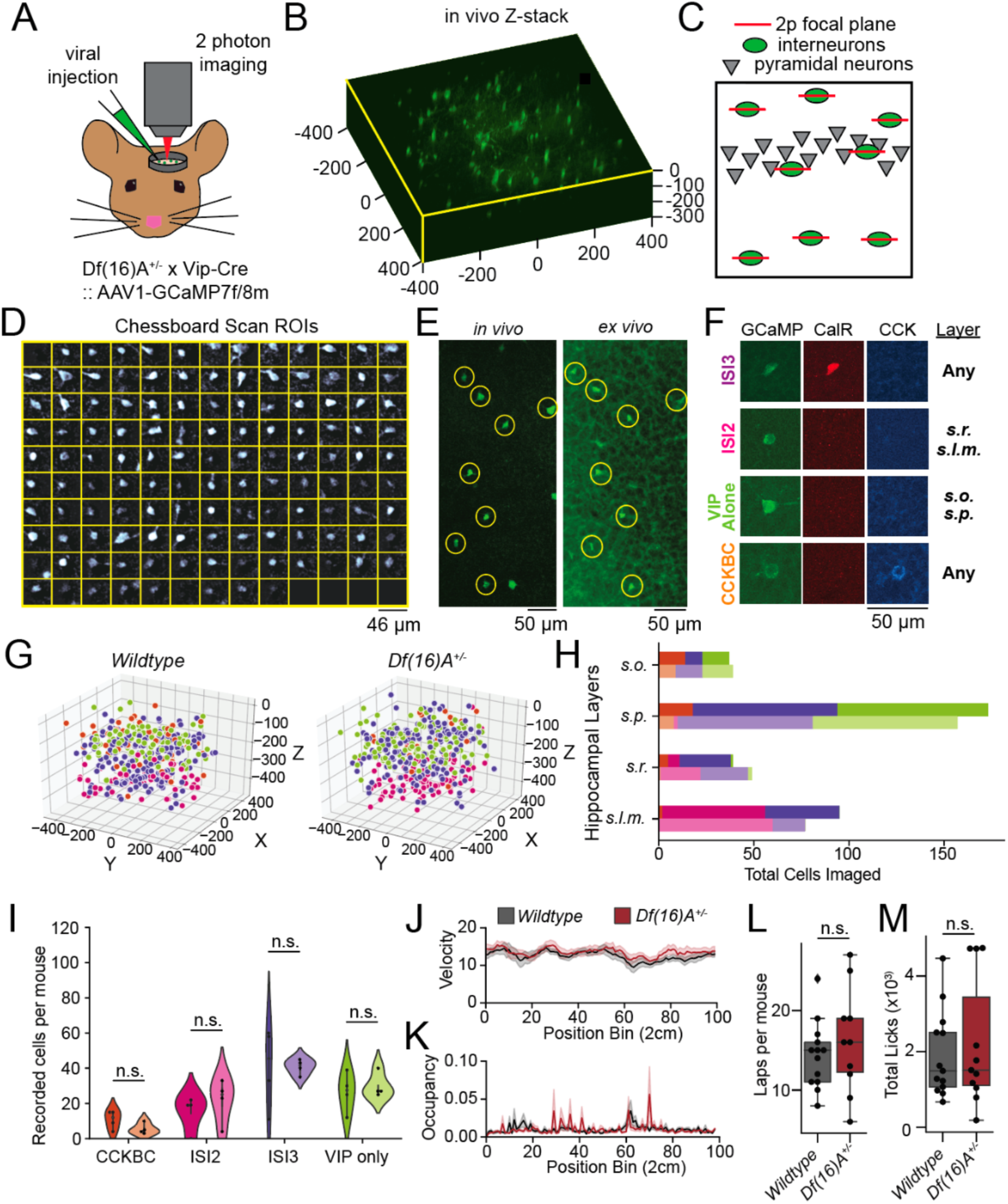
VIP+ Cell imaging and targeting strategy. **A.** Schematic for mouse crossing and viral injection and calcium imaging setup. **B.** 3D reconstruction of Z-stack from VIP-cre mice **C.** Schematic of 3D regions of interest (ROIs) imaged with AOD 2-photon microscopy. **D.** Example chessboard scan view of imaged ROIs. **E.** Manual registration of *in vivo* Z-stack section with *ex vivo* 75 µm transverse section for *post hoc* subtype classification with immunohistochemistry. **F.** Histological strategy for defining VIP-expressing cell subtypes. **G.** 3D distribution of all cells imaged from both genotypes. **H.** Distribution of VIP+ subtypes across CA1 hippocampal layers. **I.** Number of cells recorded per mouse (n=4, ISI3: ISI2: VIP+ Alone: CCKBC:). **J.** Average velocity in cm per second plotted by location on the belt of WT (black) and *Df(16)A^+/-^*(brown) mice. **K.** Average percent occupancy on the belt by position per genotype plotted by location on the belt of WT (black) and *Df(16)A^+/-^*(brown) mice. **L.** Total laps across mice within genotype (WT: 14.4 ± 1.1 laps, *Df(16)A^+/-^*: 16.2 ± 2.0 laps, Mann Whitney U = 51.5, P = 0.2). **M**. Licking proportion of WT (black) and *Df(16)A^+/-^*(brown). There is no significant difference in total licks across the belt during Random Foraging in WT (*n* = 4) or *Df(16)A^+/-^*(*n* = 4) depicted by box-and-whisker plot, WT: 1916 ± 304 licks, *Df(16)A^+/-^*: 2200 ± 489 licks, Mann-Whitney U Test, P = 0.4.

**Figure S14.**
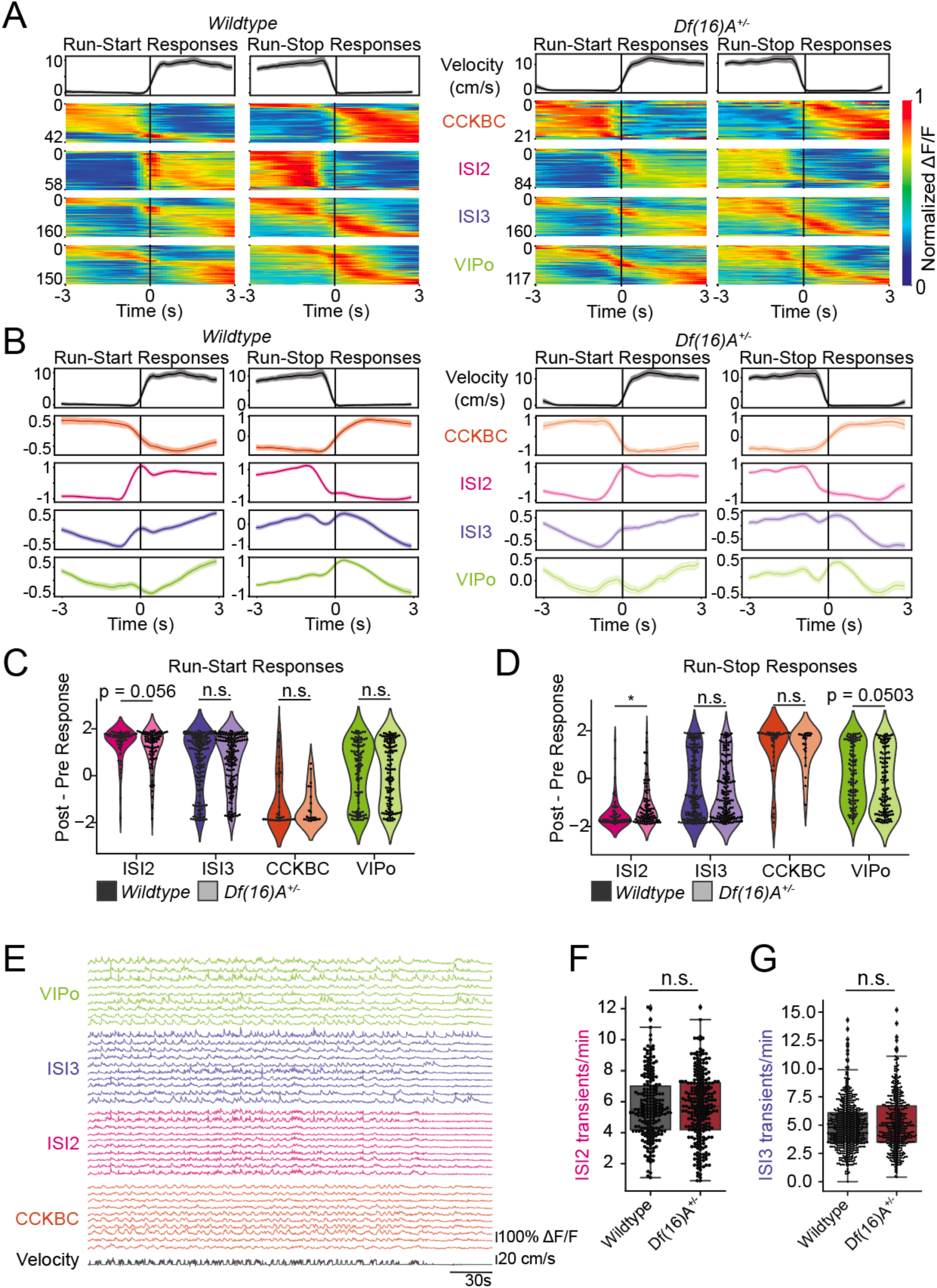
Subtype-specific changes during run-start and run-stop responses in VIP Interneurons. **A.** Individual, z-scored cell responses to run-start or run-stop behavior during random foraging task. Left: WT; Right: *Df(16)A^+/-^*. **B**. Averaged z-scored reward-onset peri-event time averages (PETAs) between -1.0 and 1.0 z-scores by cell type across all cells within cell type during the Random Foraging task for 3 seconds pre- and 3 seconds post-event. Left: WT; Right: *Df(16)A^+/-^*. **C**. Post - Pre PETA responses per cell type before and after Run-Start onset comparing the 3 seconds pre- to 3 seconds post-event. **D**. Post - Pre PETA responses per cell type before and after Run-Stop onset comparing the 3 seconds pre- to 3 seconds post-event. **E**. 10 example traces per cell type from a single session from mouse ID sah102, illustrating typical waveform distinction between subtypes. **F**. Number of transients from ISI2 cells per cell between genotypes (Mann-Whitney U test: P = 0.388; WT: 5.716 ± 0.142; *Df(16)A^+/-^*: 5.669 ± 0.137). **G**. Number of transients from ISI3 cells per cell between genotypes (Mann-Whitney U test: P = 0.093; WT: 4.989 ± 0.088; *Df(16)A^+/-^*: 5.211 ± 0.114). *CCKBC:* VIP+ and cholecystokinin + basket cell; *ISI3:* Interneuron-selective interneuron type 3; *ISI2:* Interneuron-selective interneuron type 2; *VIPo*: VIP+/Cal-/M2R-/CCK- cells in *stratum oriens* or *stratum pyramidale*. *: P < 0.05. n.s.: not significant.

**Figure S15.**
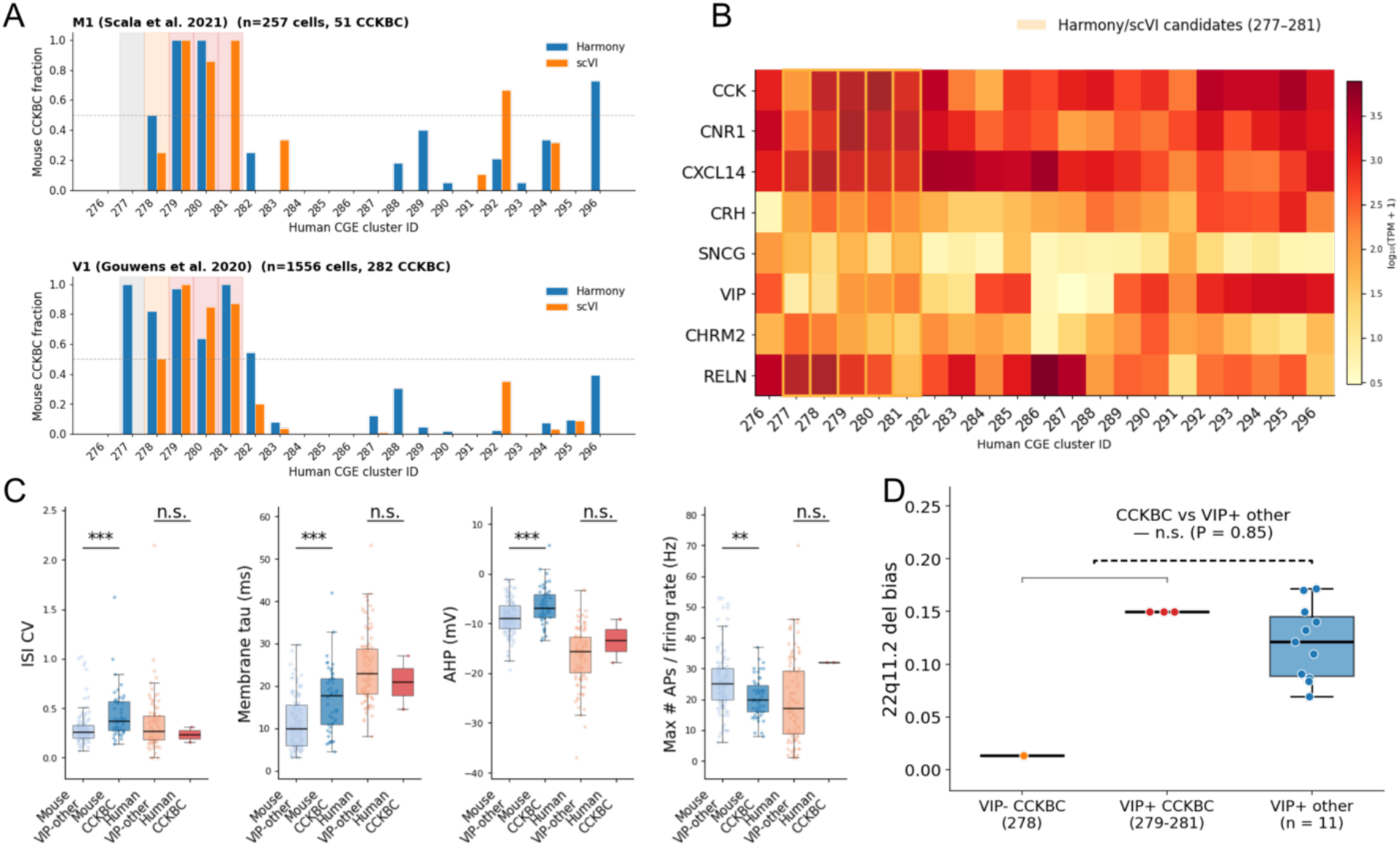
Cross-species identification of candidate human CCKBC-like clusters and 22q11.2 mutation bias within CGE interneurons. **A.** Cross-species transcriptomic mapping of mouse patch-seq CGE cells, including putative CCKBCs, onto the human Siletti et al. CGE supercluster (21 clusters, 276–296) using Harmony (blue) and scVI (orange). Per-cluster "mouse CCKBC fraction" = (assigned CCKBC cells) / (total assigned cells), shown for the M1 (Scala et al., 2021; top) and V1 (Gouwens et al., 2020; bottom) patch-seq datasets. Amber bands mark the five human CGE clusters (277–281) with majority CCKBC assignments in at least one method/dataset combination. **B.** Pseudobulk expression of canonical CCKBC markers and discriminating markers across the 21 human Siletti CGE clusters (log₁₀(TPM + 1) color scale). The five Harmony/scVI candidate clusters are marked by amber outlines. **C.** Cross-species electrophysiology for mouse VIP-other (n = 112), mouse CCKBC (n = 50), human VIP-other (n = 100), and human CCKBC (n = 2) on four features that discriminate mouse CCKBCs from mouse VIP-other interneurons. Brackets indicate within-species Mann–Whitney U two-sided P-values (*** P < 0.001, ** P < 0.01, n.s. = not significant). **D.** 22q11.2 deletion mutation bias for the four putative CCKBC clusters and the 11 VIP+ other CGE clusters. All-CCKBC vs VIP+ other: Mann– Whitney U, two-sided P = 0.85 (n.s., dashed bracket).

## References

1. Singh, T., Poterba, T., Curtis, D., Akil, H., Al Eissa, M., Barchas, J.D., Bass, N., Bigdeli, T.B., Breen, G., Bromet, E.J., et al. (2022). Rare coding variants in ten genes confer substantial risk for schizophrenia. Nature 604, 509–516.

2. Zhou, X., Feliciano, P., Shu, C., Wang, T., Astrovskaya, I., Hall, J.B., Obiajulu, J.U., Wright, J.R., Murali, S.C., Xu, S.X., et al. (2022). Integrating de novo and inherited variants in 42,607 autism cases identifies mutations in new moderate-risk genes. Nat. Genet. 54, 1305–1319.

3. Gilman, S.R., Chang, J., Xu, B., Bawa, T.S., Gogos, J.A., Karayiorgou, M., and Vitkup, D. (2012). Diverse types of genetic variation converge on functional gene networks involved in schizophrenia. Nat. Neurosci. 15, 1723–1728.

4. Gilman, S.R., Iossifov, I., Levy, D., Ronemus, M., Wigler, M., and Vitkup, D. (2011). Rare de novo variants associated with autism implicate a large functional network of genes involved in formation and function of synapses. Neuron 70, 898–907.

5. Gandal, M.J., Haney, J.R., Parikshak, N.N., Leppa, V., Ramaswami, G., Hartl, C., Schork, A.J., Appadurai, V., Buil, A., Werge, T.M., et al. (2018). Shared molecular neuropathology across major psychiatric disorders parallels polygenic overlap. Science 359, 693–697.

6. Rees, E., Creeth, H.D.J., Hwu, H.-G., Chen, W.J., Tsuang, M., Glatt, S.J., Rey, R., Kirov, G., Walters, J.T.R., Holmans, P., et al. (2021). Schizophrenia, autism spectrum disorders and developmental disorders share specific disruptive coding mutations. Nat. Commun. 12, 5353.

7. Kaladjian, A., Jeanningros, R., Azorin, J.-M., Anton, J.-L., and Mazzola-Pomietto, P. (2011). Impulsivity and neural correlates of response inhibition in schizophrenia. Psychol. Med. 41, 291–299.

8. Trevisan, D.A., Foss-Feig, J.H., Naples, A.J., Srihari, V., Anticevic, A., and McPartland, J.C. (2020). Autism spectrum disorder and schizophrenia are better differentiated by positive symptoms than negative symptoms. Front. Psychiatry 11, 548.

9. Andreasen, N.C., and Olsen, S. (1982). Negative v positive schizophrenia. Definition and validation. Arch. Gen. Psychiatry 39, 789–794.

10. Forbes, N.F., Carrick, L.A., McIntosh, A.M., and Lawrie, S.M. (2009). Working memory in schizophrenia: a meta-analysis. Psychol. Med. 39, 889–905.

11. Green, M., and Walker, E. (1985). Neuropsychological performance and positive and negative symptoms in schizophrenia. J. Abnorm. Psychol. 94, 460–469.

12. Spek, A.A., and Wouters, S.G.M. (2010). Autism and schizophrenia in high functioning adults: Behavioral differences and overlap. Res. Autism Spectr. Disord. 4, 709–717.

13. Schneider, M., Debbané, M., Bassett, A.S., Chow, E.W.C., Fung, W.L.A., van den Bree, M., Owen, M., Murphy, K.C., Niarchou, M., Kates, W.R., et al. (2014). Psychiatric disorders from childhood to adulthood in 22q11.2 deletion syndrome: results from the International Consortium on Brain and Behavior in 22q11.2 Deletion Syndrome. Am. J. Psychiatry 171, 627–639.

14. Karayiorgou, M., Simon, T.J., and Gogos, J.A. (2010). 22q11.2 microdeletions: linking DNA structural variation to brain dysfunction and schizophrenia. Nature Reviews Neuroscience *2010 11:6 11*, 402–416.

15. Rees, E., Kendall, K., Pardiñas, A.F., Legge, S.E., Pocklington, A., Escott-Price, V., MacCabe, J.H., Collier, D.A., Holmans, P., O’Donovan, M.C., et al. (2016). Analysis of intellectual disability copy number variants for association with schizophrenia. JAMA Psychiatry 73, 963–969.

16. Miller, D.T., Adam, M.P., Aradhya, S., Biesecker, L.G., Brothman, A.R., Carter, N.P., Church, D.M., Crolla, J.A., Eichler, E.E., Epstein, C.J., et al. (2010). Consensus statement: chromosomal microarray is a first-tier clinical diagnostic test for individuals with developmental disabilities or congenital anomalies. Am. J. Hum. Genet. 86, 749–764.

17. Ropers, H.H. (2008). Genetics of intellectual disability. Curr. Opin. Genet. Dev. 18, 241–250.

18. Vissers, L.E.L.M., Gilissen, C., and Veltman, J.A. (2016). Genetic studies in intellectual disability and related disorders. Nat. Rev. Genet. 17, 9–18.

19. Millan, M.J., Agid, Y., Brüne, M., Bullmore, E.T., Carter, C.S., Clayton, N.S., Connor, R., Davis, S., Deakin, B., DeRubeis, R.J., et al. (2012). Cognitive dysfunction in psychiatric disorders: characteristics, causes and the quest for improved therapy. Nat. Rev. Drug Discov. 11, 141–168.

20. Harvey, P.D., and Strassnig, M. (2012). Predicting the severity of everyday functional disability in people with schizophrenia: cognitive deficits, functional capacity, symptoms, and health status. World Psychiatry 11, 73–79.

21. Bowie, C.R., and Harvey, P.D. (2006). Cognitive deficits and functional outcome in schizophrenia. Neuropsychiatr. Dis. Treat. 10.2147/nedt.s24531.

22. Bowie, C.R., Reichenberg, A., Patterson, T.L., Heaton, R.K., and Harvey, P.D. (2006). Determinants of real-world functional performance in schizophrenia subjects: Correlations with cognition, functional capacity, and symptoms. Am. J. Psychiatry 163, 418–425.

23. Trubetskoy, V., Pardiñas, A.F., Qi, T., Panagiotaropoulou, G., Awasthi, S., Bigdeli, T.B., Bryois, J., Chen, C.Y., Dennison, C.A., Hall, L.S., et al. (2022). Mapping genomic loci implicates genes and synaptic biology in schizophrenia. Nature 604, 502–508.

24. Bryois, J., Skene, N.G., Hansen, T.F., Kogelman, L.J.A., Watson, H.J., Liu, Z., Eating Disorders Working Group of the Psychiatric Genomics Consortium, International Headache Genetics Consortium, 23andMe Research Team, Brueggeman, L., et al. (2020). Genetic identification of cell types underlying brain complex traits yields insights into the etiology of Parkinson’s disease. Nat. Genet. *52*, 482–493.

25. Duncan, L.E., Li, T., Salem, M., Li, W., Mortazavi, L., Senturk, H., Shahverdizadeh, N., Vesuna, S., Shen, H., Yoon, J., et al. (2025). Mapping the cellular etiology of schizophrenia and complex brain phenotypes. Nat. Neurosci., 1–11.

26. Yao, S., Harder, A., Darki, F., Chang, Y.-W., Li, A., Nikouei, K., Volpe, G., Lundström, J.N., Zeng, J., Wray, N., et al. (2025). Connecting genomic results for psychiatric disorders to human brain cell types and regions reveals convergence with functional connectivity. Nature Communications. 10.1101/2024.01.18.24301478.

27. Broglio, C., Martín-Monzón, I., Ocaña, F.M., Gómez, A., Durán, E., Salas, C., and Rodríguez, F. (2015). Hippocampal pallium and map-like memories through vertebrate evolution. J. Behav. Brain Sci. 05, 109–120.

28. Rodríguez, F., López, J.C., Vargas, J.P., Broglio, C., Gómez, Y., and Salas, C. (2002). Spatial memory and hippocampal pallium through vertebrate evolution: insights from reptiles and teleost fish. Brain Res. Bull. 57, 499–503.

29. Buzsáki, G. (2002). Theta oscillations in the hippocampus. Neuron 33, 325–340.

30. Pelkey, K.A., Vargish, G.A., Pellegrini, L.V., Calvigioni, D., Chapeton, J., Yuan, X., Hunt, S., Cummins, A.C., Eldridge, M.A.G., Pickel, J., et al. (2023). Evolutionary conservation of hippocampal mossy fiber synapse properties. Neuron 111, 3802–3818.e5.

31. Hamm, J.P., Peterka, D.S., Gogos, J.A., and Yuste, R. (2017). Altered Cortical Ensembles in Mouse Models of Schizophrenia. Neuron 94, 153–167.e8.

32. Herrlinger, S.A., Rao, B.Y., Conde Paredes, M.E., Tuttman, A.L., Arain, H., Varol, E., Gogos, J.A., and Losonczy, A. (2024). Disorganized Inhibitory Dynamics and Functional Connectivity in Hippocampal area CA1 of 22q11.2 Deletion Mutant Mice. bioRxiv, 2024.04.28.591464. 10.1101/2024.04.28.591464.

33. Gogos, J.A., Crabtree, G., and Diamantopoulou, A. (2020). The abiding relevance of mouse models of rare mutations to psychiatric neuroscience and therapeutics. Schizophr. Res. 217, 37–51.

34. Diamantopoulou, A., and Gogos, J.A. (2019). Neurocognitive and Perceptual Processing in Genetic Mouse Models of Schizophrenia: Emerging Lessons. Neuroscientist 25, 597–619.

35. Arguello, P.A., and Gogos, J.A. (2006). Modeling madness in mice: one piece at a time. Neuron 52, 179–196.

36. Gordon, A., Forsingdal, A., Klewe, I.V., Nielsen, J., Didriksen, M., Werge, T., and Geschwind, D.H. (2019). Transcriptomic networks implicate neuronal energetic abnormalities in three mouse models harboring autism and schizophrenia-associated mutations. Molecular Psychiatry 2019 26:5 26, 1520–1534.

37. Siletti, K., Hodge, R., Mossi Albiach, A., Lee, K.W., Ding, S.-L., Hu, L., Lönnerberg, P., Bakken, T., Casper, T., Clark, M., et al. (2023). Transcriptomic diversity of cell types across the adult human brain. Science 382, eadd7046.

38. Geschwind, D.H., and Flint, J. (2015). Genetics and genomics of psychiatric disease. Science 349, 1489–1494.

39. Iossifov, I., O’Roak, B.J., Sanders, S.J., Ronemus, M., Krumm, N., Levy, D., Stessman, H.A., Witherspoon, K.T., Vives, L., Patterson, K.E., et al. (2014). The contribution of de novo coding mutations to autism spectrum disorder. Nature 515, 216–221.

40. Matson, J.L., and Shoemaker, M. (2009). Intellectual disability and its relationship to autism spectrum disorders. Res. Dev. Disabil. 30, 1107–1114.

41. Woodberry, K.A., Giuliano, A.J., and Seidman, L.J. (2008). Premorbid IQ in schizophrenia: a meta-analytic review. Am. J. Psychiatry 165, 579–587.

42. Khandaker, G.M., Barnett, J.H., White, I.R., and Jones, P.B. (2011). A quantitative meta-analysis of population-based studies of premorbid intelligence and schizophrenia. Schizophr. Res. 132, 220–227.

43. Creeth, H.D.J., Rees, E., Legge, S.E., Dennison, C.A., Holmans, P., Walters, J.T.R., O’Donovan, M.C., and Owen, M.J. (2022). Ultrarare coding variants and cognitive function in schizophrenia. JAMA Psychiatry 79, 963–970.

44. Nuechterlein, K.H., Green, M.F., Kern, R.S., Baade, L.E., Barch, D.M., Cohen, J.D., Essock, S., Fenton, W.S., Frese, F.J., 3rd, Gold, J.M., et al. (2008). The MATRICS Consensus Cognitive Battery, part 1: test selection, reliability, and validity. Am. J. Psychiatry *165*, 203– 213.

45. Mesholam-Gately, R.I., Giuliano, A.J., Goff, K.P., Faraone, S.V., and Seidman, L.J. (2009). Neurocognition in first-episode schizophrenia: a meta-analytic review. Neuropsychology 23, 315–336.

46. Minshew, N.J., Goldstein, G., and Siegel, D.J. (1997). Neuropsychologic functioning in autism: profile of a complex information processing disorder. J. Int. Neuropsychol. Soc. 3, 303–316.

47. Hill, E.L. (2004). Executive dysfunction in autism. Trends Cogn. Sci. 8, 26–32.

48. Jutla, A., Foss-Feig, J., and Veenstra-VanderWeele, J. (2022). Autism spectrum disorder and schizophrenia: An updated conceptual review. Autism Res. 15, 384–412.

49. Palmer, D.S., Howrigan, D.P., Chapman, S.B., Adolfsson, R., Bass, N., Blackwood, D., Boks, M.P.M., Chen, C.-Y., Churchhouse, C., Corvin, A.P., et al. (2022). Exome sequencing in bipolar disorder identifies AKAP11 as a risk gene shared with schizophrenia. Nat. Genet. 54, 541–547.

50. Kaplanis, J., Samocha, K.E., Wiel, L., Zhang, Z., Arvai, K.J., Eberhardt, R.Y., Gallone, G., Lelieveld, S.H., Martin, H.C., McRae, J.F., et al. (2020). Evidence for 28 genetic disorders discovered by combining healthcare and research data. Nature 586, 757–762.

51. Aleman, A., and Kahn, R. (2005). Strange feelings: Do amygdala abnormalities dysregulate the emotional brain in schizophrenia? Prog. Neurobiol. 77, 283–298.

52. Baron-Cohen, S., Ring, H.A., Bullmore, E.T., Wheelwright, S., Ashwin, C., and Williams, S.C. (2000). The amygdala theory of autism. Neurosci. Biobehav. Rev. 24, 355–364.

53. Banker, S.M., Gu, X., Schiller, D., and Foss-Feig, J.H. (2021). Hippocampal contributions to social and cognitive deficits in autism spectrum disorder. Trends Neurosci. 44, 793–807.

54. Lieberman, J.A., Girgis, R.R., Brucato, G., Moore, H., Provenzano, F., Kegeles, L., Javitt, D., Kantrowitz, J., Wall, M.M., Corcoran, C.M., et al. (2018). Hippocampal dysfunction in the pathophysiology of schizophrenia: a selective review and hypothesis for early detection and intervention. Mol. Psychiatry 23, 1764–1772.

55. Zierhut, K., Bogerts, B., Schott, B., Fenker, D., Walter, M., Albrecht, D., Steiner, J., Schütze, H., Northoff, G., Düzel, E., et al. (2010). The role of hippocampus dysfunction in deficient memory encoding and positive symptoms in schizophrenia. Psychiatry Res. 183, 187–194.

56. Moberg, S., and Takahashi, N. (2022). Neocortical layer 5 subclasses: From cellular properties to roles in behavior. Front. Synaptic Neurosci. 14, 1006773.

57. Chang, J., Gilman, S.R., Chiang, A.H., Sanders, S.J., and Vitkup, D. (2015). Genotype to phenotype relationships in autism spectrum disorders. Nat. Neurosci. 18, 191–198.

58. Peça, J., Feliciano, C., Ting, J.T., Wang, W., Wells, M.F., Venkatraman, T.N., Lascola, C.D., Fu, Z., and Feng, G. (2011). Shank3 mutant mice display autistic-like behaviours and striatal dysfunction. Nature 472, 437–442.

59. Rothwell, P.E., Fuccillo, M.V., Maxeiner, S., Hayton, S.J., Gokce, O., Lim, B.K., Fowler, S.C., Malenka, R.C., and Südhof, T.C. (2014). Autism-associated neuroligin-3 mutations commonly impair striatal circuits to boost repetitive behaviors. Cell 158, 198–212.

60. Van Derveer, A.B., Bastos, G., Ferrell, A.D., Gallimore, C.G., Greene, M.L., Holmes, J.T., Kubricka, V., Ross, J.M., and Hamm, J.P. (2021). A role for somatostatin-positive interneurons in neuro-oscillatory and information processing deficits in schizophrenia. Schizophr. Bull. 47, 1385–1398.

61. Batiuk, M.Y., Tyler, T., Dragicevic, K., Mei, S., Rydbirk, R., Petukhov, V., Deviatiiarov, R., Sedmak, D., Frank, E., Feher, V., et al. (2022). Upper cortical layer-driven network impairment in schizophrenia. Sci. Adv. 8, eabn8367.

62. Satterstrom, F.K., Kosmicki, J.A., Wang, J., Breen, M.S., De Rubeis, S., An, J.-Y., Peng, M., Collins, R., Grove, J., Klei, L., et al. (2020). Large-Scale Exome Sequencing Study Implicates Both Developmental and Functional Changes in the Neurobiology of Autism. Cell 180, 568–584.e23.

63. Karczewski, K.J., Francioli, L.C., Tiao, G., Cummings, B.B., Alföldi, J., Wang, Q., Collins, R.L., Laricchia, K.M., Ganna, A., Birnbaum, D.P., et al. (2020). The mutational constraint spectrum quantified from variation in 141,456 humans. Nature 581, 434–443.

64. Chen, C.-Y., Tian, R., Ge, T., Lam, M., Sanchez-Andrade, G., Singh, T., Urpa, L., Liu, J.Z., Sanderson, M., Rowley, C., et al. (2023). The impact of rare protein coding genetic variation on adult cognitive function. Nat. Genet. 55, 927–938.

65. Fawns-Ritchie, C., and Deary, I.J. (2020). Reliability and validity of the UK Biobank cognitive tests. PLoS One 15, e0231627.

66. Marshall, C.R., Howrigan, D.P., Merico, D., Thiruvahindrapuram, B., Wu, W., Greer, D.S., Antaki, D., Shetty, A., Holmans, P.A., Pinto, D., et al. (2017). Contribution of copy number variants to schizophrenia from a genome-wide study of 41,321 subjects. Nat. Genet. 49, 27– 35.

67. Matsunami, N., Hadley, D., Hensel, C.H., Christensen, G.B., Kim, C., Frackelton, E., Thomas, K., da Silva, R.P., Stevens, J., Baird, L., et al. (2013). Identification of rare recurrent copy number variants in high-risk autism families and their prevalence in a large ASD population. PLoS One 8, e52239.

68. Robin, N.H., and Shprintzen, R.J. (2005). Defining the clinical spectrum of deletion 22q11.2. J. Pediatr. 147, 90–96.

69. Tang, K.L., Antshel, K.M., Fremont, W.P., and Kates, W.R. (2015). Behavioral and psychiatric phenotypes in 22q11.2 deletion syndrome. J. Dev. Behav. Pediatr. 36, 639–650.

70. Swillen, A., Devriendt, K., Legius, E., Eyskens, B., Dumoulin, M., Gewillig, M., and Fryns, J.P. (1997). Intelligence and psychosocial adjustment in velocardiofacial syndrome: a study of 37 children and adolescents with VCFS. J. Med. Genet. 34, 453–458.

71. Kepecs, A., and Fishell, G. (2014). Interneuron cell types are fit to function. Nature 505, 318– 326.

72. Chamberland, S., and Topolnik, L. (2012). Inhibitory control of hippocampal inhibitory neurons. Front. Neurosci. 0, 165.

73. Letzkus, J.J., Wolff, S.B.E., and Lüthi, A. (2015). Disinhibition, a circuit mechanism for associative learning and memory. Neuron 88, 264–276.

74. Pfeffer, C.K., Xue, M., He, M., Huang, Z.J., and Scanziani, M. (2013). Inhibition of inhibition in visual cortex: the logic of connections between molecularly distinct interneurons. Nat. Neurosci. 16, 1068–1076.

75. Fu, Y., Tucciarone, J.M., Espinosa, J.S., Sheng, N., Darcy, D.P., Nicoll, R.A., and Stryker M (2014). A cortical circuit for gain control by behavioral state. Cell 156, 1139–1152.

76. Bennett, C., Arroyo, S., Berns, D., and Hestrin, S. (2012). Mechanisms generating dual-component nicotinic EPSCs in cortical interneurons. J. Neurosci. 32, 17287–17296.

77. Letzkus, J.J., Wolff, S.B.E., Meyer, E.M.M., Tovote, P., Courtin, J., Herry, C., and Lüthi, A. (2011). A disinhibitory microcircuit for associative fear learning in the auditory cortex. Nature 480, 331–335.

78. Poorthuis, R.B., Enke, L., and Letzkus, J.J. (2014). Cholinergic circuit modulation through differential recruitment of neocortical interneuron types during behaviour: Cholinergic circuit modulation via interneurons. J. Physiol. 592, 4155–4164.

79. Froemke, R.C., Merzenich, M.M., and Schreiner, C.E. (2007). A synaptic memory trace for cortical receptive field plasticity. Nature 450, 425–429.

80. Lee, A.T., Cunniff, M.M., See, J.Z., Wilke, S.A., Luongo, F.J., Ellwood, I.T., Ponnavolu, S., and Sohal, V.S. (2019). VIP interneurons contribute to avoidance behavior by regulating information flow across hippocampal-prefrontal networks. Neuron 102, 1223–1234.e4.

81. Stark, K.L., Xu, B., Bagchi, A., Lai, W.S., Liu, H., Hsu, R., Wan, X., Pavlidis, P., Mills, A.A., Karayiorgou, M., et al. (2008). Altered brain microRNA biogenesis contributes to phenotypic deficits in a 22q11-deletion mouse model. Nat. Genet. 40, 751–760.

82. Drew, L.J., Stark, K.L., Fénelon, K., Karayiorgou, M., Macdermott, A.B., and Gogos, J.A. (2011). Evidence for altered hippocampal function in a mouse model of the human 22q11.2 microdeletion. Mol. Cell. Neurosci. 47, 293–305.

83. Zaremba, J.D., Diamantopoulou, A., Danielson, N.B., Grosmark, A.D., Kaifosh, P.W., Bowler, J.C., Liao, Z., Sparks, F.T., Gogos, J.A., and Losonczy, A. (2017). Impaired hippocampal place cell dynamics in a mouse model of the 22q11.2 deletion. Nat. Neurosci. 20, 1612– 1623.

84. Sun, Z., Williams, D.J., Xu, B., and Gogos, J.A. (2018). Altered function and maturation of primary cortical neurons from a 22q11.2 deletion mouse model of schizophrenia. Transl. Psychiatry 8. 10.1038/s41398-018-0132-8.

85. Rodriguez, E., Sakurai, K., Xu, J., Chen, Y., Toda, K., Zhao, S., Han, B.-X., Ryu, D., Yin, H., Liedtke, W., et al. (2017). A craniofacial-specific monosynaptic circuit enables heightened affective pain. Nat. Neurosci. 20, 1734–1743.

86. Geiller, T., Vancura, B., Terada, S., Troullinou, E., Chavlis, S., Tsagkatakis, G., Tsakalides, P., Ócsai, K., Poirazi, P., Rózsa, B.J., et al. (2020). Large-Scale 3D Two-Photon Imaging of Molecularly Identified CA1 Interneuron Dynamics in Behaving Mice. Neuron 108, 968–983.e9.

87. Szalay, G., Judák, L., Katona, G., Ócsai, K., Juhász, G., Veress, M., Szadai, Z., Fehér, A., Tompa, T., Chiovini, B., et al. (2016). Fast 3D imaging of spine, dendritic, and neuronal assemblies in behaving animals. Neuron 92, 723–738.

88. Katona, G., Szalay, G., Maák, P., Kaszás, A., Veress, M., Hillier, D., Chiovini, B., Vizi, E.S., Roska, B., and Rózsa, B. (2012). Fast two-photon in vivo imaging with three-dimensional random-access scanning in large tissue volumes. Nat. Methods 9, 201–208.

89. Vancura, B., Geiller, T., Grosmark, A., Zhao, V., and Losonczy, A. (2023). Inhibitory control of sharp-wave ripple duration during learning in hippocampal recurrent networks. Nat. Neurosci. 26, 788–797.

90. Somogyi, P., and Klausberger, T. (2005). Defined types of cortical interneurone structure space and spike timing in the hippocampus. J. Physiol. 562, 9–26.

91. Guet-McCreight, A., Skinner, F.K., and Topolnik, L. (2020). Common principles in functional organization of VIP/calretinin cell-driven disinhibitory circuits across cortical areas. Front. Neural Circuits 14, 32.

92. Varga, C., Golshani, P., and Soltesz, I. (2012). Frequency-invariant temporal ordering of interneuronal discharges during hippocampal oscillations in awake mice. Proc. Natl. Acad. Sci. U. S. A. 109, E2726–34.

93. Lapray, D., Lasztoczi, B., Lagler, M., Viney, T.J., Katona, L., Valenti, O., Hartwich, K., Borhegyi, Z., Somogyi, P., and Klausberger, T. (2012). Behavior-dependent specialization of identified hippocampal interneurons. Nat. Neurosci. 15, 1265–1271.

94. Katona, L., Lapray, D., Viney, T.J., Oulhaj, A., Borhegyi, Z., Micklem, B.R., Klausberger, T., and Somogyi, P. (2014). Sleep and movement differentiates actions of two types of somatostatin-expressing GABAergic interneuron in rat hippocampus. Neuron 82, 872–886.

95. Somogyi, P., Katona, L., Klausberger, T., Lasztóczi, B., and Viney, T.J. (2014). Temporal redistribution of inhibition over neuronal subcellular domains underlies state-dependent rhythmic change of excitability in the hippocampus. Philos. Trans. R. Soc. Lond. B Biol. Sci. 369, 20120518.

96. Turi, G.F., Li, W.-K., Chavlis, S., Pandi, I., Hare, J.O., Priestley, J.B., Grosmark, A.D., Liao, Z., Ladow, M., Zhang, J.F., et al. (2019). Vasoactive Intestinal Polypeptide-Expressing Interneurons in the Hippocampus Support Goal-Oriented Spatial Learning. Neuron, 1–16.

97. Dudok, B., Klein, P.M., Hwaun, E., Lee, B.R., Yao, Z., Fong, O., Bowler, J.C., Terada, S., Sparks, F.T., Szabo, G.G., et al. (2021). Alternating sources of perisomatic inhibition during behavior. Neuron 109, 997–1012.e9.

98. Friedrich, J., Zhou, P., and Paninski, L. (2017). Fast online deconvolution of calcium imaging data. PLoS Comput. Biol. 13, e1005423.

99. Buzsáki, G., and Moser, E.I. (2013). Memory, navigation and theta rhythm in the hippocampal-entorhinal system. Nature Neuroscience 2013 16:2 16, 130–138.

100. O’Keefe, J., and Dostrovsky, J. (1971). The hippocampus as a spatial map. Preliminary evidence from unit activity in the freely-moving rat. Brain Res. 34, 171–175.

101. Robinson, E.B., Samocha, K.E., Kosmicki, J.A., McGrath, L., Neale, B.M., Perlis, R.H., and Daly, M.J. (2014). Autism spectrum disorder severity reflects the average contribution of de novo and familial influences. Proc. Natl. Acad. Sci. U. S. A. 111, 15161–15165.

102. Gonzalez-Burgos, G., and Lewis, D.A. (2012). NMDA receptor hypofunction, parvalbumin-positive neurons, and cortical gamma oscillations in schizophrenia. Schizophr. Bull. 38, 950– 957.

103. Buzsáki, G., and Draguhn, A. (2004). Neuronal oscillations in cortical networks. Science 304, 1926–1929.

104. Möhler, H., and Rudolph, U. (2017). Disinhibition, an emerging pharmacology of learning and memory. F1000Res. *6*, 101.

105. Lovett-Barron, M., and Losonczy, A. (2014). Behavioral consequences of GABAergic neuronal diversity. Curr. Opin. Neurobiol. 26, 27–33.

106. Luo, X., Guet-McCreight, A., Villette, V., Francavilla, R., Marino, B., Chamberland, S., Skinner, F.K., and Topolnik, L. (2020). Synaptic mechanisms underlying the network state-dependent recruitment of VIP-expressing interneurons in the CA1 hippocampus. Cereb. Cortex 30, 3667–3685.

107. Kamigaki, T., and Dan, Y. (2017). Delay activity of specific prefrontal interneuron subtypes modulates memory-guided behavior. Nat. Neurosci. 20, 854–863.

108. Goff, K.M., and Goldberg, E.M. (2021). A role for vasoactive intestinal peptide interneurons in neurodevelopmental disorders. Dev. Neurosci. 43, 168–180.

109. Mossner, J.M., Batista-Brito, R., Pant, R., and Cardin, J.A. (2020). Developmental loss of MeCP2 from VIP interneurons impairs cortical function and behavior. Elife 9. 10.7554/eLife.55639.

110. Goff, K.M., Liebergall, S.R., Jiang, E., Somarowthu, A., and Goldberg, E.M. (2023). VIP interneuron impairment promotes in vivo circuit dysfunction and autism-related behaviors in Dravet syndrome. Cell Rep. 42, 112628.

111. Murray, J.D., Anticevic, A., Gancsos, M., Ichinose, M., Corlett, P.R., Krystal, J.H., and Wang, X.-J. (2014). Linking microcircuit dysfunction to cognitive impairment: effects of disinhibition associated with schizophrenia in a cortical working memory model. Cereb. Cortex 24, 859– 872.

112. Ioannidis, N.M., Rothstein, J.H., Pejaver, V., Middha, S., McDonnell, S.K., Baheti, S., Musolf, A., Li, Q., Holzinger, E., Karyadi, D., et al. (2016). REVEL: An Ensemble Method for Predicting the Pathogenicity of Rare Missense Variants. Am. J. Hum. Genet. 99, 877–885.

113. Samocha, K.E., Kosmicki, J.A., Karczewski, K.J., O’Donnell-Luria, A.H., Pierce-Hoffman, E., MacArthur, D.G., Neale, B.M., and Daly, M.J. (2017). Regional missense constraint improves variant deleteriousness prediction. bioRxiv, 148353. 10.1101/148353.

114. Samocha, K.E., Robinson, E.B., Sanders, S.J., Stevens, C., Sabo, A., McGrath, L.M., Kosmicki, J.A., Rehnström, K., Mallick, S., Kirby, A., et al. (2014). A framework for the interpretation of de novo mutation in human disease. Nat. Genet. 46, 944–950.

115. Karczewski, K.J., Solomonson, M., Chao, K.R., Goodrich, J.K., Tiao, G., Lu, W., Riley-Gillis, B.M., Tsai, E.A., Kim, H.I., Zheng, X., et al. (2022). Systematic single-variant and gene-based association testing of thousands of phenotypes in 394,841 UK Biobank exomes. Cell Genom. 2, 100168.

116. Kang, H.J., Kawasawa, Y.I., Cheng, F., Zhu, Y., Xu, X., Li, M., Sousa, A.M.M., Pletikos, M., Meyer, K.A., Sedmak, G., et al. (2011). Spatio-temporal transcriptome of the human brain. Nature 478, 483–489.

117. Lovett-Barron, M., Kaifosh, P., Kheirbek, M.A., Danielson, N., Zaremba, J.D., Reardon, T.R., Turi, G.F., Hen, R., Zemelman, B.V., and Losonczy, A. (2014). Dendritic inhibition in the hippocampus supports fear learning. Science 343, 857–863.

118. Acsády, L., Görcs, T.J., and Freund, T.F. (1996). Different populations of vasoactive intestinal polypeptide-immunoreactive interneurons are specialized to control pyramidal cells or interneurons in the hippocampus. Neuroscience 73, 317–334.

119. Acsády, L., Arabadzisz, D., and Freund, T.F. (1996). Correlated morphological and neurochemical features identify different subsets of vasoactive intestinal polypeptide-immunoreactive interneurons in rat hippocampus. Neuroscience 73, 299–315.

120. Tyan, L., Chamberland, S., Magnin, E., Camiré, O., Francavilla, R., David, L.S., Deisseroth, K., and Topolnik, L. (2014). Dendritic inhibition provided by interneuron-specific cells controls the firing rate and timing of the hippocampal feedback inhibitory circuitry. J. Neurosci. 34, 4534–4547.

121. Pelkey, K.A., Chittajallu, R., Craig, M.T., Tricoire, L., Wester, J.C., and Mcbain, X.C.J. (2017). Hippocampal GABAergic inhibitory interneurons. Physiol. Rev. 97, 1619–1747.

122. Francavilla, R., Villette, V., Luo, X., Chamberland, S., Muñoz-Pino, E., Camiré, O., Wagner, K., Kis, V., Somogyi, P., and Topolnik, L. (2018). Connectivity and network state-dependent recruitment of long-range VIP-GABAergic neurons in the mouse hippocampus. Nat. Commun. 9, 5043.

123. Darmanis, S., Sloan, S.A., Zhang, Y., Enge, M., Caneda, C., Shuer, L.M., Hayden Gephart, M.G., Barres, B.A., and Quake, S.R. (2015). A survey of human brain transcriptome diversity at the single cell level. Proc. Natl. Acad. Sci. U. S. A. 112, 7285–7290.

124. Scala, F., Kobak, D., Bernabucci, M., Bernaerts, Y., Cadwell, C.R., Castro, J.R., Hartmanis, L., Jiang, X., Laturnus, S., Miranda, E., et al. (2021). Phenotypic variation of transcriptomic cell types in mouse motor cortex. Nature 598, 144–150.

125. Gouwens, N.W., Sorensen, S.A., Baftizadeh, F., Budzillo, A., Lee, B.R., Jarsky, T., Alfiler, L., Baker, K., Barkan, E., Berry, K., et al. (2020). Integrated morphoelectric and transcriptomic classification of cortical GABAergic cells. Cell 183, 935–953.e19.

126. Korsunsky, I., Millard, N., Fan, J., Slowikowski, K., Zhang, F., Wei, K., Baglaenko, Y., Brenner, M., Loh, P.-R., and Raychaudhuri, S. (2019). Fast, sensitive and accurate integration of single-cell data with Harmony. Nat. Methods 16, 1289–1296.

127. Lopez, R., Regier, J., Cole, M.B., Jordan, M.I., and Yosef, N. (2018). Deep generative modeling for single-cell transcriptomics. Nat. Methods 15, 1053–1058.

128. Yao, Z., van Velthoven, C.T.J., Kunst, M., Zhang, M., McMillen, D., Lee, C., Jung, W., Goldy, J., Abdelhak, A., Aitken, M., et al. (2023). A high-resolution transcriptomic and spatial atlas of cell types in the whole mouse brain. Nature 624, 317–332.

129. Bakken, T.E., Jorstad, N.L., Hu, Q., Lake, B.B., Tian, W., Kalmbach, B.E., Crow, M., Hodge, R.D., Krienen, F.M., Sorensen, S.A., et al. (2021). Comparative cellular analysis of motor cortex in human, marmoset and mouse. Nature 598, 111–119.

130. Berg, J., Sorensen, S.A., Ting, J.T., Miller, J.A., Chartrand, T., Buchin, A., Bakken, T.E., Budzillo, A., Dee, N., Ding, S.-L., et al. (2021). Human neocortical expansion involves glutamatergic neuron diversification. Nature 598, 151–158.

131. Kaifosh, P., Zaremba, J.D., Danielson, N.B., and Losonczy, A. (2014). SIMA: Python software for analysis of dynamic fluorescence imaging data. Front. Neuroinform. 8, 80.

